# A Minimal Chemo-mechanical Markov Model for Rotary Catalysis of F_1_-ATPase

**DOI:** 10.1101/2025.06.26.661389

**Authors:** Yixin Chen, Helmut Grubmüller

## Abstract

F_1_-ATPase, the catalytic domain of ATP synthase, is pivotal for mechanochemical energy conversion in mitochondria. Aiming at a minimal yet quantitative and thermodynamically consistent model for its rotary catalysis mechanism, here we developed a chemo-mechanical Markov model incorporating essential conformational and chemical degrees of freedom. By systematically evaluating over 14,000 model variants via Bayesian inference and cross-validation, we find that a fully functional minimal model requires four functionally distinct **β**-subunit conformations. Our model reconciles the decade-long bi-site versus tri-site controversy, showing that both pathways contribute depending on ATP concentration. Furthermore, our model suggests a Brownian-ratchet-like mechanism that explains the observation that one ATP hydrolysis event can trigger larger than 120° rotations, thereby explaining seemingly over 100% efficiency. Beyond this prototypic example of a complex biomolecular machine, our approach should enable one to study other enzymatic mechanisms that implement close coupling between conformational motions, substrate binding, and chemical reactions.

## Introduction

The F_1_F_O_-ATP synthase (ATP synthase) is a highly conserved biological rotary motor that synthesizes ATP from ADP and inorganic phosphate (Pi), using energy derived from the proton concentration gradient across membranes [1, 2, 3]. The enzyme comprises two rotary motors, the membrane-embedded F_O_ motor and the soluble F_1_-ATPase, connected by a rotating central stalk (Fig. 1a). In vivo, ATP synthase typically operates in the synthesis mode. Here, proton flow through the F_O_ motor drives the clockwise rotation of the central stalk, which in turn drives ATP synthesis within the three catalytic sites of the F_1_-ATPase domain. Here we focus on the minimal α_3_β_3_γ structure of F_1_-ATPase (Fig. 1b). This core structure consists of a stator ring formed by three α- and three β-subunits (gray and purple, respectively), with the γ-subunit (green) located centrally as an axle, forming part of the central stalk [4, 5]. Each β-subunit forms a catalytic site at an α-β interface. Remarkably, F_1_-ATPase can reach near 100% efficiency [6, 7, 8, 9, 10]. Accordingly, the F_1_-ATPase can also operate in reverse (the hydrolysis) mode, where ATP hydrolysis within the catalytic sites drives rotation of the γ-subunit [11, 12, 13, 14, 8, 15].

**Figure 1.**
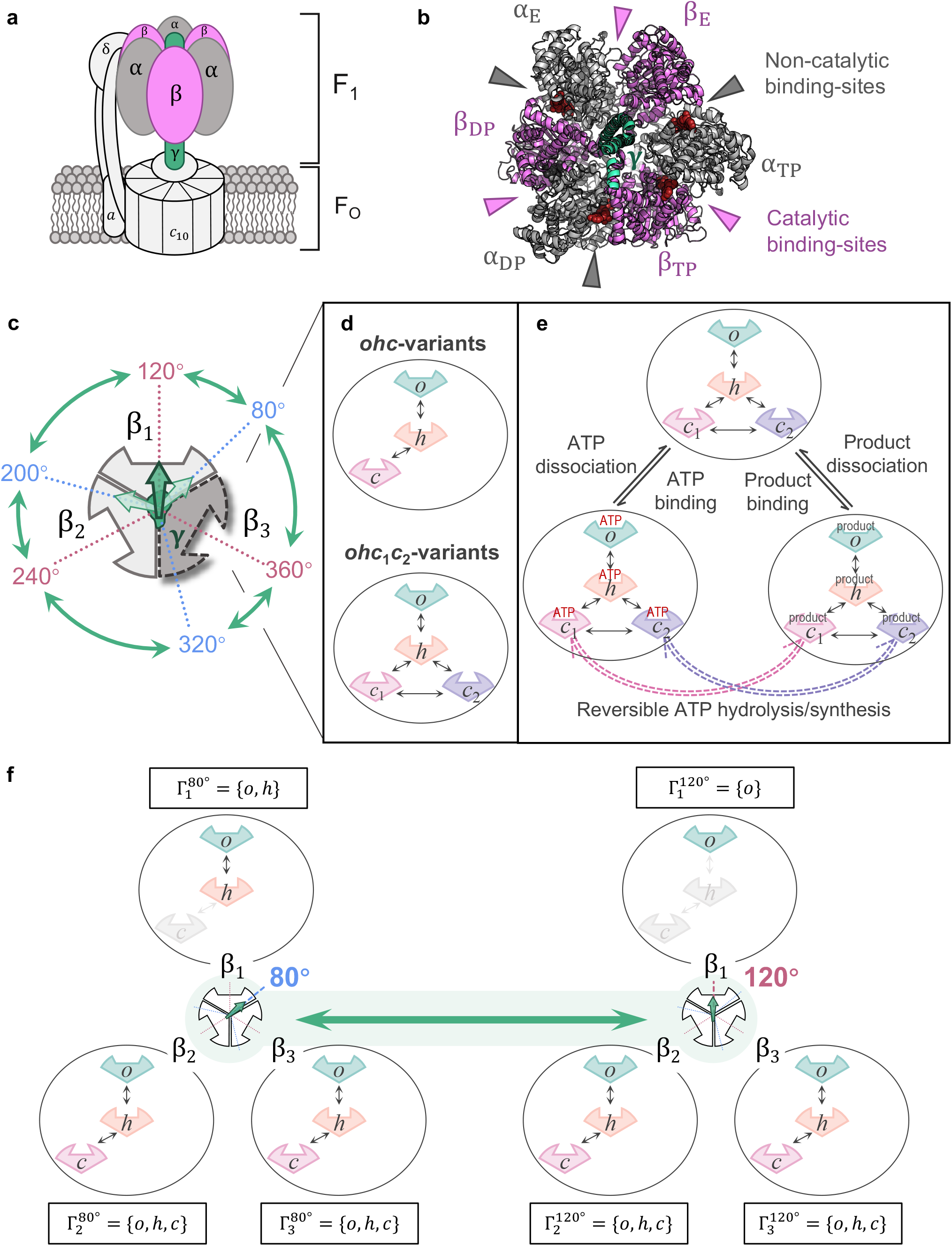
The chemo-mechanical Markov model of F_1_-ATPase. **a** Schematic of the F_1_F_O_-ATP synthase, showing the membrane-embedded F_O_ sector and the soluble F_1_ catalytic sector (F_1_-ATPase). **b** Top-down view of the α_3_β_3_γ structure of F_1_-ATPase (PDB: 1BMF) [4], highlighting the three catalytic sites (purple arrows) and three non-catalytic sites (gray arrows) at the α-β interfaces. **c**–**e** Illustration of the DOFs defining the Markov states (Table 1). **c** The orientation of the central γ-subunit is discretized into six states corresponding to the catalytic (e.g., 80°) and ATP-waiting (e.g., 120°) dwells observed in single-molecule experiments [8, 15]. The γ-subunit rotates in discrete substeps between every two adjacent orientations. **D** Each β-subunit converts between three ({*o, h, c*} for *ohc*-variants) or four ({*o, h, c*_1_, *c*_2_} for *ohc*_1_*c*_2_-variants) distinct conformations. Direct transitions between open (*o*) and closed (*c, c*_1_, *c*_2_) conformations are disallowed, positioning the half-closed (*h*) conformation as a mandatory intermediate. **e** Each β-subunit converts between three binding states: apo (E), ATP-bound (T) and product-bound (D). Reversible ATP hydrolysis/synthesis (dashed arrows) occurs only when the β-subunit is in a catalytically active conformation. **f** Visualization of the γ-β restrictions (the hyperparameter 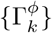) that define a specific model variant. This visualized example corresponds to the *ohc*-*w* variant (Table 2). The set of accessible conformations (e.g., {*o*}, {*o, h*}, or {*o, h, c*}) is illustrated for each β-subunit at 80° (left) and 120° (right). In this example, at 120°, the γ-subunit sterically forces β_1_ into the *o* conformation, while β_2_ and β_3_ are unrestricted. These restrictions demonstrate how the asymmetric γ-subunit results in different conformational ensembles of the three chemically identical β-subunits.

To rationalize this rotary catalysis, Paul Boyer proposed the initial binding change model [16, 17, 18], in which rotation of the γ-subunit induces alternating conformational changes within the three β-subunits. These different conformations exhibit different binding affinities for ATP and ADP [19, 20, 21, 22], thereby driving major catalytic steps such as substrate ATP binding, bound ATP hydrolysis, and product ADP/Pi dissociation. The tight coupling between these conformational and chemical degrees of freedom (DOFs) is at the core of this model. Recognized as a universal principle underlying rotary catalysis, this tight coupling mechanism was later generalized to the closely related V- and A-ATPases, grounded in the evolutionary, structural, and functional homology between the F-ATPase subunits (α, β, γ) and their V/A-ATPase counterparts (B, A, DF) [23, 24, 25, 26, 27, 28, 29, 30].

Over time, the original binding change model [17] has been repeatedly modified to accommodate new experimental observations, leading to a family of related models [20, 31, 22, 32, 5, 8, 33, 34, 35]. However, despite decades of intensive research and a wealth of experimental data, there is yet no consensus on the precise mechanism of F_1_-ATPase. Particularly, several questions remain unresolved, reflected in the differences among these models: (i) how many functionally distinct conformational states of the β-subunits are required for F_1_-ATPase function (three [16, 20], four [31, 22, 32], or potentially more [5, 36]); (ii) how many catalytic sites are simultaneously active during catalysis (bi-site [16, 6, 8, 37, 38, 39] versus tri-site [40, 20, 21, 22, 15, 34, 32] mechanisms); (iii) how are the static structural snapshots from cryoEM and X-ray crystallography correlated with the functional dwells in F_1_-ATPase catalytic cycle.

Methodologically, these earlier models typically focused on very few pre-assumed chemo-mechanical states and catalytic pathways, often motivated by biochemical intuition or static structures. This focus has made it challenging to achieve a fully quantitative and thermodynamically consistent description for the rather complex F_1_-ATPase system. Complementing these models, highly coarse-grained continuum ratchet models [41, 42, 43] have been analyzed, and coarse-grained [36], atomistic [44, 45, 46, 47, 48, 49], and QM/MM simulations [50, 51] have been performed. While these studies have revealed valuable mechanistic insights into rotary catalysis, they have not yet provided a complete thermodynamic description of the full catalytic cycle. Ultimately, a coherent framework integrating the transition rates with the free energies for all possible combinations of the DOFs has not been established.

More fundamentally, the classical binding change models typically provide a macroscopic, ensemble-average view of rotary catalysis, and do not explicitly incorporate the inherent stochasticity at the molecular level. This perspective tends to intuitively align with a power-stroke mechanism for the chemo-mechanical energy transduction, assuming ATP hydrolysis directly generates the torque needed for uni-directional rotation. Nevertheless, extensive single-molecule experiments demonstrate large rotational fluctuations of the central stalk, pointing to a highly stochastic energy transduction process [13, 14, 8, 52, 15, 23, 53]. Accordingly, a Brownian ratchet mechanism has been proposed: instead of directly exerting torque on the central stalk, ATP hydrolysis induces affinity changes in the catalytic subunits that directionally rectify these random thermal fluctuations [30, 29]. How to quantitatively reconcile the classical binding change paradigm with the stochastic nature of these microscopic molecular machines remains an open question [8, 52, 15, 29, 30].

In this work, we construct a minimal chemo-mechanical Markov model that unifies the conformational and chemical DOFs governing F_1_-ATPase rotary catalysis to quantitatively capture its inherent stochasticity. To simultaneously explain a heterogeneous set of experimental data – including measured titration curves [21], near 100% chemo-mechanical coupling efficiency [8], and the consensus chemo-mechanical coupling scheme [8, 54, 33, 15, 55, 56, 57] – we systematically evaluated various model variants via extensive Bayesian parameter inference and cross-validation. This systematic model comparison reveals that a fully functional model requires at least four, rather than three, distinct β-subunit conformations, contrary to previous assumptions [16, 20]. Furthermore, we find that a nucleotide-bound β-subunit energetically favors the closed conformation, thus resolving a previous controversy [58, 46, 48]. Remarkably, our model also reconciles the bi-site versus tri-site controversy by showing that both pathways contribute depending on ATP concentration. By construction, our model provides a quantitative understanding of the thermodynamics, kinetics, and inherent stochasticity of F_1_-ATPase function.

## Results

### Construction of a chemo-mechanical Markov model

To establish our chemo-mechanical Markov model, we specifically focus on the well-characterized F_1_-ATPases from *E. coli* and *Bacillus* PS3 (EF_1_, TF_1_), because the mechanistic details of F_1_-ATPases from other species may differ, e.g., in their sub-stepping behaviors [59, 60, 8, 15, 61, 62, 63]. Hereafter, unless otherwise specified, F_1_-ATPase refers to EF_1_ and TF_1_. Below, we outline the conceptual framework of this model, while the rigorous mathematical formulations are provided in Methods.

In this model, a unique chemo-mechanical state of F_1_-ATPase (a Markov state) is specified by the orientation of the γ-subunit, and the conformational and binding states of the three β-subunits (Equation (1)), totaling seven DOFs listed in Table 1.

**Table 1.** Degrees of freedom (DOFs) of our chemo-mechanical Markov model. The table lists the possible values of the seven DOFs that specify a Markov state (Equation (1)) and their experimental justification.

| DOFs | Possible values | Experimental justification |
| --- | --- | --- |
| Orientation of the γ-subunit ($\phi$) | $\phi = 80^\circ, 200^\circ, 320^\circ$, (catalytic dwells)<br>$120^\circ, 240^\circ, 360^\circ$ (ATP-waiting dwells) | Single-molecule experiments [8, 15, 59] |
| Conformational states of the β-subunits ($\mathcal{C}^{(k)}$, $k = 1, 2, 3$) | $\mathcal{C}^{(k)} \in \{o, h, c\}$ (*ohc*-variants)<br>or $\mathcal{C}^{(k)} \in \{o, h, c_1, c_2\}$ (*ohc<sub>1</sub>c<sub>2</sub>*-variants) | Observation of several β-subunit conformations in crystal and cryoEM structures ($\beta_E$, $\beta_{TP}/\beta_{DP}$, $\beta_{HC}/\beta_{HO}$) [4, 5, 32, 65, 64] |
| Binding states of the β-subunits ($\mathcal{B}^{(k)}$, $k = 1, 2, 3$) | $\mathcal{B}^{(k)} = E$ (empty),<br>T (ATP-bound),<br>D (product-bound) | $\beta_E$, $\beta_{TP}/\beta_{DP}$, $\beta_{HC}/\beta_{HO}$ observed in different binding states [4, 5, 32, 65, 64] |

First, we consider the rotation of the γ-subunit. For EF_1_ and TF_1_, this rotation is known to proceed via well-characterized 80° and 40° substeps [8, 54, 15, 59], resulting in the six different γ-subunit orientations in our model (Fig. 1c).

Second, we consider for each β-subunit transitions between several conformational states (Fig. 1d). Crystal and cryoEM structures of F_1_-ATPase reveal three clusters of clearly distinct conformations, often termed open (β_E_), half-open/half-closed (β_HO_/β_HC_), and closed (β_TP_/β_DP_) [4, 5, 32, 64, 65]. Accordingly, we include three distinct conformations in our model, denoted as *o, h*, and *c* respectively. While tentatively associated with the experimentally resolved structures, these conformations are defined in our model by their functional roles: (1) the *o* conformation embeds a widely open catalytic site of very low nucleotide binding affinity that awaits substrate ATP binding; (2) the *c* conformation is the catalytically active state where ATP hydrolysis and synthesis occur reversibly and almost in equilibrium (*K*_eq_ ≈ 1 [6, 66, 67, 68]); (3) the *h* conformation is an intermediate state between the *o* and *c* conformations.

It is proposed that the two closed conformations, β_TP_ and β_DP_, despite high structural similarity, are also functionally distinct, due to fine details at the catalytic interface between adjacent α- and β-subunits, e.g., involving the “arginine finger” [4, 5, 69, 67, 70, 32]. To systematically address the minimal number of functionally distinct conformations required for F_1_-ATPase function, we tested two hypotheses – three-conformation versus four-conformation – by constructing two major classes of model variants (Fig. 1d): the *ohc*- and *ohc*_1_*c*_2_-variants, assuming one (*c*) or two functionally distinct closed conformations (*c*_1_, *c*_2_). Further, for the ohc_1_c_2_-variants, because it is controversially debated whether or not both c_1_ and *c*_2_ are catalytically active [5, 71, 72], we tested both hypotheses by investigating 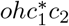 -variants and 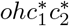 -variants, where the asterisks (*) denote the catalytically active conformations.

Third, we consider for each β-subunit conversion between three binding states (Fig. 1e) [4, 5, 1, 32, 65, 64]: an apo state (E), an ATP-bound state (T), and a single product-bound state (D) that merges the ADP-, Pi-, and ADP+Pi-bound states. This merged D state, a simplification also adopted in a recent theoretical analysis [73], results in an effective description of the kinetics of two chemical dissociation events, namely the dissociation of ADP and of Pi (further explained below).

As a key conceptual step, we a priori include all combinatorially possible combinations of the seven DOFs, resulting in 6 × (3 × 3)^3^*/*3 = 1458 (*ohc*-variants) or 6 × (4 × 3)^3^*/*3 = 3456 (*ohc*_1_*c*_2_-variants) Markov states in our model; here, the counts already take into account the three-fold rotational symmetry of F_1_-ATPase, hence the division by three. Further, we restrict the transitions between these 1458 (or 3456) Markov states to those involving the change of only one DOF at a time, and, crucially, assumed all these transitions to be reversible. These elementary transitions, as illustrated in Fig. 1c–e, include stepwise rotations of the γ-subunit in both directions, conformational transitions of each individual β-subunit between the three or four conformations, binding and dissociation of ATP or product, and reversible ATP hydrolysis/synthesis in a catalytically active β-subunit.

Notably, our definition of elementary transitions inherently decouples conformational changes from chemical events like binding/dissociation or hydrolysis/synthesis. Without losing generality, such formal decoupling facilitates easier modeling of complex, multi-step catalytic processes as sequences of elementary events. For example, we describe product release in our model as a sequence of conformational changes (opening of the catalytic site) and subsequent chemical dissociation of the products. The fact that the former must precede the latter is described in our model by a very low product dissociation rate from a closed catalytic site (Supplementary Equation (5) and Supplementary Table 1). In principle, ADP- and Pi-release could be described separately and similarly; however, and aiming at a minimal yet kinetically sufficiently accurate model, we decided to merge the ADP-, Pi-, and ADP+Pi-bound states into a single product-bound state D [73]. In this approximation, the rates for dissociation of the two products are combined into an effective rate, governed by the slower of these two individual rates. This approximation is supported by experiments suggesting that only one of these two dissociation events is indeed rate-limiting [15, 55, 56, 57, 74].

By including all possible Markov states and reversible elementary transitions (Fig. 1c-e), our model explores all potential catalytic pathways without a priori bias, in contrast to previous binding change models [16, 20, 31, 22, 32, 5, 8, 33, 34, 35]. To quantitatively describe the dynamics of this Markov model (Equation (8)), a transition rate needs to be specified for each elementary transition (Supplementary Table 2). Following transition state theory (Equation (5)), the rate coefficient is determined by the free energy barrier between the two connected states; further, as required by the principle of local detailed balance, the rate coefficient of the backward transition is related to the forward rate coefficient by the difference between the free energies of the two states (Equation (6)). However, to compare our model against experimental data characterizing steady-state properties of F_1_-ATPase, we evaluated relevant model predictions under non-equilibrium steady-state (NESS) conditions (Methods), mimicking typical experimental setups where F_1_-ATPase operates in contact with large external reservoirs maintaining constant ATP and ADP concentrations [19, 21, 8]. In NESS, while local detailed balance is preserved for every individual elementary transition, the sustained chemical potential difference between ATP and ADP (Equation (7)) breaks the global detailed balance of the system, driving a net flux through the network of states that manifests as observable ATP turnover and rotation.

We next decomposed the state free energies that dictate the transition rates into several contributions (Equation (2)). Conformational free energies and binding free energies of individual β-subunits are included, characterizing the thermodynamic properties of the several functionally distinct conformations. Note that these energy terms also implicitly account for local interactions between adjacent α- and β-subunits at the catalytic interface [4, 5, 69, 67, 70, 32]. Because single-molecule and structural studies [1, 75, 2, 4, 9, 70] clearly point to a dominant role of the direct interaction between the γ- and β-subunits in governing F_1_-ATPase rotary catalysis, we decided to include these γ-β interactions, but not the long-range allosteric interactions across the α_3_β_3_ stator ring. Whereas these allosteric interactions are proposed to facilitate residual activity of γ-depleted and γ-truncated constructs of F_1_-ATPase [76, 77, 78, 79], they are much weaker than the γ-β interactions, as evidenced by the greatly reduced turnover rates and coupling efficiency of the γ-depleted/truncated constructs [76, 77]. Along similar lines, and again aiming at a minimal model, we also refrained from including separate energy terms for local α-β interactions.

For defining the γ-β interaction energies (Equation (4)), we note that structural [4, 5, 64, 32, 65] and theoretical studies [48, 36] have suggested steric repulsions between the γ- and β-subunits: the γ-subunit pushes the N-terminal domain of the β-subunit that faces its bulge side outwards, forcing this β-subunit to adopt an open or partially open conformation. Accordingly, we defined an infinitely high interaction energy for some of the non-open conformation(s) of a β-subunit at a particular γ-orientation, which precludes these conformations from being populated. However, precisely which combinations of γ-orientations and conformations should be subjected to such restrictions is not immediately obvious from previous studies. Therefore, we introduced γ-β restrictions – an ensemble of accessible conformations defined for each β-subunit at each γ-orientation – as a key hyperparameter defining specific model variants (graphically illustrated in Fig. 1f, and formally defined in Methods). Even with symmetry considerations and plausible constraints, this hyperparameter leads to a large number of possibilities, e.g., more than 600 *ohc*-variants (Methods). To resolve this ambiguity, we first investigated several plausible *ohc*-variants, followed by a more systematic model exclusion analysis, as detailed below.

The above definitions of transition rates and state free energies have introduced ∼20 parameters into our model, including conformational free energies and binding free energies assigned to each β-subunit conformation, as well as several free energy barriers (Methods).

### No *ohc*-variant explains all available experimental data

To rigorously evaluate competing mechanistic hypotheses, we first tested whether a minimal model including only three functionally distinct β-subunit conformations *o, h*, and *c* (three-conformation hypothesis) could suffice to explain the available experimental data. Specifically, we examined three plausible *ohc*-variants (Table 2), asking if any of them, with optimized parameters, could simultaneously reproduce three key macroscopic properties defining F_1_-ATPase rotary-catalysis function (training data, see Methods): (1) the turnover rates *k*_cat_ [19], (2) 100% chemo-mechanical coupling efficiency [8, 54, 6, 80], and (3) average catalytic site occupancies under varying ATP/ADP concentrations (nucleotide titration curves) [21]. We employed a Bayesian approach (Methods) to broadly sample the parameter space and obtain a large ensemble of parameter sets covering potentially multimodal high-posterior regions.

**Table 2.**
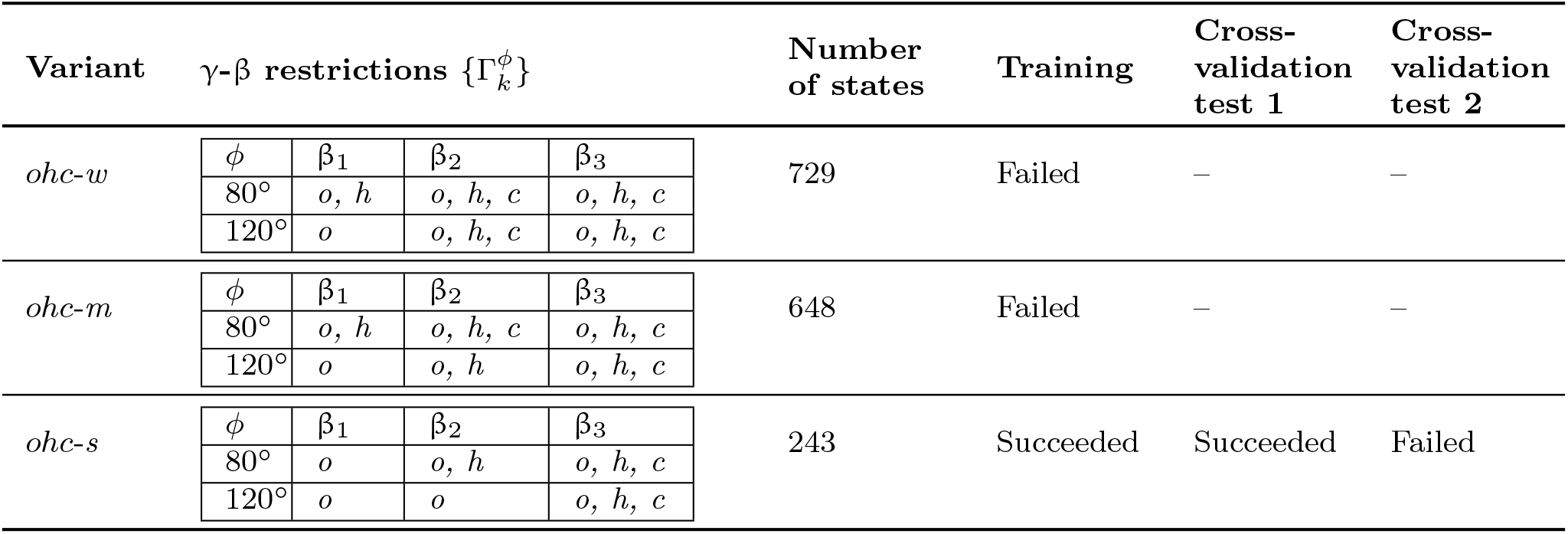
Definition and evaluation of three plausible *ohc*-variants. For each variant, the table lists its defining γ-β restrictions, the number of resulting Markov states, and the outcome of the training and cross-validation procedures (Methods). Training: A Bayesian approach was used to find parameter sets that reproduce the training data. Cross-validation: The trained parameter sets were tested against two sets of validation data, first (cross-validation test 1) the observed population shift between dwells [8] and second (cross-validation test 2) the consensus chemo-mechanical coupling scheme (Table 6 and Supplementary Fig. 2). “Succeeded” or “Failed” indicates whether parameter sets consistent with the respective data were found.

Only the *ohc*-*s* variant yielded ∼9000 parameter sets of similarly high posterior probabilities that successfully reproduced the training data within experimental uncertainty (Fig. 2a&b). Notably, some predicted chemo-mechanical coupling efficiencies exceed 100% (Fig. 2a). This does not violate energy conservation, because this efficiency is defined as a ratio of rates (Equation (12)) rather than energies, consistent with previous experimental observations [8] (mechanistic explanation provided in Discussion).

**Figure 2.**
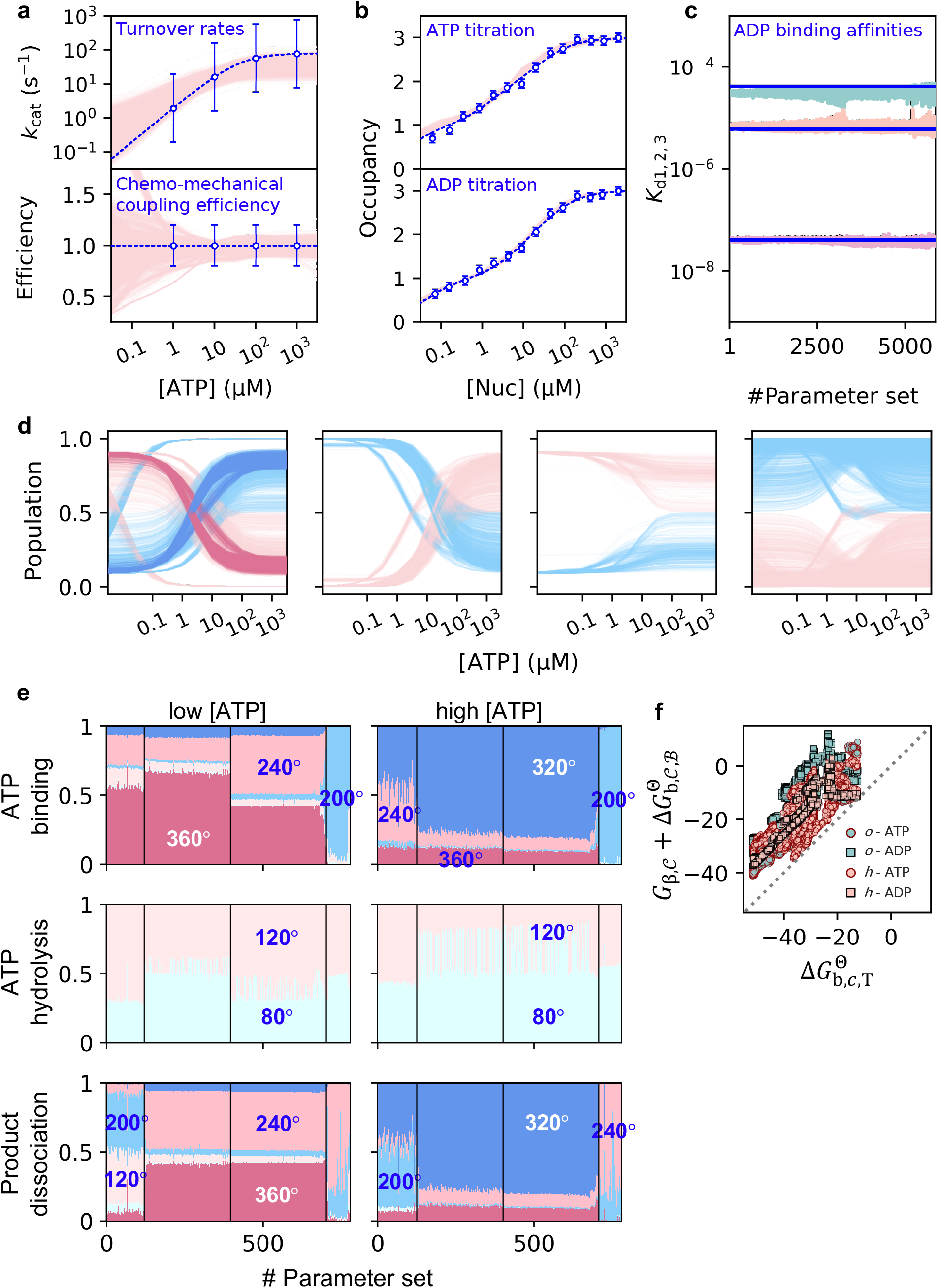
Training and cross-validation of the *ohc*-*s* variant. **a, b** Model predictions from the ensemble of trained parameter sets (pink curves) versus training data (blue circles), including turnover rates, chemo-mechanical coupling efficiency, and nucleotide titration curves. Vertical bars indicate the standard deviations of the likelihood functions (normal or log-normal distributions) adopted in the Bayesian training approach, as detailed in Supplementary Table 7. **c** Predicted apparent ADP binding affinities (green, orange and pink curves) versus the experimentally determined values (blue horizontal bars). **d** First cross-validation test: predicted populations of the 80°-(blue) and 120°-states (pink) versus ATP concentration ([ATP]). The trained parameter sets are classified into four groups based on their qualitative predictions. Only the leftmost group correctly reproduces the experimentally observed population shift from 120°-dwells at low [ATP] to 80°-dwells at high [ATP]. 750 parameter sets are selected from this successful group in (highlighted in darker blue and darker pink) and subjected to the second cross-validation test. **e** Second cross-validation test: predicted probabilities of the major catalytic steps (ATP binding, hydrolysis, product dissociation) at different γ-orientations for low (0.1 µM) and high (1 mM) [ATP] are represented by the height of the differently colored areas in each panel. The selected 750 parameter sets are further classified into four mechanistically distinct clusters (separated by vertical bars). **f** A common property of all trained parameter sets: the free energy of a nucleotide-bound β-subunit is lowest when it adopts the catalytically active, closed (*c*) conformation. The plot compares the total free energies of an ATP/product-bound β-subunits in the *o* or *h* conformations (y-axis) against that of an ATP-bound β-subunit in the *c* conformation (x-axis). Source data are provided as a Source Data file.

To quantitatively assess agreement with experimental ADP titration curves [21], we derived apparent ADP binding affinities by fitting the curves (Fig. 2b) to an equilibrium binding model (Equation (17), see Methods). For the *ohc*-*s* variant (Fig. 2c), the predicted apparent affinities (pink, orange and green curves) agree well with the experimental values (horizontal lines). In contrast, the *ohc*-*w* and *ohc*-*m* variants disagree with the experimental values (Supplementary Fig. 1), and were therefore excluded from further analysis.

Training the *ohc*-*s* variant provided two key insights. First, despite reproducing the training data equally well, the ∼9000 trained parameter sets predicted several distinct mechanistic scenarios, highlighting the need for additional experimental data to constrain our model. Second, and remarkably, across all scenarios, the catalytically active, closed conformation *c* is the state of lowest free energy for a nucleotide-bound β-subunit (Fig. 2f). Thus, unless sterically restricted by the γ-subunit, a nucleotide-bound β-subunit always remains closed. This energetic property is likely a necessary condition for our model to achieve near 100% efficiencies (Discussion).

To identify which mechanistic scenario is consistent with experimental observations, we cross-validated the model against experimental data that characterize mechanistic details of F_1_-ATPase at a more microscopic level (validation data, see Methods).

The first cross-validation test examined the population shift between the catalytic (γ-orentation ∼80°) and ATP-waiting dwells (∼120°). Experimentally, the dominant state shifts from ATP-waiting to catalytic dwells as ATP concentration ([ATP]) increases [8]. The trained parameter sets split into four groups predicting different trends (Fig. 2d). Only the first group correctly showed 120°-states and 80°-states dominating at low and high [ATP], respectively (leftmost panel); the other three groups were therefore discarded.

The second cross-validation test evaluated whether the 750 parameter sets selected from the first group (darker pink and blue curves) reproduce the γ-orientations of ATP binding, hydrolysis, and product dissociation steps, as defined by a consensus chemomechanical coupling scheme (see Methods and Supplementary Fig. 2; synthesized from a broad range of experimental studies [8, 54, 33, 15, 57, 34, 70, 20, 21, 22, 32, 65, 56, 55, 67, 66, 75] as reviewed in Supplementary Note 2). While this consensus scheme adopts a deterministic view where each catalytic step is assigned to a specific γ-orientation, our model inherently predicts stochastic probability distributions for these steps to occur at different γ-orientations (Methods, see particularly Equation (13)). Therefore, we tested whether these probabilities concentrated at the consensus γ-orientations.

Based on the predicted probabilities (Fig. 2e, represented by the height of differently colored areas), we classified these parameter sets into four mechanistically distinct clusters (separated by vertical bars). However, none aligns with the consensus coupling scheme (disagreements summarized in Supplementary Table 3), for instance by predicting ATP hydrolysis at γ-orientations earlier than observed. Thus, the *ohc*-*s* variant ultimately fails this cross-validation test.

In summary, despite their large parameter spaces (14 free parameters), all three tested *ohc*-variants failed to explain all available experimental data. If true, this finding would suggest that the three-conformation hypothesis is fundamentally incompatible with the data, at least for these tested γ-β restrictions. However, before finally rejecting this hypothesis, we had to conclusively rule out two alternaive explanations for this failure: insufficient parameter sampling and incorrect γ-β restrictions. We excluded the former because independent optimization runs from random starting points consistently converged to the same scenarios (Fig. 2e). The latter is addressed below, by systematically examining whether a viable choice was hidden among the other ∼600 untested *ohc*-variants.

### The three-conformation hypothesis is fundamentally incompatible with experimental data

Rather than exhaustively test the remaining *ohc*-variants via the computationally expensive training + cross-validation procedures, we first derived several physical constraints on the γ-β restrictions from the consensus coupling scheme to systematically reduce the number of candidates. These constraints are summarized in Supplementary Table 4 (see Methods and Supplementary Note 3 for detailed explanations). Briefly, this analysis leverages the fundamental binding-change principle [16, 17, 6, 15, 35, 68, 73]: γ-subunit rotation induces sequential β-subunit conformational changes, altering their nucleotide binding affinities to drive the catalytic steps. Complemented by our earlier finding that a nucleotide-bound β-subunit energetically favors the closed conformation unless sterically forced open, this principle implies that γ-subunit rotation immediately following a catalytic step must modulate the accessible conformational ensemble of the β-subunit to induce the required conformational change.

Specifically, the constraint governing product dissociation at +320° relative to the ATP binding angle differentiates two families of model variants (Supplementary Equation (7) in Supplementary Table 4): Family A forces the β-subunit to fully open (*h* → *o*) during the +320° to +360° rotation, while Family B allows it to partially close (*o* → *h*). Thermodynamically, Family A implies a decrease in product binding affinity upon this rotation, whereas Family B implies an increase (Supplementary Equations (41),(42)). We excluded all Family B variants because an affinity increase contradicts the physical intuition that product dissociation is driven by reduced affinity [6, 8, 15], also supported by recent titration experiments observing a low-affinity site at 120°-dwells [22]. While Family B aligns with recent cryoEM structures [32], this apparent discrepancy is resolved by re-interpreting the functional roles of these structures (Discussion).

Together with an additional plausible constraint for model parsimony (Condition *monotonic*, see Supplementary Table 5), these conditions (Supplementary Table 4) reduced the number of candidate *ohc*-variants to just three (Supplementary Table 8). However, all three were further excluded due to inconsistencies with other experimental data (Supplementary Table 8). Particularly, one of these is the previously-tested *ohc*-*m* variant, which disagrees with the experimental apparent ADP binding affinities.

This finding raises a deeper question: physicochemically, why are variants like *ohc*-*m* and *ohc*-*w* incompatible with the experimental apparent affinities? The key insight comes from recognizing that these apparent affinities are not properties of single, static conformations in our model. Instead, analytical derivations based on equilibrium approximation show that each apparent affinity emerges as a superposition – a Boltzmann-weighted ensemble average – of the microscopic binding affinities of all accessible conformations dictated by the γ-β restrictions (Methods, see particularly Equation (19)). Thus, for our model to reproduce the vastly different experimental apparent affinities [21], the γ-β restrictions should impose different conformational ensembles onto the three β-subunits. Nevertheless, the *ohc*-*w* variant has identical γ-β restrictions for two β-subunits, inherently leading to two similar apparent affinities (Supplementary Fig. 1c) that contradict the experimental values. Similarly, although mathematically more complex, the *ohc*-*m* variant is also incompatible with the experimental apparent affinities (Supplementary Note 5).

Taken together, this systematic analysis considering physical-chemical constraints demonstrates that no *ohc*-variant can simultaneously explain all available experimental data, strongly suggesting that the underlying three-conformation hypothesis is insufficient.

### Several *ohc*_1_*c*_2_-variants are found to reproduce all available experimental data

We were therefore prompted to consider the next simplest extension of our model: including a fourth β-subunit conformation, resulting in *ohc*_1_*c*_2_-variants featuring two distinct closed states, *c*_1_ and *c*_2_ (Fig. 1d), tentatively associated with the experimentally observed β_TP_ and β_DP_ structures [4, 32, 65]. While the combinatorial possibilities of γ-β restrictions initially yield over 14,000 *ohc*_1_*c*_2_-variants (Methods), we systematically reduced this number to fifteen candidates (Supplementary Note 4), applying the constraints derived from the consensus chemo-mechanical coupling scheme (Supplementary Table 4), supplemented by a few additional constraints based on observed structures and model parsimony (Supplementary Table 5).

As the five simplest and most plausible candidates, we selected three 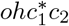 - and two 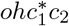 -variants, and applied the training + cross-validation procedures (Methods) to conclusively determine if any *ohc*_1_*c*_2_-variant could explain all available data (Table 3). All five candidates successfully reproduced both the training and initial validation datasets (Supplementary Figs. 3–5). Interestingly, these five variants predicted two distinct mechanistic scenarios regarding the γ-orientations of the major catalytic steps. The 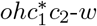 variant predicted these γ-orientations independent of [ATP] (Scenario 1; Fig. 3a), whereas the other four variants predicted ATP-dependent γ-orientations (Scenario 2; e.g., Fig. 3b, Supplementary Fig. 5b–d). For instance, in Scenario 2, product dissociation shifts to an earlier orientation (+240°) at low [ATP], while ATP binding occurs immediately after product dissociation (+320°) at high [ATP]. Both scenarios, however, remain compatible with the consensus chemo-mechanical coupling scheme.

**Table 3.** Definition and evaluation of the five most plausible *ohc*_1_*c*_2_-variants. For each variant, the table lists its defining γ-β restrictions, the number of resulting Markov states, and the outcome of the training and cross-validation procedures (Methods). Training: A Bayesian approach was used to find parameter sets that reproduce the training data. Cross-validation: The trained parameter sets were tested against three sets of validation data, first (cross-validation test 1) the observed population shift between dwells, second (cross-validation test 2) the consensus chemo-mechanical coupling scheme (Table 6 and Supplementary Fig. 2), and third (cross-validation test 2) an additional test against the measured ADP:ATP occupancy ratio. “Succeeded” or “Failed” indicates whether parameter sets consistent with the respective data were found.

| Variant | γ-β restrictions $\{\Gamma_k^\phi\}$ | | | | Number of states | Training + Cross-validation tests 1, 2 | Cross-validation test 3 |
| --- | --- | --- | --- | --- | --- | --- | --- |
| $ohc_1^*c_2-w$ | $\phi$ | $\beta_1$ | $\beta_2$ | $\beta_3$ | 972 | Succeeded | Succeeded |
| | $80^\circ$ | $o, h$ | $o, h, c_1, c_2$ | $o, h, c_2$ | | | |
| | $120^\circ$ | $o$ | $o, h, c_2$ | $o, h, c_1, c_2$ | | | |
| $ohc_1^*c_2-m.1$ | $\phi$ | $\beta_1$ | $\beta_2$ | $\beta_3$ | 864 | Succeeded | Failed |
| | $80^\circ$ | $o, h$ | $o, h, c_1, c_2$ | $o, h, c_2$ | | | |
| | $120^\circ$ | $o$ | $o, h$ | $o, h, c_1, c_2$ | | | |
| $ohc_1^*c_2^*-m.1$ | $\phi$ | $\beta_1$ | $\beta_2$ | $\beta_3$ | 864 | Succeeded | Succeeded |
| | $80^\circ$ | $o, h$ | $o, h, c_1, c_2$ | $o, h, c_2$ | | | |
| | $120^\circ$ | $o$ | $o, h$ | $o, h, c_1, c_2$ | | | |
| $ohc_1^*c_2-m.2$ | $\phi$ | $\beta_1$ | $\beta_2$ | $\beta_3$ | 864 | Succeeded | Succeeded |
| | $80^\circ$ | $o, h$ | $o, h, c_1$ | $o, h, c_1, c_2$ | | | |
| | $120^\circ$ | $o$ | $o, h$ | $o, h, c_1, c_2$ | | | |
| $ohc_1^*c_2^*-m.2$ | $\phi$ | $\beta_1$ | $\beta_2$ | $\beta_3$ | 864 | Succeeded | Succeeded |
| | $80^\circ$ | $o, h$ | $o, h, c_1$ | $o, h, c_1, c_2$ | | | |
| | $120^\circ$ | $o$ | $o, h$ | $o, h, c_1, c_2$ | | | |

**Figure 3.**
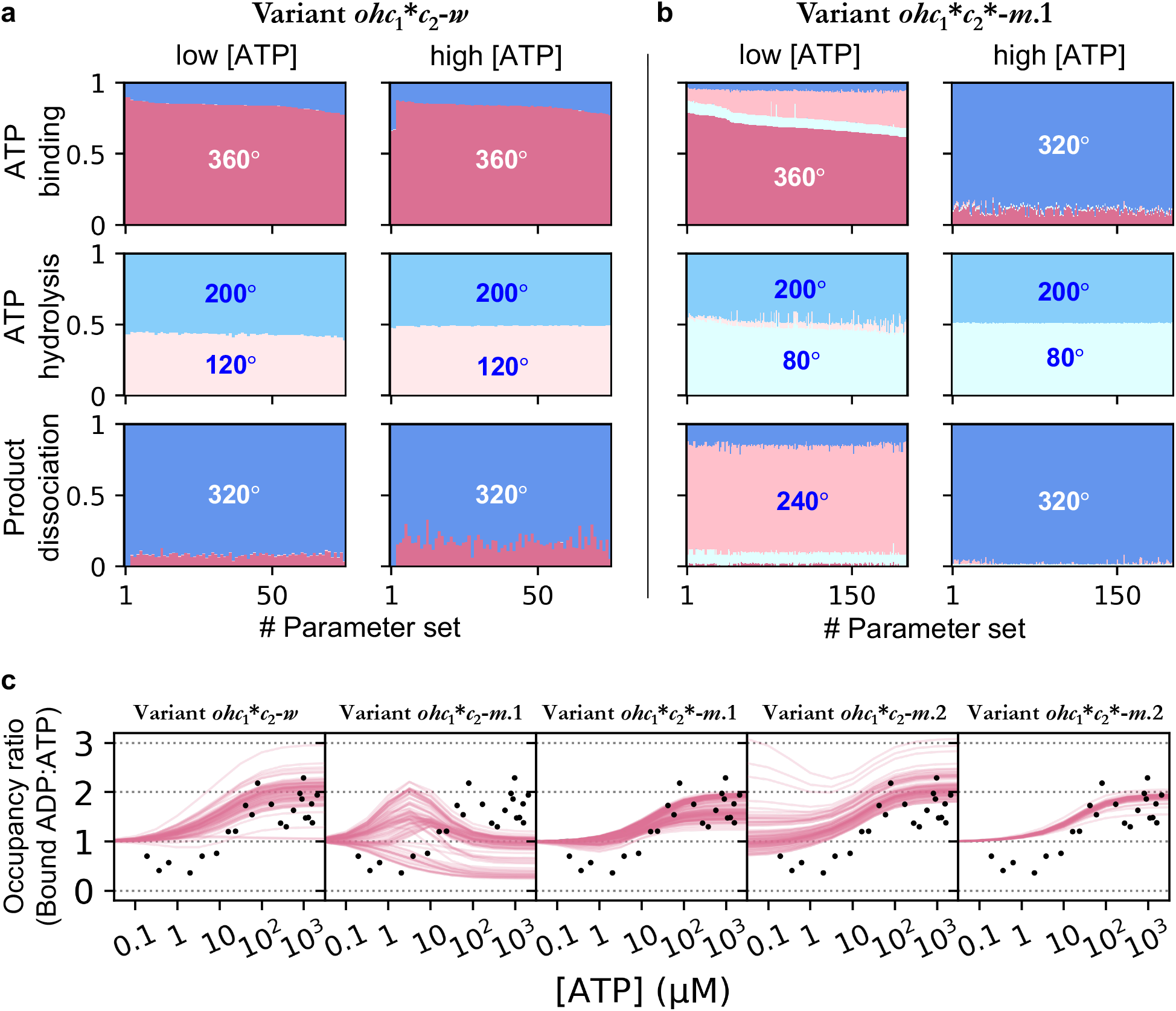
Cross-validation of the *ohc*_1_*c*_2_-variants. **a, b** Second cross-validation test: predicted probabilities of the major catalytic steps at different γ-orientations for two representative variants, (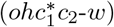) in (a) and (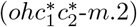) in (b). The representation is consistent with Fig. 2e. **c** Third cross-validation test: the predicted ratio of bound ADP to ATP (pink curves) for all five tested *ohc*_1_*c*_2_-variants, compared against the validation data from tryptophan fluorescence experiments (black dots) [20]. Source data are provided as a Source Data file.

As an additional cross-validation test, in Fig. 3c, we compared the predicted ratios of bound ADP to ATP (*ν*_D_ : *ν*_T_) against data from an early tryptophan fluorescence experiment [20]. The model predictions (pink curves) were tested against the experimental observation (black dots, adapted from Fig. 6 in Ref. [20]) that *ν*_*D*_ : *ν*_*T*_ ≈ 2 : 1 at saturating [ATP].

The variant 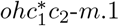 predicted *ν*_*D*_ : *ν*_*T*_ below or around 1:1 at high [ATP] (second panel), thus was excluded. In contrast, the other four variants reproduced this observed 2:1 ratio. Therefore, this cross-validation test leaves these four variants (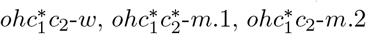, and 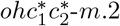) as candidates for a minimal model of F_1_-ATPase.

### The candidate variants also reproduce observed single-molecule trajectories

We finally tested if the four candidate *ohc*_1_*c*_2_-variants also agree with the single-molecule experiments where γ-subunit rotation was tracked over time using attached probes like actin filaments or colloidal gold beads [13, 14, 8]. To this end, we performed kinetic Monte-Carlo (KMC) simulations of these variants, closely resembling the experiments.

As depicted in Fig. 4a, the probe (e.g., a bead, yellow) acts as an external load coupled via a linker (modeled as a harmonic spring with force constant *κ*) to the γ-subunit (green). Following the experiments, we tracked the circular motion of the probe centroid, rather than the γ-subunit itself. Given the probe’s large size relative to F_1_-ATPase, its motion in the solvent was assumed to be overdamped and thus described by an overdamped Langevin equation (OLE) with rotational friction drag coefficient, *ξ* (Equation (21)). We estimated *κ* ≈ 5 *k*_B_*T* and *ξ* ≈ 6.7 × 10^−5^ *k*_B_*T* · s, based on experimental reports [13, 14, 8]. By coupling the KMC simulation of our Markov model with numerical integration of the OLE (Methods), we generated trajectories of the angular positions of the probe centroid comparable to those measured experimentally.

**Figure 4.**
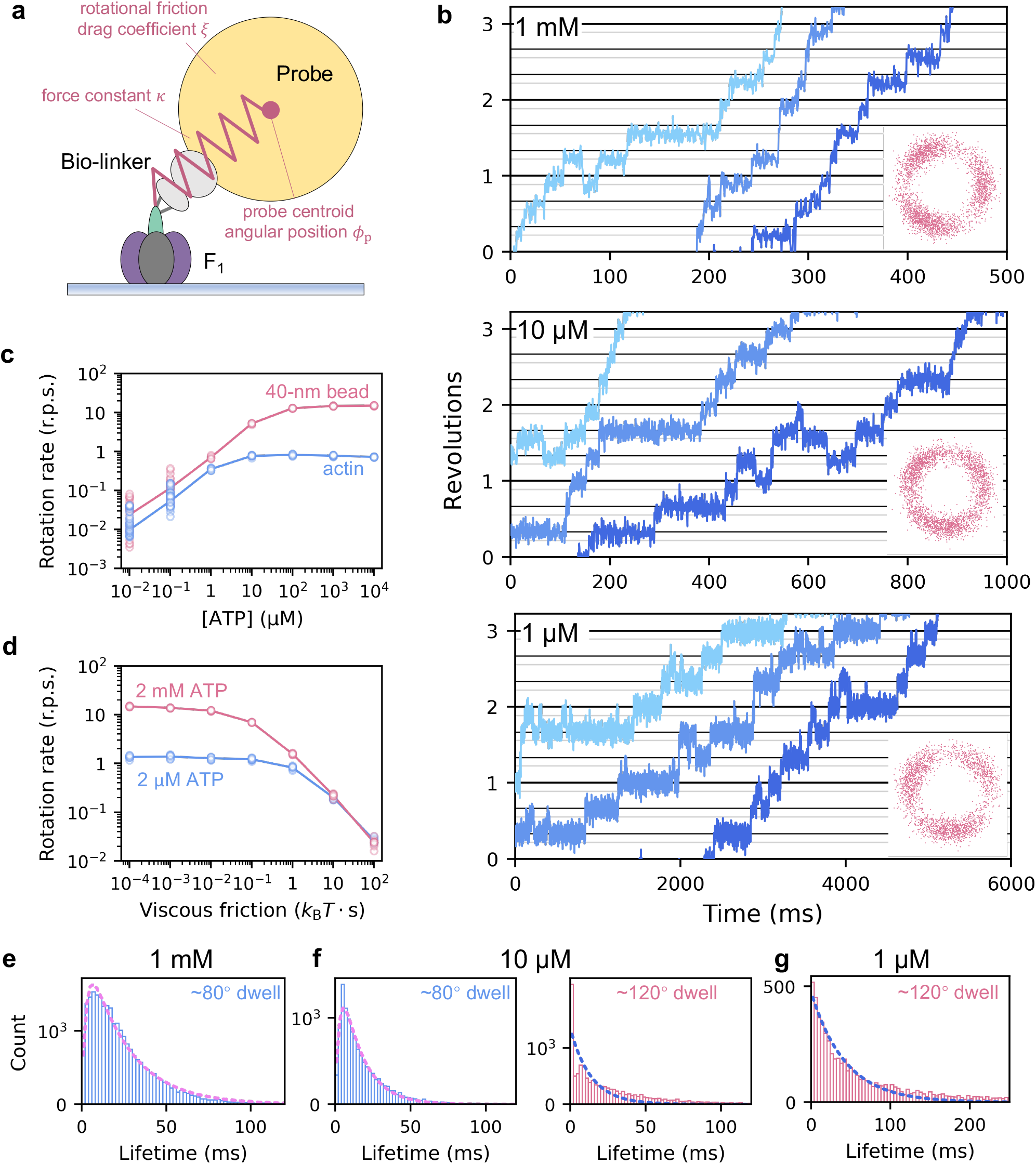
Kinetic Monte-Carlo simulations of the 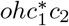 -*w* variant reproduce single-molecule observations. **a** Kinetic Monte-Carlo (KMC) simulation setup mimicking single-molecule experiments [8], where a probe (yellow) is coupled to the γ-subunit (green) via a bio-linker (force constant *κ*), and is subjected to rotational friction drag coefficient *ξ*. Adapted from Ref. [8] (cf. Fig. 1b). **b** Sample KMC trajectories of the probe rotation at 1 mM, 10 µM, and 1 µM ATP. In each panel, the three curves are continuous; later curves are shifted to save space. The gray horizontal lines are placed 40° below the black lines. Insets: angular positions of the probe taken from the sample trajectories and randomly placed around the unit cycle. The simulations qualitatively reproduce the experimentally observed discrete 80°- and 40°-substeps. **c** Predicted rotational rate versus ATP concentration for two loads: a 40-nm gold colloidal bead (pink), and a 1-µm actin (blue). **d** Predicted rotational rate versus viscous friction at high (2 mM, pink) and low ATP concentrations (2 µM, blue). In **c** and **d**, circles represent rates calculated from individual trajectories; solid curves are averages. Both **c** and **d** show good quantitative agreement with experimental measurements (cf. Figs. 2, 3 in Ref. [8]). **e**–**g** Dwell-time distributions (histograms) extracted from the simulated trajectories via Hidden Markov Analysis (Methods). **e** At 1 mM ATP, only ∼80°-dwells are identified. **f** At 10 µM ATP, both ∼80°-dwells and ∼120°-dwells are identified. **g** At 1 µM ATP, only ∼120°-dwells are identified. The dwell-time distributions for the 80°- and 120°-dwells are fitted by double and single exponential functions, respectively (violet and blue dashed curves). Source data are provided as a Source Data file.

The simulations of the variant 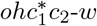 are reported in Fig. 4b–g. Fig. 4b shows sample trajectories for [ATP] ranging from 1 µM to 1 mM. These trajectories closely resemble those recorded experimentally (cf., e.g., Fig. 4 in Ref. [8]), notably reproducing the characteristic 80° and 40° rotational substeps. Furthermore, the dependencies of rotational rate on viscous friction and [ATP] (Fig. 4c, d) well agree with the single-molecule measurements (cf. Figs. 2, 3 in Ref. [8]). The other three candidate variants produced similar trajectories and dependencies that also match the experimental observations.

A more detailed quantitative analysis via hidden Markov model (Methods) of the simulated trajectories of the 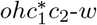 variant yielded dwell-time distributions (Fig. 4e–g) consistent with ATP binding limiting the 120° dwells and product dissociation limiting the 80° dwells, again matching the experimental conclusions [8, 15]. Given that all four candidate variants share similar chemo-mechanical coupling schemes (Fig. 3 and Supplementary Fig. 5), simulated trajectories, and dependencies, we expect them to also share similar dwell-time distributions. This overall consistency provides further cross-validation for all these four candidate variants.

## Discussion

Aiming at a minimal yet thermodynamically consistent model for F_1_-ATPases from two bacterial species (*E. coli* and *Bacillus* PS3), here we have developed a chemomechanical Markov model that integrates conformational and chemical degrees of freedom, as well as steric interactions between the γ- and β-subunits. We used a Bayesian training approach to infer the free energies and free energy barriers from the relevant experimental data extracted from literature [19, 21, 8, 75]. By systematic model comparison via our training + cross-validation strategy, we tested competing mechanistic hypotheses, particularly the three-conformation versus four-conformation hypotheses, and eventually identified several model variants with four distinct conformations of individual β-subunits (*ohc*_1_*c*_2_-variants) which all agree with and predict all relevant experimental data (Methods). These resulting candidates provide a coherent thermodynamic and kinetic framework that quantitatively explains the data for these ∼80°/ ∼40° sub-stepping enzymes.

Our study reconciles the decade-long controversy over whether a bi-site or a tri-site mechanism best describes ATP hydrolysis at full activity. In contrast to previous binding change models [20, 31, 22, 32, 5, 8, 33, 34, 35], which assume only one or at most very few catalytic pathways, our model involves by construction many parallel catalytic pathways, and predicts these to contribute to the overall activity with different fluxes that change with ATP concentration (Supplementary Fig. 6). At physiological (millimolar) ATP concentration, pathways resembling the tri-site mechanism [40, 52] dominate, wherein the total nucleotide occupancy alternates between three and transiently two. At micromolar ATP concentration, in contrast, bi-site pathways dominate, wherein the total occupancy alternates between one and transiently two. Accordingly, the total occupancy gradually decreases from three to one with decreasing ATP concentration, agreeing with published ATP titration data [19, 66, 67, 21].

Strikingly, we found no model variant with only three functionally distinct conformational states of each β-subunit (e.g., open, half-open, closed) that agrees with all relevant experimental data. We therefore conclude that a minimum of four functional states is required – and, from the above findings, also suffices – for proper F_1_-ATPase function. Indeed, it was proposed early-on that a tri-site mechanism requires at least four conformations [67]; furthermore, recent tryptophan fluorescence experiments [22] suggested the existence of at least four conformations with distinct nucleotide binding affinities. Particularly, several authors have postulated at least two different closed conformations, based on the observed structural and thermodynamic asymmetry of the three β-subunits [4, 5, 71], in line with our *ohc*_1_*c*_2_-variants. This idea is also supported by a principal component analysis of all available β-subunit structures (Supplementary Fig. 7), demonstrating considerable heterogeneity of those structures classified as closed.

Resolving another debate [81, 82, 48], our study strongly supports the notion that for a nucleotide-bound β-subunit, the catalytically active, closed conformation is energetically more favorable than the open and partially open conformations. In fact, according to our Markov model, this energetic property of the β-subunits is a necessary condition for establishing tight coupling between catalysis and rotation (see Fig. 2f and Supplementary Note 1). A plausible explanation is that under this condition, a β-subunit is unlikely to open spontaneously and release the product after ATP hydrolysis, unless forced by the γ-subunit, thereby ensuring tight chemo-mechanical coupling. Conversely, if the closed conformation for a nucleotide-bound β-subunit were not the most stable one, the β-subunits could go through catalytic cycles spontaneously and independent of the γ-subunit orientation, resulting in a catalytic rate higher than the rotational rate and a correspondingly reduced coupling efficiency. Notably, this energetic property was observed independently in previous atomistic simulation studies [81, 82], where the β-subunit that was initially kept open by the bulge of the γ-subunit closed spontaneously after removing the bulge through enforced rotation of the γ-subunit.

Our model quantitatively reconciles the classical binding change paradigm with the inherent stochastic nature of rotary catalysis. In line with Arai et al. [29], our model by construction conceptualizes the γ-subunit rotation as random thermal fluctuations which are neither directly nor deterministically coupled to ATP hydrolysis (Supplementary Table 2). In this sense, our model represents a Brownian-ratchet mechanism, which is able to explain the complex and intriguing stochastic rotational behaviors of the central stalk at low ATP concentrations observed in single-molecule experiments [8, 52, 15]. First, in our kinetic Monte-Carlo simulations, the γ-subunit often rotates back and forth stochastically over several 120° steps, directly mirroring the irregular rotations observed experimentally [52, 15]. Remarkably, when no nucleotide is bound within the three catalytic sites, the simulated irregular rotations can accumulate up to several revolutions. This prediction directly supports the hypothesis that the irregular rotations represent “Brownian hopping among the three equivalent orientations in a nucleotide-free F_1_-ATPase” [52]. Second, our model predicts that one ATP hydrolysis event can trigger larger than 120° rotations, explaining the experimental measurements somewhat misleadingly termed over 100% coupling efficiencies [8]. Notably, these events do not violate energy conservation; they occur only under sufficiently low external load, meaning the work performed even over several 120° steps remains smaller than the free energy released by one ATP hydrolysis. Rather, these results are perfectly in line with a Brownian-ratchet mechanism [30, 29]. Specifically, at low ATP concentrations where the nucleotide occupancy is only one or less, large rotational fluctuations occur almost uncoupled from ATP hydrolysis. As nucleotide exchange alters the affinity landscape of the catalytic subunits, these stochastic fluctuations are directionally rectified, allowing one hydrolysis event to trigger a full revolution in the absence of mechanical load.

With increasing ATP concentration and nucleotide occupancy, the rotation becomes more tightly coupled to ATP hydrolysis. Irregular rotations rarely occur, and the coupling efficiency approaches 100% (Fig. 2 and Supplementary Figs. 3, 4). In this high-ATP-concentration regime, the macroscopic behavior of the F_1_-ATPase becomes more similar in spirit to the deterministic picture as in classical binding change models, functioning apparently like a power-stroke. Taken together, the agreement of our model with multiscale experimental data – including both the microscopic, stochastic single-molecule observations [8, 52, 15] and the macroscopic, deterministic measurements [19, 21] – demonstrates that the classical binding change paradigm is fully compatible with the microscopic stochasticity of this molecular machine, which underscores that an intrinsic Brownian-ratchet mechanism is both feasible and sufficient for rotary catalysis.

Although the four candidate models agree with all relevant experimental data, and although they share many other properties, they do differ in mechanistic details (Table 4). An example is which of the two closed β-subunits in the classic crystal structure nomenclature [4], β_TP_ or β_DP_, is the high affinity site (third column); here, the experimental evidence is still inconclusive, with some results pointing to β_TP_ [72, 31, 83] and others to β_DP_ [4, 5]. Similarly, the number of β-subunits that can be simultaneously catalytically active varies between one and two among the four variants (fourth column); experimental evidences supporting both possibilities have been reported [71, 5, 72]. Therefore, based on these current results, we cannot conclude which of these four candidates is more or less likely. As of now, it seems to us that the available experimental data does not allow to distinguish between these different mechanistic details.

**Table 4.** Comparison of the four candidate models.

| Variant | Representative functional states of the catalytic ($80^\circ$) and ATP-waiting ($120^\circ$) dwells | High affinity site | Catalytically active β-subunit(s) in the catalytic dwell |
| --- | --- | --- | --- |
| $ohc_1^*c_2-w$ | $80^\circ$: $hc_1c_2, oc_1c_2$;<br>$120^\circ$: $oc_2c_1$ | $\beta_{DP} (c_1^*)$ | Only $\beta_{DP}$ |
| $ohc_1^*c_2^*-m.1$ | $80^\circ$: $hc_1c_2, oc_1c_2$;<br>$120^\circ$: $ohc_1, ooc_1$ | $\beta_{DP} (c_1^*)$ | Both $\beta_{TP}$ and $\beta_{DP}$ |
| $ohc_1^*c_2-m.2$ | $80^\circ$: $hc_1c_2, oc_1c_2$;<br>$120^\circ$: $ohc_2, ooc_2$ | $\beta_{TP} (c_2)$ | Both $\beta_{TP}$ and $\beta_{DP}$, although $\beta_{TP}$ energetically favors the inactive $c_2$ conformation |
| $ohc_1^*c_2^*-m.2$ | $80^\circ$: $hc_1c_2, oc_1c_2$;<br>$120^\circ$: $ohc_2, ooc_2$ | $\beta_{TP} (c_2^*)$ | Both $\beta_{TP}$ and $\beta_{DP}$ |

It also remains a challenge to interpret the functional roles of experimentally resolved atomistic structures within the F_1_-ATPase catalytic cycle [84, 70, 85, 86, 63], particularly given the differing cryoEM structures reported by Nakano et al. [65] and Sobti et al. [32] for the ∼80° catalytic and ∼120° ATP-waiting dwells (Fig. 9c– f). Whereas these structures share similar nomenclatures fully compatible with single-molecule FRET measurements [70], there is a key difference regarding the conformational progression of the β_1_-subunit upon the 80° → 120° rotation: does it open further (as suggested by Nakano et al. [65], Supplementary Fig. 9c,d), or close partially (as suggested by Sobti et al. [32], Supplementary Fig. 9e,f)? Our model provides a thermodynamic framework to resolve this ambiguity by mapping these structures to distinct functional stages within the catalytic cycle. Specifically, for the joint functional states of the three β-subunits in the 80°- and 120°-dwells, our model predicts that the most populated pathway involves the β_1_-subunit opening (*h* → *o*) and decreasing in nucleotide binding affinity to facilitate product dissociation [6, 15, 73, 70, 22] (Table 4, second column; Supplementary Figs. 9, 10a,b). Along this pathway, the 80°-dwell is most populated by an *hcc* state (Supplementary Fig. 9a), representing the stage where the β_1_-subunit awaits product dissociation. Upon rotation to 120°, the enzyme converts to an *ohc* state (Supplementary Fig. 9b), representing the ATP-waiting stage. We note that in this context, the *oc*_2_*c*_1_ state predicted by the 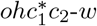 variant can also be interpreted as an *ohc*-like state (Supplementary Note 7).

Notably, the Nakano structures (Supplementary Fig. 9c,d) align well with this predicted progression: despite both being labeled open (β_E_), the catalytic interface of the β_1_-subunit appears more compact at 80° than at 120°. Therefore, we suggest to assign the Nakano structures to these predicted primary functional stages (*hcc* and *ohc*). In contrast, the Sobti structures (Supplementary Fig. 9e,f) suggest an opposite *o* → *h* progression: here, the β_1_-subunit structure is less compact at 80° than at 120° (β_O_ versus β_HC_). However, as demonstrated by our analysis of Family B variants which predict such a progression as the dominant pathway (Results), this progression would imply an affinity increase upon the 80° → 120° rotation, in contrast to what has been measured previously [22], as well as to analysis of single-molecule data [8, 15]. Furthermore, several structures resembling the Nakano *ohc*-like structure (Supplementary Fig. 9d) have been reported for various modified F_1_-ATPase constructs in the ATP-waiting dwell [64, 78, 79], featuring a β_1_-subunit consistently more open than the Sobti β_HC_ structure at 120° (Supplementary Fig. 9f). Given these discrepancies, we suggest that the Sobti structures likely capture different functional stages. For the 80°-dwell, we would assign their structure (Supplementary Fig. 9e) rather to our the *occ* state of our model (Supplementary Fig. 9a), which is transient in the native cycle, appearing only after product dissociation in the β_1_-subunit but before ATP hydrolysis completes in the β_2_-subunit. This assignment raises the question why this transient structure was actually seen in the cryoEM sample, which can be explained by the use of a mutant that intentionally slowed down the hydrolysis step [32], thereby sufficiently stabilizing this structure. Similarly, for the 120°-dwell, the Sobti structure (Supplementary Fig. 9f) might represent an intermediate captured shortly after ATP binding, as the β_1_-subunit begins to close.

Taken together, our model integrates these differing structures into a unified mechanistic picture. For the 80°-dwell, the Nakano *hcc*-like (Supplementary Fig. 9c) and the Sobti *occ*-like structures (Supplementary Fig. 9e) likely correspond to the pre- and post-product-dissociation stages, respectively, consistent with the observation of two distinct rate-limiting steps in single-molecule experiments [8, 15]. For the 120°-dwell, the Nakano *ohc*-like (Supplementary Fig. 9d) and Sobti *hhc*-like structures (Supplementary Fig. 9f) likely represent the pre- and post-ATP-binding stages, respectively. This interpretation requires the involved structures to be rather flexible, which is in line with previously observed large fluctuations of the γ-subunit within this dwell [84, 15].

Striking the best balance between model complexity and sufficiently few model parameters is a notorious challenge. Building upon our minimal model, several routes exist for future refinement and extension. For example, our minimal model assumes that the steric repulsions between the γ- and β-subunits are the main (and only) determinants for efficient chemo-mechanical coupling. It is known, however, that weaker effects such as allosteric communication within the α_3_β_3_ ring or finer details of the γ-β interaction landscape (e.g., electrostatics) also contribute and possibly modify the mechanism of the chemo-mechanical energy transduction. One line of evidence is the structural asymmetry and residual activity of γ-depleted and γ-truncated F_1_-ATPase constructs [76, 77, 78, 79]. It would therefore be a tempting future extension of our model to also include energy terms – of course requiring additional parameters – which describe these weaker and, in our view, secondary effects. Other possible extensions include explicitly modeling phosphate to dissect the timing and kinetics of its release; testing the model against a wider range of experimental data, such as those from mutant F_1_-ATPase [34, 15], and potentially moving beyond binding change models to consider continuous rotation of the γ-subunit.

While our minimal Markov model was developed specifically for bacterial F_1_-ATPases, the underlying Brownian-ratchet picture can serve as a universal principle for chemo-mechanical energy transduction across the entire rotary ATPase superfamily, including F-ATPase from other species [61, 62, 63] as well as V- and A-ATPases [23, 24, 25, 26, 27, 28, 30, 29]. For the specific example of bacterial F-ATPase our study demonstrates how this general principle can be implemented in quantitative and thermodynamically consistent terms. Similarly, for each of the other rotary ATPases, a fine-tuned and high-dimensional free energy landscape underlies and supports this principle, which, in our model, is explicitly quantified by the parameters including free energies and activation barriers.

While the fundamental energy transduction principle is shared, these rotary ATPases differ significantly in subunit complexity, regulatory mechanisms, and rotational sub-stepping behaviors [23, 24, 25, 26, 27, 28, 30, 29]. These differences imply that the underlying energy landscapes are fine-tuned differently by evolution, and, accordingly, respective Markov models will be defined by different free energy levels and barrier heights. Determining these will require similarly comprehensive measurements as were available for F-ATPase, and it will be exciting to compare the obtained different free energy landscapes to further extract common fundamental principles as well as differences between families and between species, thereby revealing how evolution has tuned their stabilities, binding affinities, catalytic rates, and coupling efficiencies to optimally adapt their functional mechanisms to the respective biological demands.

Beyond rotary ATPases, the theoretical framework established in this study for developing thermodynamically consistent Markov models is very general and therefore can be applied to the broad range of macromolecular machines that combine conformational motions with ligand binding/unbinding or enzymatic reactions to achieve their function. Particularly, the use of thermodynamically consistent Markov models which include both all relevant states as well as conformational and catalytic pathways should help to overcome the limitations and sometimes misleading interpretations of simpler, Michaelis-Menten-like descriptions that consider only a few presumed pathways and apparent binding affinities. For more complex and conformationally coupled enzyme dynamics, the latter may be insufficient, as testified by the bi-site versus tri-site controversy.

Finally, the systematic, data-driven model comparison approach demonstrated here also serves to assess competing mechanistic hypotheses against available experimental data. An essential aspect of this approach is a training + cross-validation strategy, which, as detailed in Methods, renders complex systems computationally tractable while ensuring thermodynamic consistency.

## Methods

### Mathematical formulation of the chemo-mechanical Markov model

Here, we present the detailed mathematical formulation for our chemo-mechanical Markov model, including the definitions of the Markov state space, the free energies of the Markov states, and the transition rates, as well as the master equation governing the time evolution of the system.

Each chemo-mechanical state (Markov state) **s** in our model is uniquely defined by a seven-dimensional vector specifying the values of the seven DOFs introduced previously (Table 1):

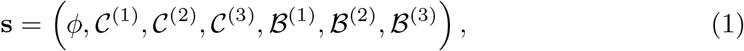

where *ϕ* is the γ-subunit orientation, 𝒞^(*k*)^ is the conformation of the *k*-th β-subunit, and ℬ^(*k*)^ is its nucleotide binding state (*k* = 1, 2, 3). The complete set of Markov states includes all combinatorially possible combinations of these seven DOFs. Formally, this state space is constructed by the Cartesian product of the seven DOFs.

For each Markov state, its free energy (state free energy) is decomposed into a sum of the conformational free energies of individual β-subunits, their binding free energies, and the interaction energies between the γ- and β-subunits:

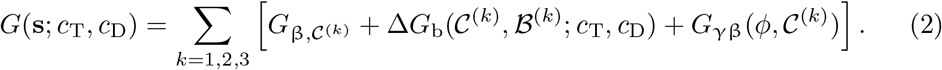

Here, a free energy *G*_β, 𝒞_ is assigned to each β-subunit conformation (𝒞 = *o, h, c* for *ohc*-variants; 𝒞= *o, h, c*_1_, *c*_2_ for *ohc*_1_*c*_2_-variants), and is defined as a model parameter (conformational free energies, Table 5).

**Table 5.**
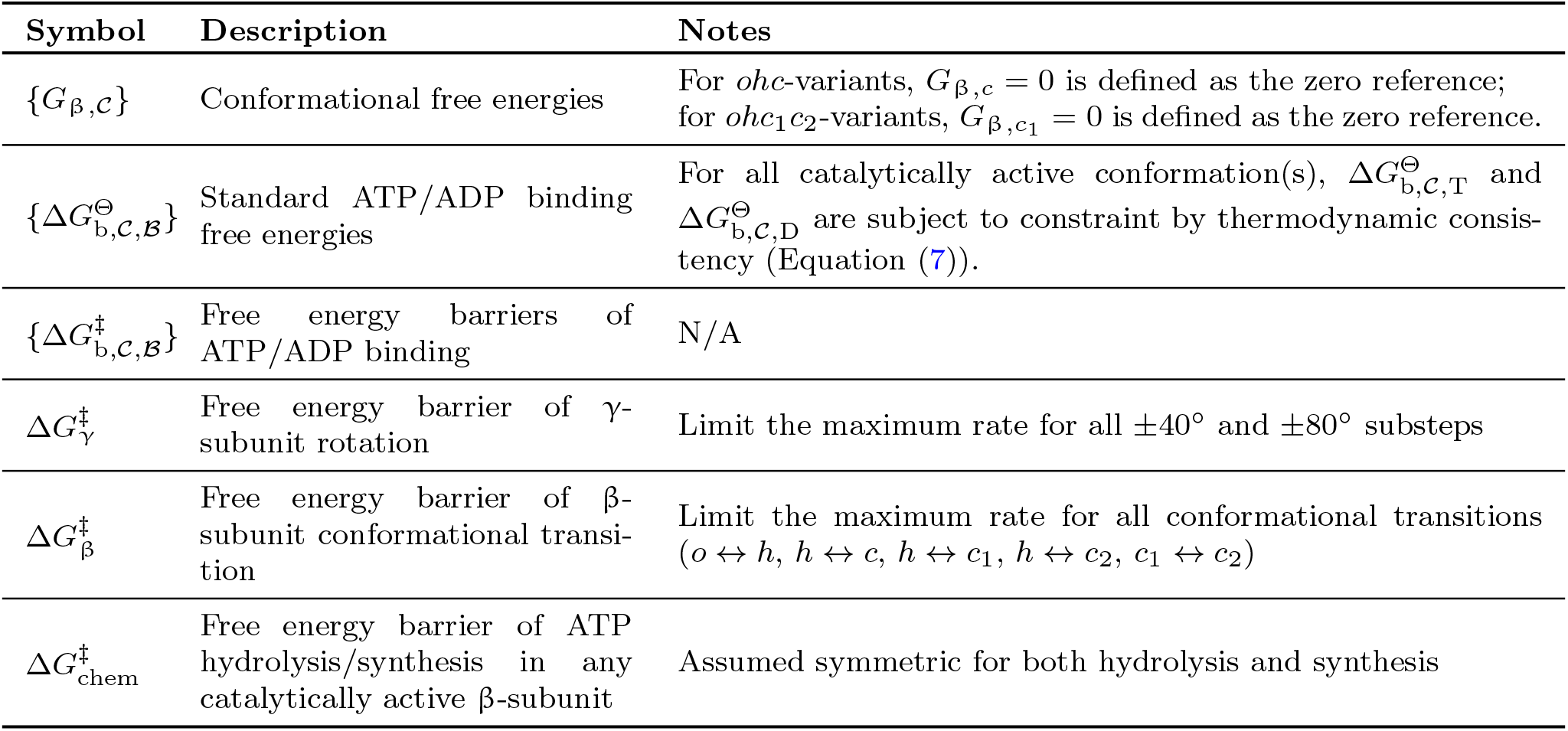
Model parameters. The β-subunit conformation 𝒞 = *o, h, c* (*ohc*-variants) or 𝒞 = *o, h, c*_1_, *c*_2_ (*ohc*_1_*c*_2_-variants). The binding state ℬ = T, D for ATP and ADP, respectively. The catalytically active conformations refer to *c* for *ohc*-variants, *c*_1_ for 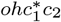 -variants, and both *c*_1_ and *c*_2_ for 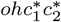 -variants.

The binding free energy, Δ*G*_b_(𝒞 ^(*k*)^, ℬ ^(*k*)^; *c*_T_, *c*_D_), which depends on the bulk nucleotide concentrations *c*_T_ and *c*_D_, is defined by:

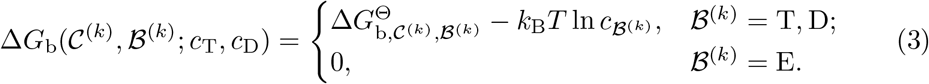

Here, two free energies 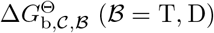 are assigned to each β-subunit conformation for ATP and ADP binding, respectively, under the standard condition of 1 mM ATP/ADP. They are also defined as model parameters (standard ATP/ADP binding free energies, Table 5).

The γ-β interaction energy *G*_γβ_(*ϕ*, 𝒞^(*k*)^) primarily models the steric repulsions between the γ- and β-subunits. As introduced in Results, we use a strong simplification, assuming an infinitely high energy for those sterically disfavored conformations, thus precluding them from being populated; the remaining conformations are thermodynamically accessible. The hyperparameter γ-β restrictions, also introduced previously, specifies the set of accessible conformations, 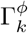, for each β-subunit (β_*k*_), at each γ-orientation (*ϕ*). Formally, this interaction energy *G*_γβ_(*ϕ*, 𝒞 ^(*k*)^) is defined by:

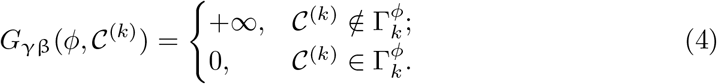

Additionally, for Family A variants (Supplementary Equation (7) in Supplementary Table 4), a slightly favorable interaction energy of − 2.3 *k*_B_*T* is assumed for the β_*k*_-subunit adopting the *o* conformation at γ-orientation *ϕ* = 120° × *k* (*G*_γβ_(*ϕ*, 𝒞^(*k*)^) = − 2.3 *k*_B_*T* if *ϕ* = 120° × *k* and 𝒞^(*k*)^ = *o*), describing the ∼120° ATP-waiting dwell observed in single-molecule experiments [8, 15].

The transition rates for all elementary transitions included in our model are defined in Supplementary Table 2. These definitions are grounded in transition state theory and the principle of local detailed balance, as outlined below.

The transition rate from a Markov state *i* to another Markov state *j* in our model, denoted *r*_*i*→*j*_, is determined by the corresponding rate coefficient *k*_*i*→*j*_ and the molecularity (reaction order) of the process. According to transition state theory, the rate coefficient *k*_*i*→*j*_ is determined by the free energy barrier 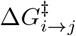:

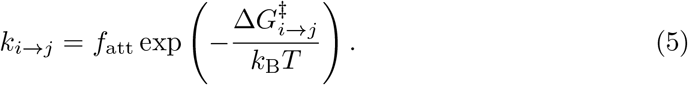

Here, *f*_att_ is a constant, the attempt frequency. It serves as a global pre-exponential factor; its specific value (here chosen to be 10^9^ s^−1^) is not critical, as any different choice would be absorbed by a uniform shift in the fitted free energy barriers.

To enforce the principle of local detailed balance for every elementary transition (Results), the backward rate coefficient *k*_*j*→*i*_ (from state *j* to state *i*) is related to *k*_*i*→*j*_ by:

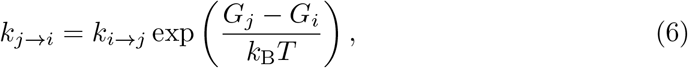

where *G*_*i*_, *G*_*j*_ are the free energies of the two states.

For a uni-molecular (first-order) transition, e.g., rotation of the γ-subunit, *k*_*i*→*j*_ defined above is a first-order rate coefficient, while the transition rate *r*_*i*→*j*_ = *k*_*i*→*j*_. For a bi-molecular (second-order) transition, e.g., nucleotide binding, *k*_*i*→*j*_ defined above is a second-order rate coefficient, while *r*_*i*→*j*_ is the product of the *k*_*i*→*j*_ and the nucleotide concentration. Note that, because our model describes the internal dynamics of a single F_1_-ATPase molecule, *r*_*i*→*j*_ does not need to include its own concentration.

Notably, as demonstrated in Supplementary Table 2, to define the large number of transition rates without introducing an excessive amount of free parameters, we made simplifying assumptions about the energy barriers. For example, we assume a uniform free energy barrier 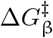 governs all β-subunit conformational transitions (Supplementary Equation (1) in Supplementary Table 2), limiting their maximum rate to 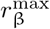.

Another important assumption is that the rates of ATP hydrolysis and synthesis in a catalytically active β-subunit are equal. This assumption is based on experimental evidence suggesting that the hydrolysis and synthesis reactions are approximately in equilibrium within the catalytic site [6, 66, 67, 68]. To ensure overall thermodynamic consistency, this assumption implies that the difference between the standard ATP and ADP binding free energies must exactly compensate for the free energy of ATP hydrolysis in solution. We therefore enforce this condition by imposing a corresponding constraint on the standard binding free energies: for any catalytically active conformation 𝒞,

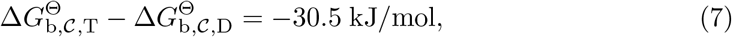

where −30.5 kJ/mol is the standard free energy of ATP hydrolysis in solution [87].

These transition rates are used to construct the transition rate matrix, **R**. The time evolution of our Markov model, modeled as a continuous time Markov process, is described by the master equation:

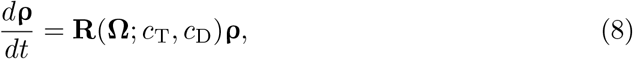

where **ρ** is the vector of the populations of the Markov states. The off-diagonal element of **R** represents the transition rate from state *j* to state *i*: *R*_*ij*_ = *r*_*j*→*i*_ (*i* ≠ *j*); the diagonal element *R*_*ii*_ = − ∑_*j*≠ *i*_ *r*_*i*→*j*_ ≡ *r*_*i*_ is the total rate leaving state *i*. The transition rates *R*_*ij*_ depend on the model parameters (Table 5) collectively denoted as a vector **Ω**, and the bulk nucleotide concentrations *c*_T_ and *c*_D_.

### Model hyperparameters and parameters

In summary, our Markov model includes three hyperparameters. These hyperparameters have been introduced conceptually in Results, from which specific variants of our model are derived to test competing hypotheses, regarding:

1. The set of distinct β-subunit conformations (Three-conformation vs. Four-conformation). Does F_1_-ATPase function require a minimal number of three functionally distinct conformations for individual β-subunits, or four conformations? The choices are 𝒞^(*k*)^ ∈ {*o, h*, } (*ohc*-variants), or 𝒞^(*k*)^ ∈ {*o, h, c*_1_, *c*_2_} (*ohc*_1_*c*_2_-variants).
2. The catalytically active conformation(s) (One-active vs. Two-active). Within *ohc*_1_*c*_2_-variants, are both closed conformations (*c*_1_, *c*_2_) catalytically active, or only one? The choices are that either only *c*_1_ is active (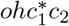 -variants), or both *c*_1_ and *c*_2_ are active (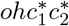 -variants).
3. The γ-β restrictions. How does the γ-subunit sterically restrict β-subunit conformations at each orientation? For *ohc*-variants, each 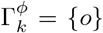, {*o, h*}, or {*o, h, c*}; for *ohc*_1_*c*_2_-variants, each 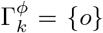, {*o, h*}, {*o, h, c*_1_}, {*o, h, c*_2_}, or {*o, h, c*_1_, *c*_2_}. Each combination of all six 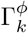 ultimately defines a specific variant (see, e.g., Tables 2 and 3).

Here, we introduce a more formal notation for the hyperparameter γ-β restrictions. Specifically, we define 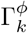, the ensemble of conformations accessible to each β-subunit β_*k*_ at each γ-orientation *ϕ*. Due to the three-fold rotational pseudo-symmetry of F_1_-ATPase structure, specifying six 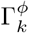, e.g., 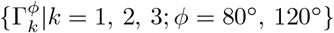, is sufficient to uniquely define a specific variant, as demonstrated in Fig. 2f and Tables 2, 3. Because these γ-β restrictions are defined to primarily model steric repulsion, a more open conformation should not be precluded when a more closed conformation is allowed. Accordingly, we assume that for *ohc*-variants, each 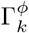 can be {*o*}, {*o, h*}, or {*o, h, c*}. As a rough estimation, the combinations of the six 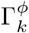 give rise to 3^6^ − 2^6^ = 665 *ohc*-variants; here, 2^6^ combinations where *c* is completely missing are excluded. Similarly, for *ohc*_1_*c*_2_-variants, 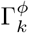 can be {*o*}, {*o, h*}, {*o, h, c*_1_}, {*o, h, c*_2_}, or {*o, h, c*_1_, *c*_2_}. Their combinations give rise to roughly 5^6^ − 2^6^ − (3^6^ − 2^6^) − (3^6^ − 2^6^) = 14, 231 *ohc*_1_*c*_2_-variants. Here, the subtractions exclude the 2^6^ combinations where both *c*_1_ and *c*_2_ are completely missing, the 3^6^ − 2^6^ combinations missing *c*_1_, and the 3^6^ − 2^6^ combinations missing *c*_2_.

Finally, the parameters that are introduced into our model when defining the state free energies (Equations (2), (3)) and transition rates (Supplementary Table 2) are summarized in Table 5, including conformational free energies, binding free energies, and several free energy barriers. They count to 17 parameters for the *ohc*-variants and 22 parameters for the *ohc*_1_*c*_2_-variants (some are subjected to the constraint of Equation (7)).

### Derivation of statistical and kinetic observables from the non-equilibrium steady-state

Several statistical and kinetic observables are derived from the steady-state solution of the master equation (Equation (8)), including (1) nucleotide occupancies, (2) populations of the 80°- and 120°-dwells, (3) turnover rates, rotation rates, and chemomechanical coupling efficiencies, and (4) the probabilities that a major catalytic step occurs at different γ-subunit orientations. These derived observables constitute the primary predictions of our model, and are systematically compared against experimental data during the training and cross-validation procedures (detailed in Results and Methods).

In this derivation, we assume the nucleotide concentrations *c*_T_ and *c*_D_ are constant. As mentioned in Results, this setup directly mimics typical experimental conditions (e.g., single-molecule experiments) where F_1_-ATPase is in contact with external chemical reservoirs [8]. More generally, this assumption also serves as a valid approximation for typical bulk biochemical assays, where nucleotide concentrations are substantially higher than the enzyme concentration, [19, 21], rendering substrate depletion negligible. Consequently, this derivation describes an open system driven into a non-equilibrium steady-state (NESS) by the constant ATP/ADP chemical potential difference.

The steady-state populations of the Markov states under the NESS condition, **ρ**^st^, are obtained as the null space of the transition rate matrix **R**, i.e., by solving the equation

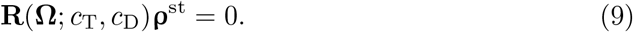

Although the steady-state populations **ρ**^st^ remain constant, the NESS condition established above allows a persistent net flux to flow through the network of states along thermodynamically favorable pathways. This net flux is defined for each transition between states *i* and *j* as:

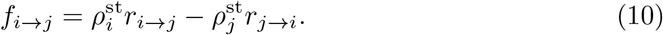

It is the summation of these net fluxes over relevant transitions that generates the macroscopic observables like turnover and rotation rates, even though every elementary transition strictly satisfies local detailed balance (Equation (6)).

Equation (9) is solved numerically by eigenvalue decomposition of the transition rate matrix **R**. One eigenvalue of **R** is zero; the corresponding eigenvector, after normalization, gives the steady-state populations **ρ**^st^. Subsequently, {*f*_*i*→*j*_} are calculated via Equation (10). From these quantities, several basic observables can be directly evaluated, as summarized in Supplementary Table 6.

Based on these basic observables, we then define several composite observables that are used in the training and cross-validation procedures (Results). The total nucleotide occupancy *ν* is the sum of individual ATP and ADP occupancies:

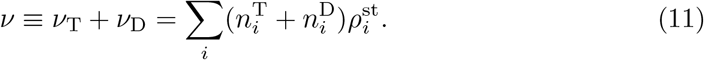

The chemo-mechanical coupling efficiency *η*, a key measure of the motor’s performance, is defined as the ratio of the rotational rate (scaled by 3 for a full cycle) to the turnover rate:

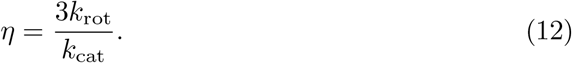

Finally, to compare our model prediction against the consensus chemo-mechanical coupling scheme (Table 6 and Supplementary Fig. 2) in the cross-validation procedure, we calculate the probability 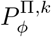 that a major catalytic step Π (ATP binding, ATP hydrolysis, or product dissociation) occurs within the β_*k*_-subunit at a given γ-orientation *ϕ*. This probability is calculated by normalizing the orientation- and site-specific rate 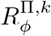 in Supplementary Table 6 by the average turnover rate per site (*k*_cat_*/*3, considering the three-fold pseudo-symmetry of F_1_-ATPase):

**Table 6.** Experimental data used for training and cross-validation. All experimental data used for training and cross-validation were obtained from studies on F_1_-ATPase from two bacterial species, *Escherichia coli* (*E. coli*) and the thermophilic *Bacillus* PS3 (EF_1_, TF_1_). Both EF_1_ and TF_1_ have been shown to exhibit similar well-characterized 80°/40° sub-stepping pattern [8, 15, 59], suggesting mechanistic consistency. Note that some studies employed specific F_1_-ATPase mutants (detailed in the footnotes below); however, these mutations have been shown to have only minor effects on enzymatic activity [19, 20, 13, 14, 8].

| Observable | Details | Source species |
| --- | --- | --- |
| *Training Data* |  |  |
| Turnover rates | $k_{\text{cat}}([\text{ATP}])$ at $[\text{ATP}] = 1 \mu\text{M}, 10 \mu\text{M}, 0.1 \text{ mM},$ and $1 \text{ mM}$, derived from Michaelis-Menten constants $V_{\text{max}} = 80 \text{ s}^{-1}$ and $K_{\text{M}} = 0.04 \text{ mM}$ [19]. | *E. coli* * |
| Chemo-mechanical coupling efficiency | A chemo-mechanical coupling efficiency $\eta = 100\%$ is used as the training target for $[\text{ATP}]$ from micromolar to millimolar. This target value is consistent with the long-standing view of the enzyme as a near-perfect reversible motor [6, 80], and also strongly supported by the single-molecule data [8, 54]. | Generally applicable |
| ATP/ADP titration curves | Measurements from tryptophan fluorescence experiments reported in Ref. [21] (data points recovered from Fig. 1A). | *E. coli* * |
| *Validation Data* |  |  |
| Population shift between 80°- and 120°-dwells | Single-molecule experiments show that the steady-state population of F<sub>1</sub>-ATPase shifts from the 120°-dwells (ATP-waiting dwells) to the 80°-dwells (catalytic dwells) as $[\text{ATP}]$ increases from sub-micromolar to millimolar levels [8]. | *Bacillus* PS3† |
| Known γ-orientations of the major catalytic steps (consensus chemo-mechanical coupling scheme) | A consensus chemo-mechanical coupling scheme is synthesized from a broad range of experimental studies [8, 54, 33, 15, 57, 34, 70, 20, 21, 22, 32, 65, 56, 55, 67, 66, 75] (visualized in Supplementary Fig. 2; see also Supplementary Note 2), specifying the γ-orientations of the major catalytic steps: (i) under low $[\text{ATP}]$, ATP binding in a β-subunit occurs at a 120°-dwell (defined as 0° for reference); in this β-subunit, subsequently, (ii) ATP hydrolysis occurs at +200°; (iii) under high $[\text{ATP}]$, the rate-limiting product dissociation step occurs at +320°. | *E. coli* and *Bacillus* PS3 |
| Ratio between ADP and ATP occupancies | Tryptophan fluorescence experiments show that at saturating $[\text{ATP}]$, the ratio of bound ADP to bound ATP ($(\nu_D : \nu_T)$) is approximately 2:1 [20]. | *E. coli*§ |

\* Data obtained using *E. coli* F<sub>1</sub>-ATPase carrying the $\beta$ -Y331W mutations [19, 21]
† Data obtained using *Bacillus* PS3 F<sub>1</sub>-ATPase carrying the following mutations: $\gamma$ -S107C, $\gamma$ -I210C, and $\alpha$ -C193S; an extra his-tag (consisted of ten histidines) at the amino terminus of each β-subunit [13, 14, 8]
§ Data obtained using *E. coli* F<sub>1</sub>-ATPase carrying the $\beta$ -F148W mutations [20]

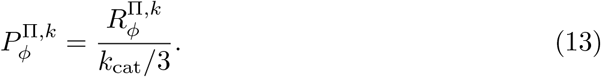

The probabilities 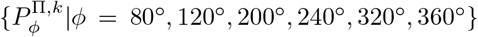 visualized in Figs. 2e and 3a,b quantify how a catalytic step Π is distributed over all possible γ-orientations as catalysis proceeds in the given catalytic site, thus characterizing the stochastic nature of F_1_-ATPase rotary catalysis.

### Systematic model comparison strategy

To objectively select the optimal model variant that is consistent with all experimental data on bacterial F_1_-ATPase, we employ a systematic model comparison strategy of training + cross-validation. This strategy involves partitioning the available experimental data of bacterial F_1_-ATPase into two non-overlapping sets for training and cross-validation, respectively. These two sets of data, termed training data and validation data, are summarized in Table 6.

In the training procedure, we employ a Bayesian approach (detailed below) to broadly sample the high-dimensional parameter space (Table 5) and obtain a large ensemble of parameter sets that reproduce the training data within experimental uncertainty. These training data are quantitative measurements of several macroscopic properties that are fundamental to F_1_-ATPase function, namely turnover rates, chemo-mechanical coupling efficiency, and nucleotide titration curves. After training, an ensemble of parameter sets may be identified that reproduce the basic rotary catalysis behavior of F_1_-ATPase as core functional output.

Subsequently, we perform cross-validation, by comparing the predictions of these trained parameter sets to the validation data. These validation data include qualitative to semi-quantitative observations that provide further insights into the mechanistic details of F_1_-ATPase on a more microscopic scale, mostly from single-molecule experiments, namely population shifts, consensus chemo-mechanical coupling scheme, and ratio between ADP and ATP occupancies. This cross-validation procedure allows us to further identify trained parameter sets that are also consistent with known mechanistic details. A model variant is considered successful only if it can reproduce both the training data and the validation data.

The rationale for this training + cross-validation strategy operates on two levels. Conceptually, they reflect a hierarchy in model evaluation. A minimal functional model must at least reproduce the training data, which are essential metrics that define F_1_-ATPase as a rotary molecular motor. Yet, there may be, and, as demonstrated in Results, there indeed are, multiple mechanistic scenarios on a microscopic level by which the similar rotary catalysis behavior on the macroscopic level can be achieved. Cross-validation is therefore performed to test which specific underlying mechanistic scenarios are consistent with the further experimental observations.

Practically, this strategy aligns with the quantitative-ness of the data and the requirements of Bayesian parameter inference. The training data, being quantitative and often associated with well-defined experimental uncertainties, are readily incorporated into a statistically rigorous likelihood function (Equation (15) and Supplementary Table 7), making them suitable for parameter inference. In contrast, for the qualitative or semi-quantitative validation data, formulating a precise likelihood term would be challenging and potentially introduce subjective assumptions.

We choose this training + cross-validation strategy over a formal, single-step Bayesian model comparison method (e.g., evaluating the marginal likelihood) for both pragmatic and scientific reasons. Pragmatically, an accurate estimation of the marginal likelihood requires extensive sampling of the high-dimensional posterior distribution, which is computationally infeasible for our model due to the high cost of evaluating the likelihood function for each parameter set.

More importantly, cross-validation is a scientifically necessary step to rigorously test a model’s ability to generalize beyond simply fitting the data it is trained on and to validate its underlying physical assumptions. While Bayesian inference has a built-in mechanism to penalize model complexity (the Bayesian Occam’s Razor), this mechanism operates under the assumption that the model framework is a reasonable representation of the data-generating process. If a model’s foundational assumptions are flawed, a full Bayesian analysis might still yield a high-posterior model that has poor predictive power on new data — a form of model misspecification. Therefore, an external check like cross-validation remains a crucial step to ensure the robustness and physical realism of the model.

### Bayesian training approach

The aim of our Bayesian training approach is to sample an ensemble of parameter sets (Table 5) that adequately reproduce the training data (Table 6). According to the Bayes theorem, the posterior probability, *P* (**Ω**|**Ψ**^exp^), i.e., the probability of a parameter set **Ω** given the training data **Ψ**^exp^, is proportional to the likelihood *P* (**Ψ**^exp^|**Ω**) and the prior probability *P* (**Ω**):

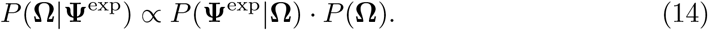

Here, the likelihood quantifies probability of the training data given the parameters, while the prior defines our initial assumptions about the parameters. Assuming statistical independence of all data points in **Ψ**^exp^ and of all parameters, we define the likelihood function *P* (**Ψ**^exp^|**Ω**) as the product of contributions from the three datasets (turnover rates 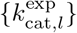, chemo-mechanical coupling efficiencies 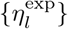, and nucleotide occupancies 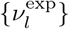), assuming statistical independence of all data points:

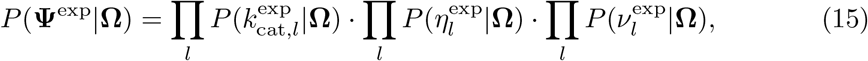

where the product index *l* runs over all data points within each respective dataset; the numbers of data points in the three datasets may differ. Supplementary Table 7 details the specific distribution assumed for each component. Similarly, we define the prior distribution *P* (**Ω**) as the product of the prior distributions of each individual parameter:

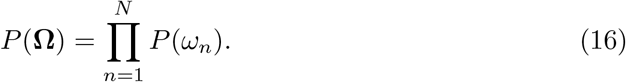

We assume a uniform distribution for each individual prior distributions *P* (*ω*_*n*_) (detailed in Supplementary Table 1). This choice is equivalent to assuming log-uniform priors for the transition rates that are determined by these parameters (free energies and free energy barriers) via the Boltzmann factor, ensuring that no a priori bias towards any particular timescale is imposed for a transition rate. The ranges for our uniform priors of the parameters (Supplementary Table 1) were chosen to restrain the resulting rates to biophysically plausible limits, e.g., by precluding conformational transitions faster than nanosecond timescale.

Critically, our goal is not to identify a single maximum a posteriori (MAP) estimate of the parameters. Instead, acknowledging the potential complexity of the posterior landscape which might contain multiple local maxima, we aim to sample an ensemble of high-posterior parameter sets that broadly covers these distinct regions. To this end, we perform hundreds of independent optimization runs, each initiated from a different, randomly chosen starting point. A stochastic hill-climbing search algorithm is employed for each individual optimization run, where the parameter set **Ω** stochastically explores the parameter space, each step increasing the posterior, until a maximum is reached. At each step within an optimization run, denoting the current parameter set as **Ω**^now^, a number *N*_try_ of guesses {**Ω**^try^} are sampled, where each individual parameter 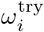 is drawn from a Gaussian distribution centered at 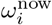 with standard deviation *σ*_try_. When some **Ω**^try^ are found to give higher posteriors than **Ω**^now^, the parameter set **Ω** then moves to the **Ω**^try^ of the highest posterior. Otherwise, the parameter set **Ω** stays at **Ω**^now^, and *σ*_try_ is tuned down by a factor *ϵ* (0 *< ϵ <* 1) to allow searching within a smaller area around **Ω**^now^, which is likely near a local maximum.

Finally, all resulting parameter sets that adequately reproduce the training data are collected into an ensemble (trained parameter sets) and carried forward for the subsequent cross-validation procedure.

### Evaluation of nucleotide titration curves and derivation of apparent binding affinities

Here we detail the calculation and analysis of nucleotide titration curves, which are crucial for evaluating our model against experimental data (Results). The nucleotide titration curve represents the steady-state total nucleotide occupancy *ν* (Equation (11)) as a function of nucleotide concentration. Mimicking experimental protocols for ATP titration curves (where [ADP] is kept low while [ATP] is varied from nanomolar to millimolar ranges) [21], we calculated the ATP titration curve numerically as part of the training procedure (Results). This calculation involved repeatedly solving for the steady-state distribution **ρ**^st^ (Equation (9)) while varying [ATP] from 100 nM to 1 mM (assuming a constant [ADP] of 1 nM), and then evaluating the total occupancy *ν* (Equation (11)). Similarly, an ADP titration curve was evaluated by varying [ADP] over the same range (assuming a constant [ATP] of 1 nM).

To quantitatively assess the agreement between our model-predicted ADP titration curves and the experimentally measured one [21], we utilize the apparent ADP binding affinities derived from fitting both curves to an equilibrium binding model [20, 21, 22]:

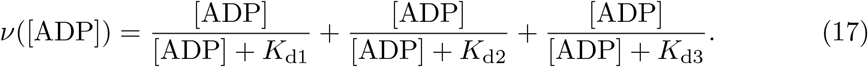

The three fitting parameters, *K*_d1,2,3_, resulting from this analysis are interpreted as (apparent) ADP binding affinities (dissociation constants) of the three β-subunits. They quantitatively characterize of the shape of the titration curve, as their relative magnitudes determine the number of distinct binding steps identifiable therein. The experimentally determined values exhibit distinct magnitudes (*K*_d1_ = 41 nM, *K*_d2_ = µM, *K*_d3_ = 42 µM) [21], providing the specific benchmark against which our model predictions are compared (Results).

Notably, these apparent binding affinities *K*_*d*1,2,3_ obtained from fitting (Equation (17)) are emergent, macroscopic properties reflecting the ensemble behaviors of the three differentiated β-subunits. Given the physical picture underlying our model where each β-subunit dynamically transitions between multiple conformations, an apparent affinity *K*_d*k*_ does not necessarily correspond to the intrinsic thermodynamic property of a single, static conformation. The latter, which we term a microscopic binding affinity, is instead directly related to a specific conformation 𝒞 and defined via the standard thermodynamic relation

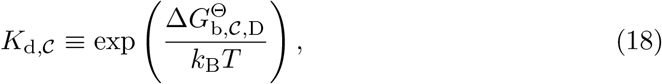

where 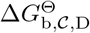 represents the standard ADP binding free energy of conformation 𝒞, one of our model parameters (Table 5).

We establish a theoretical link between the apparent affinities *K*_*d*1,2,3_ and the microscopic affinities {*K*_d, 𝒞_}, via an analytical expression derived in Supplementary Note 5. This analytical expression provides the theoretical foundation for our understanding of the incompatibility of the *ohc*-*w* and *ohc*-*m* variants with the experimentally determined apparent ADP binding affinities (Results). Below, we outline the key steps of this derivation.

First, we derive an approximate analytical solution for the steady-state distribution **ρ**^st^ (Supplementary Equations (48)–(53)). Under the low [ATP] when evaluating an ADP titration curve, the turnover rate is approximately zero, and the system approaches equilibrium (equilibrium approximation). Thus, the steady-state distribution approximates a Boltzmann distribution, which can be analytically expressed in terms of the underlying free energies of the respective Markov states (Equation (2)). These state free energies, in turn, depend on our model parameters, our model parameters (specifically, the conformational free energies 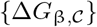 and the standard ADP binding free energies 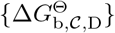, Table 5). Using this analytical expression for the steady-state distribution, the ADP titration curve *ν*([ADP]), and subsequently the apparent ADP binding affinities derived from it, can also be analytically expressed in terms of these model parameters (Supplementary Equations (55)–(59)).

Particularly, for model variants where the three β-subunits are decoupled (i.e., 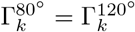 for at least two β-subunits, see Supplementary Note 5 for explanation), e.g., variants *ohc*-*w*/*s*, the analytical expressions for the apparent ADP binding affinities are rather straightforward. Each apparent ADP binding affinity *K*_d*k*_ is expressed as a superposition of the microscopic binding affinities 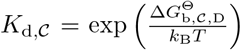 of the several accessible conformations,

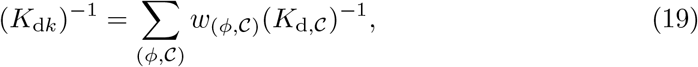

where the summation is over all combinations of *ϕ* and 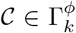; the weights {*w*_(*ϕ*, 𝒞)_} depend on the conformational free energies {*G*_β,𝒞_} of the accessible conformations (see Supplementary Equations (77),(78)).

### Physical constraints on γ-β restrictions

To systematically reduce the vast number of potential model variants (∼600 for *ohc*-variants and ∼14,000 for *ohc*_1_*c*_2_-variants), we derive several necessary conditions of γ-β restrictions for our model to be compatible with the consensus chemo-mechanical coupling scheme (Table 6 and Supplementary Fig. 2), as summarized in Supplementary Table 4 and detailed in Supplementary Note 3. These conditions are the technical implementation of the fundamental binding-change principle [16, 17, 6, 15, 35, 68, 73], complemented by our finding that a nucleotide-bound β-subunit energetically favors the catalytically active *c* conformation (both discussed in Results). For example, for ATP hydrolysis to occur in a β-subunit at +200° after ATP binding, the γ-β restrictions must allow this β-subunit to adopt the catalytically active *c* conformation at +200°, but then enforce its transition into an inactive conformation (*h* or *o*) upon rotation to +240° (Supplementary Equations (8), (9)). If *c* remained accessible at +240°, this energetically favored closed state would likely persist, potentially stalling the cycle or preventing subsequent steps, e.g., product dissociation preparation, which require less closed conformations. Applying this allow-and-enforce logic to all major catalytic steps yields the set of conditions summarized in Supplementary Table 4.

Note that all Family B variants (Supplementary Equation (7) in Supplementary Table 4) are further excluded as reasoned in Results. To further reduce the choices of plausible *ohc*_1_*c*_2_-variants, we introduce several additional constraints, considering the resolved structures of F_1_-ATPase, the physical parsimony of our aimed minimal model, and the intrinsic structural asymmetry of the γ-subunit (Supplementary Table 5). More detailed explanations for these constraints are provided in Supplementary Notes 3&4.

### Kinetic Monte-Carlo simulations of F_1_-ATPase coupled to external load

Kinetic Monte-Carlo (KMC) simulations mimicking the single-molecule experiments on F_1_-ATPase are presented in Results (see particularly Fig. 4a). In these simulations, the rotation of the *γ*-subunit and the circular motion of the probe centroid are coupled by a harmonic potential with force constant *κ*:

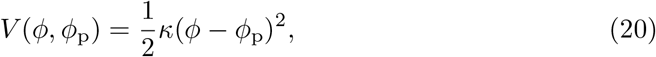

where *ϕ* and *ϕ*_p_ are the angular positions of the γ-subunit and the probe centroid, respectively. Further, the probe is subjected to viscous friction of the solution with rotational friction drag coefficient *ξ*. We estimate the spring force constant *κ* ≈ 5 *k*_B_*T* and the rotational friction drag coefficient *ξ* ≈ 6.7 × 10^−5^ *k*_B_*T ·* s for a 20-nm colloidal gold bead [8]. *κ* is estimated from the widths of the distributions of the recorded probe positions. *ξ* corresponds to the midpoint of 8*πµr*^3^ ≤ *ξ* ≤ 14*πµr*^3^ estimated in Ref. [8] (bead radius *r* = 20 nm, water viscosity *µ* = 10^−9^ pN · nm^−2^ s).

Given the estimated values of *κ* and *ξ*, the inertia of the probe is negligible compared to the viscous friction and force from the spring. Therefore, we use the overdamped Langevin equation to describe the circular motions of the probe:

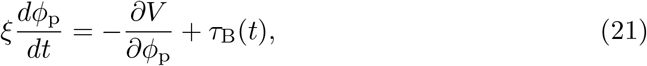

where *τ*_B_(*t*) is a fluctuating Brownian torque. This torque has the statistical properties ⟨*τ*_B_(*t*) ⟩ = 0 and ⟨*τ*_B_(*t*)*τ*_B_(*t*^′^)⟩ = 2*ξk*_B_*Tδ*(*t − t*^′^), where *δ*(*t − t*^′^) is the Dirac delta function.

We simulate the coupled system by combining stochastic integration of the over-damped Langevin equation (Equation (21)) with Gillespie algorithm [88, 89] for our Markov model of F_1_-ATPase. Consider the *n*th step starting at time *t*_*n*_, when the probe is at *ϕ*_p_(*t*_*n*_) and F_1_-ATPase is in Markov state *s*_*n*_ = *i*, with γ-orientation *ϕ*_*n*_. We assume that F_1_-ATPase remains in this state *s*_*n*_ during this step, which lasts until time *t*_*n*+1_ when the next step initiates and F_1_-ATPase jumps to a new state *s*_*n*+1_ instantaneously. During this time interval (*t*_*n*_ to *t*_*n*+1_), the γ-subunit remains at *ϕ*_*n*_, while the probe moves according to Equation (21). The probe position at the end of the interval, *ϕ*_p_(*t*_*n*+1_), is calculated via stochastic integration using the Euler-Maruyama method:

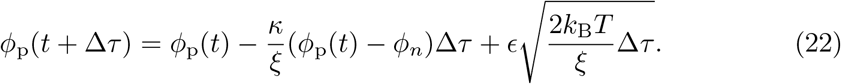

Here, *ϵ* is a random number drawn from the standard normal distribution. This equation is iteratively evaluated with a small time step Δ*τ* from *t* = *t*_*n*_ until *t*_*n*+1_. We use Δ*τ* = 10^−9^ s, which is at least one magnitude smaller than the timescale of the fastest transition in our Markov model.

The next state *s*_*n*+1_ of F_1_-ATPase and the holding time *τ* = *t*_*n*+1_ − *t*_*n*_ are determined by Gillespie algorithm [88, 89]. The next state *s*_*n*+1_ is chosen from the set of states {*j*} accessible from the current state *i*, with probability 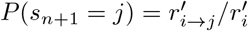, where 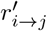 is the transition rate from *i* to *j* for F_1_-ATPase under load, and 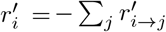 is the total rate of leaving state *i. τ* is sampled from the exponential distribution 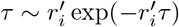. Crucially, these transition rates 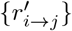 differ from the rates {*r*_*i*→*j*_} defined earlier (Supplementary Equations (1)–(5)) for unloaded F_1_-ATPase, because the harmonic potential coupling the γ-subunit and the probe modifies the free energy landscape. For transitions where the γ-subunit does not rotate, the landscape is unaffected, and thus 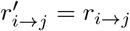.

In contrast, for a transition involving *γ*-subunit rotation by Δ*ϕ*, the harmonic potential *V* (*ϕ, ϕ*_p_) alters the free energy profile along the rotation coordinate *ϕ*. Taking the initial state *i* (at *ϕ*_*n*_) as the reference, the free energy profile becomes

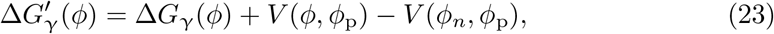

where Δ*G*_γ_(*ϕ*) is the profile for unloaded F_1_-ATPase. Assuming the transition state is located symmetrically between the initial and final states, i.e., at *ϕ*^∗^ = *ϕ*_*n*_ + *λ*Δ*ϕ* with *λ* = 0.5, the activation energy 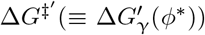 relative to the initial state is modified from the unloaded barrier Δ*G*^‡^ according to

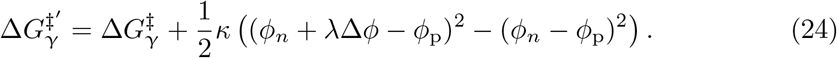

Consequently, the transition rate under load becomes:

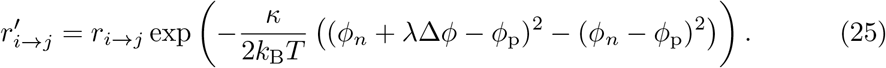

### Hidden Markov analysis of the simulated single-molecule trajectories

To extract the dwell positions and kinetics (Fig. 4e-g) from the simulated kinetic Monte-Carlo trajectories, we employ hidden Markov analysis (HMA) [90]. The detailed mathematical formulation is provided in Supplementary Note 6, while an overview is presented below.

The goal of HMA is to infer the most likely sequence of underlying discrete dwell states from the simulated trajectory of the probe’s angular position *ϕ*_*p*_, and subsequently to obtain the lifetime distributions for each dwell. Our HMA assumes the system transitions between a small number of discrete dwell states following Markovian dynamics, where transitions are restricted to adjacent dwell states (Supplementary Equations (90),(91)). The observed probe position *ϕ*_*p*_ at any time is assumed to follow a von Mises distribution centered around the true dwell angle (Supplementary Equation (92)).

We first use a modified Baum-Welch algorithm, adapted for the three-fold rotational symmetry of F_1_-ATPase (Supplementary Equations (94)–(102)), to estimate the HMA parameters, including the true dwell angles, the angular variances, and the transition probabilities. After obtaining the maximum-likelihood parameters, we use the Viterbi algorithm to infer the most probable sequence of dwell states from the simulated trajectory (Supplementary Equations (103),(104)). Based on this inferred sequence, the trajectory is segmented to generate the dwell lifetime distributions shown in Fig. 4e–g. These distributions are then fitted by a single exponential function *f* (*t*) ∝ exp(−*rt*) or a double exponential function *f* (*t*) ∝ exp(−*r*_1_*t*) − exp(− *r*_2_*t*) to determine the number of rate limiting steps and their rate constants.

## Data Availability

Source Data are provided with this paper. The previously published atomic coordinates of F_1_-ATPase referred to in this study are available in the Protein Data Bank (PDB) under accession code 1BMF [https://doi.org/10.2210/pdb1BMF/pdb].

## Code Availability

The custom code developed in this study is publicly available on the Github repository YixinChen95/MarkovianF1 (https://github.com/YixinChen95/MarkovianF1), and archived at Zenodo (https://doi.org/10.5281/zenodo.19133448). The repository contains the C and Python scripts organized into four modules for Bayesian training of the Markov model, evaluation of model predictions, kinetic Monte-Carlo simulations, and hidden Markov analysis of the simulated trajectories, respectively. A README file provides instructions for installation and execution, and includes example workflows demonstrating the main functions of the code.

## Acknowledgements

We thank Malte Schäffner and Daniel Szöllösi for helpful discussions, and Lars Bock and Petra Kellers for proofreading the manuscript.

## Author contributions

H.G. conceived the project. Y.C. conducted the research. Y.C. and H.G. interpreted the results and wrote the paper.

## References

[1] Walker, J. E. The ATP synthase: the understood, the uncertain and the unknown. Biochemical Society Transactions 41, 1–16 (2013).

[2] Junge, W. & Nelson, N. ATP synthase. Annual Review of Biochemistry 84, 631–657 (2015).

[3] Kühlbrandt, W. Structure and mechanisms of f-type ATP synthases. Annual Review of Biochemistry 88, 515–549 (2019).

[4] Abrahams, J. P., Leslie, A. G., Lutter, R. & Walker, J. E. Structure at 2.8 a resolution of F1-ATPase from bovine heart mitochondria. Nature 370, 621–628 (1994).

[5] Menz, R. I., Walker, J. E. & Leslie, A. G. Structure of bovine mitochondrial F1-ATPase with nucleotide bound to all three catalytic sites: implications for the mechanism of rotary catalysis. Cell 106, 331–341 (2001).

[6] Boyer, P. D. The ATP synthase—a splendid molecular machine. Annual Review of Biochemistry 66, 717–749 (1997).

[7] Nakamoto, R. K., Ketchum, C. J. & Al-Shawi, M. K. Rotational coupling in the f0f1 ATP synthase. Annual Review of Biophysics and Biomolecular Structure 28, 205–234 (1999).

[8] Yasuda, R., Noji, H., Yoshida, M., Kinosita Jr, K. & Itoh, H. Resolution of distinct rotational substeps by submillisecond kinetic analysis of F1-ATPase. Nature 410, 898–904 (2001).

[9] Itoh, H. et al. Mechanically driven ATP synthesis by F1-ATPase. Nature 427, 465–468 (2004).

[10] Kinosita, K., Yasuda, R., Noji, H. & Adachi, K. A rotary molecular motor that can work at near 100% efficiency. Philosophical Transactions of the Royal Society of London. Series B: Biological Sciences 355, 473–489 (2000).

[11] Sabbert, D., Engelbrecht, S. & Junge, W. Intersubunit rotation in active f-ATPase. Nature 381, 623–625 (1996).

[12] Häsler, K., Engelbrecht, S. & Junge, W. Three-stepped rotation of subunits γ and ε in single molecules of f-ATPase as revealed by polarized, confocal fluorometry. FEBS Letters 426, 301–304 (1998).

[13] Noji, H., Yasuda, R., Yoshida, M. & Kinosita Jr, K. Direct observation of the rotation of F1-ATPase. Nature 386, 299–302 (1997).

[14] Yasuda, R., Noji, H., Kinosita, K. & Yoshida, M. F1-ATPase is a highly efficient molecular motor that rotates with discrete 120 steps. Cell 93, 1117–1124 (1998).

[15] Adachi, K. et al. Coupling of rotation and catalysis in F1-ATPase revealed by single-molecule imaging and manipulation. Cell 130, 309–321 (2007).

[16] Cross, R. L. The mechanism and regulation of ATP synthesis by F1-atpases. Annual Review of Biochemistry 50, 681–714 (1981).

[17] Boyer, P. D. The binding change mechanism for ATP synthase—some probabilities and possibilities. Biochimica et Biophysica Acta (BBA) - Bioenergetics 1140, 215–250 (1993).

[18] Boyer, P. D. Energy, life, and ATP (nobel lecture). Angewandte Chemie International Edition 37, 2296–2307 (1998).

[19] Weber, J., Wilke-Mounts, S., Lee, R. S., Grell, E. & Senior, A. E. Specific placement of tryptophan in the catalytic sites of Escherichia coli F1-ATPase provides a direct probe of nucleotide binding: maximal ATP hydrolysis occurs with three sites occupied. Journal of Biological Chemistry 268, 20126–20133 (1993).

[20] Weber, J., Bowman, C. & Senior, A. E. Specific tryptophan substitution in catalytic sites of Escherichia coli F1-ATPase allows differentiation between bound substrate ATP and product ADP in steady-state catalysis. Journal of Biological Chemistry 271, 18711–18718 (1996).

[21] Mao, H. Z., Gray, W. D. & Weber, J. Does F1-ATPase have a catalytic site that preferentially binds MgADP? FEBS Letters 580, 4131–4135 (2006).

[22] Li, Y., Valdez, N. A., Mnatsakanyan, N. & Weber, J. The nucleotide binding affinities of two critical conformations of Escherichia coli ATP synthase. Archives of biochemistry and biophysics 707, 108899 (2021).

[23] Minagawa, Y. et al. Basic properties of rotary dynamics of the molecular motor Enterococcus hirae V1-ATPase. Journal of Biological Chemistry 288, 32700– 32707 (2013).

[24] Iino, R., Minagawa, Y., Ueno, H., Hara, M. & Murata, T. Molecular structure and rotary dynamics of Enterococcus hirae V1-ATP ase. IUBMB Life 66, 624–630 (2014).

[25] Ueno, H., Suzuki, K. & Murata, T. Structure and dynamics of rotary v1 motor. Cellular and Molecular Life Sciences 75, 1789–1802 (2018).

[26] Stewart, A. G., Sobti, M., Harvey, R. P. & Stock, D. Rotary atpases: models, machine elements and technical specifications. Bioarchitecture 3, 2–12 (2013).

[27] Stewart, A. G., Laming, E. M., Sobti, M. & Stock, D. Rotary atpases—dynamic molecular machines. Current opinion in structural biology 25, 40–48 (2014).

[28] Muench, S. P., Trinick, J. & Harrison, M. A. Structural divergence of the rotary atpases. Quarterly reviews of biophysics 44, 311–356 (2011).

[29] Arai, S., Maruyama, S., Shiroishi, M., Yamato, I. & Murata, T. An affinity change model to elucidate the rotation mechanism of V1-ATPase. Biochemical and Biophysical Research Communications 533, 1413–1418 (2020).

[30] Yamato, I., Kakinuma, Y. & Murata, T. Operating principles of rotary molecular motors: differences between F1 and V1 motors. Biophysics and Physicobiology 13, 37–44 (2016).

[31] Mao, H. Z. & Weber, J. Identification of the βtp site in the x-ray structure of F1-ATPase as the high-affinity catalytic site. Proceedings of the National Academy of Sciences 104, 18478–18483 (2007).

[32] Sobti, M., Ueno, H., Noji, H. & Stewart, A. G. The six steps of the complete F1-ATPase rotary catalytic cycle. Nature Communications 12, 4690 (2021).

[33] Nishizaka, T. et al. Chemomechanical coupling in F1-ATPase revealed by simultaneous observation of nucleotide kinetics and rotation. Nature Structural & Molecular Biology 11, 142–148 (2004).

[34] Ariga, T., Muneyuki, E. & Yoshida, M. F1-ATPase rotates by an asymmetric, sequential mechanism using all three catalytic subunits. Nature Structural & Molecular Biology 14, 841–846 (2007).

[35] Adachi, K., Oiwa, K., Yoshida, M., Nishizaka, T. & Kinosita Jr, K. Controlled rotation of the F1-ATPase reveals differential and continuous binding changes for ATP synthesis. Nature Communications 3, 1022 (2012).

[36] Pu, J. & Karplus, M. How subunit coupling produces the γ-subunit rotary motion in F1-ATPase. Proceedings of the National Academy of Sciences 105, 1192–1197 (2008).

[37] Milgrom, M. Y., Murataliev, B. M. & Boyer, D. P. Bi-site activation occurs with the native and nucleotide-depleted mitochondrial F1-ATPase. Biochemical Journal 330, 1037–1043 (1998).

[38] Bulygin, V. V. & Milgrom, Y. M. A bi-site mechanism for Escherichia coli F1-ATPase accounts for the observed positive catalytic cooperativity. Biochimica et Biophysica Acta (BBA) - Bioenergetics 1787, 1016–1023 (2009).

[39] Milgrom, Y. M. Characteristics of protection by MgADP and MgATP of α 3 β 3 γ subcomplex of thermophilic bacillus ps3 βy341w-mutant f 1-ATPase from inhibition by 7-chloro-4-nitrobenz-2-oxa-1, 3-diazole support a bi-site mechanism of catalysis. Biochemistry (Moscow) 76, 1253–1261 (2011).

[40] Weber, J. & Senior, A. E. Bi-site catalysis in F1-ATPase: does it exist? Journal of Biological Chemistry 276, 35422–35428 (2001).

[41] Wang, H. & Oster, G. Energy transduction in the F1 motor of ATP synthase. Nature 396, 279–282 (1998).

[42] Oster, G., Wang, H. & Grabe, M. How Fo–ATPase generates rotary torque. Philosophical Transactions of the Royal Society of London. Series B: Biological Sciences 355, 523–528 (2000).

[43] Wang, H. & Oster, G. Ratchets, power strokes, and molecular motors. Applied Physics A 75, 315–323 (2002).

[44] Ma, J. et al. A dynamic analysis of the rotation mechanism for conformational change in F1-ATPase. Structure 10, 921–931 (2002).

[45] Nam, K., Pu, J. & Karplus, M. Trapping the ATP binding state leads to a detailed understanding of the F1-ATPase mechanism. Proceedings of the National Academy of Sciences 111, 17851–17856 (2014).

[46] Böckmann, R. A. & Grubmüller, H. Conformational dynamics of the F1-ATPase β-subunit: a molecular dynamics study. Biophysical Journal 85, 1482–1491 (2003).

[47] Czub, J. & Grubmüller, H. Torsional elasticity and energetics of F1-ATPase. Proceedings of the National Academy of Sciences 108, 7408–7413 (2011).

[48] Czub, J., Wieczør, M., Tobiszewski, A. & Grubmüller, H. Rotation triggers nucleotide-independent conformational transition of the empty beta subunit of F1-ATPase. Biophysical Journal 106, 253a (2014).

[49] Czub, J., Wieczør, M., Prokopowicz, B. & Grubmüller, H. Mechanochemical energy transduction during the main rotary step in the synthesis cycle of F1-ATPase. Journal of the American Chemical Society 139, 4025–4034 (2017).

[50] Dittrich, M., Hayashi, S. & Schulten, K. On the mechanism of ATP hydrolysis in F1-ATPase. Biophysical Journal 85, 2253–2266 (2003).

[51] Dittrich, M., Hayashi, S. & Schulten, K. ATP hydrolysis in the β_T P_ and β_DP_catalytic sites of F1-ATPase. Biophysical Journal 87, 2954–2967 (2004).

[52] Sakaki, N. et al. One rotary mechanism for F1-ATPase over ATP concentrations from millimolar down to nanomolar. Biophysical Journal 88, 2047–2056 (2005).

[53] Iida, T. et al. Single-molecule analysis reveals rotational substeps and chemo-mechanical coupling scheme of Enterococcus hirae V1-ATPase. Journal of Biological Chemistry 294, 17017–17030 (2019).

[54] Shimabukuro, K. et al. Catalysis and rotation of F1 motor: cleavage of ATP at the catalytic site occurs in 1 ms before 40 substep rotation. Proceedings of the National Academy of Sciences 100, 14731–14736 (2003).

[55] Watanabe, R., Iino, R., Shimabukuro, K., Yoshida, M. & Noji, H. Temperature-sensitive reaction intermediate of F1-ATPase. EMBO reports 9, 84–90 (2008).

[56] Furuike, S. et al. Temperature dependence of the rotation and hydrolysis activities of F1-ATPase. Biophysical Journal 95, 761–770 (2008).

[57] Watanabe, R., Iino, R. & Noji, H. Phosphate release in F1-ATPase catalytic cycle follows ADP release. Nature Chemical Biology 6, 814–820 (2010).

[58] Böckmann, R. A. & Grubmüller, H. Nanoseconds molecular dynamics simulation of primary mechanical energy transfer steps in F1-ATP synthase. Nature Structural Biology 9, 198–202 (2002).

[59] Bilyard, T. et al. High-resolution single-molecule characterization of the enzymatic states in Escherichia coli F1-ATPase. Philosophical Transactions of the Royal Society B: Biological Sciences 368, 20120023 (2013).

[60] Spetzler, D. et al. Single molecule measurements of F1-ATPase reveal an inter-dependence between the power stroke and the dwell duration. Biochemistry 48, 7979–7985 (2009).

[61] Zarco-Zavala, M. et al. The 3 × 120 rotary mechanism of paracoccus denitrificans F1-ATPase is different from that of the bacterial and mitochondrial F1-atpases. Proceedings of the National Academy of Sciences 117, 29647–29657 (2020).

[62] Suzuki, T., Tanaka, K., Wakabayashi, C., Saita, E.-i. & Yoshida, M. Chemomechanical coupling of human mitochondrial F1-ATPase motor. Nature Chemical Biology 10, 930–936 (2014).

[63] Kobayashi, R., Ueno, H.Li, C.-B. & Noji, H. Rotary catalysis of bovine mitochondrial F1-ATPase studied by single-molecule experiments. Proceedings of the National Academy of Sciences 117, 1447–1456 (2020).

[64] Cingolani, G. & Duncan, T. M. Structure of the ATP synthase catalytic complex (F1) from Escherichia coli in an autoinhibited conformation. Nature Structural & Molecular Biology 18, 701–707 (2011).

[65] Nakano, A., Kishikawa, J.-i., Mitsuoka, K. & Yokoyama, K. Mechanism of ATP hydrolysis dependent rotation of bacterial ATP synthase. Nature Communications 14, 4090 (2023).

[66] Weber, J. & Senior, A. E. Catalytic mechanism of F1-ATPase. Biochimica et Biophysica Acta (BBA) -Bioenergetics 1319, 19–58 (1997).

[67] Senior, A. E., Nadanaciva, S. & Weber, J. The molecular mechanism of ATP synthesis by f1f0-ATP synthase. Biochimica et Biophysica Acta (BBA) - Bioenergetics 1553, 188–211 (2002).

[68] Watanabe, R. et al. Mechanical modulation of catalytic power on F1-ATPase. Nature Chemical Biology 8, 86–92 (2012).

[69] Bowler, M. W., Montgomery, M. G., Leslie, A. G. & Walker, J. E. Ground state structure of F1-ATPase from bovine heart mitochondria at 1.9 å resolution. Journal of Biological Chemistry 282, 14238–14242 (2007).

[70] Sugawa, M. et al. F1-ATPase conformational cycle from simultaneous single-molecule fret and rotation measurements. Proceedings of the National Academy of Sciences 113, E2916–E2924 (2016).

[71] Nadanaciva, S., Weber, J. & Senior, A. E. New probes of the F1-ATPase catalytic transition state reveal that two of the three catalytic sites can assume a transition state conformation simultaneously. Biochemistry 39, 9583–9590 (2000).

[72] Baylis Scanlon, J. A., Al-Shawi, M. K., Le, N. P. & Nakamoto, R. K. Determination of the partial reactions of rotational catalysis in F1-ATPase. Biochemistry 46, 8785–8797 (2007).

[73] Volkán-Kacsó, S., Matute, R. A., Michel-Beyerle, M.-E., Khatchikian, O. & Marcus, R. A. Nucleotide exchange mechanism involving angle-dependent rate constants extracted from F1-ATPase single-molecule rotation trajectories. The Journal of Physical Chemistry B 130, 668–676 (2025).

[74] ichi Okazaki, K. & Hummer, G. Phosphate release coupled to rotary motion of f_1_-ATPase. Proceedings of the National Academy of Sciences 110, 16468–16473 (2013).

[75] Kinosita Jr, K., Adachi, K. & Itoh, H. Rotation of F1-ATPase: how an ATP-driven molecular machine may work. Annual Review of Biophysics and Biomolecular Structure 33, 245–268 (2004).

[76] Uchihashi, T., Iino, R., Ando, T. & Noji, H. High-speed atomic force microscopy reveals rotary catalysis of rotorless F1-ATPase. Science 333, 755–758 (2011).

[77] Furuike, S. et al. Axle-less F1-ATPase rotates in the correct direction. Science 319, 955–958 (2008).

[78] Sobti, M., Ueno, H., Brown, S. H., Noji, H. & Stewart, A. G. The series of conformational states adopted by rotorless F1-ATPase during its hydrolysis cycle. Structure 32, 393–399 (2024).

[79] Furlong, E. J. et al. The molecular structure of an axle-less F1-ATPase. Biochimica et Biophysica Acta (BBA) - Bioenergetics 1866, 149521 (2025).

[80] Kinosita, K., Yasuda, R., Noji, H. & Adachi, K. A rotary molecular motor that can work at near 100% efficiency. Philosophical Transactions of the Royal Society of London. Series B: Biological Sciences 355, 473–489 (2000).

[81] Böckmann, R. A. & Grubmüller, H. Nanoseconds molecular dynamics simulation of primary mechanical energy transfer steps in F1-ATP synthase. Nature Structural Biology 9, 198–202 (2002).

[82] Böckmann, R. A. & Grubmüller, H. Conformational dynamics of the F1-ATPase β-subunit: a molecular dynamics study. Biophysical Journal 85, 1482–1491 (2003).

[83] Yang, W., Gao, Y., Cui, Q., Ma, J. & Karplus, M. The missing link between thermodynamics and structure in F1-ATPase. Proceedings of the National Academy of Sciences 100, 874–879 (2003).

[84] Sielaff, H., Rennekamp, H., Engelbrecht, S. & Junge, W. Functional halt positions of rotary fof1-ATPase correlated with crystal structures. Biophysical Journal 95, 4979–4987 (2008).

[85] Okuno, D. et al. Correlation between the conformational states of F1-ATPase as determined from its crystal structure and single-molecule rotation. Proceedings of the National Academy of Sciences 105, 20722–20727 (2008).

[86] Yasuda, R. et al. The ATP-waiting conformation of rotating F1-ATPase revealed by single-pair fluorescence resonance energy transfer. Proceedings of the National Academy of Sciences 100, 9314–9318 (2003).

[87] Rosing, J. & Slater, E. C. The value of δg° for the hydrolysis of ATP. Biochimica et Biophysica Acta (BBA) - Bioenergetics 267, 275–290 (1972).

[88] Gillespie, D. T. A general method for numerically simulating the stochastic time evolution of coupled chemical reactions. Journal of Computational Physics 22, 403–434 (1976).

[89] Gillespie, D. T. Stochastic simulation of chemical kinetics. Annual Review of Physical Chemistry 58, 35–55 (2007).

[90] Stigler, J. & Rief, M. Hidden markov analysis of trajectories in single-molecule experiments and the effects of missed events. ChemPhysChem 13, 1079–1086 (2012).

