## Supplementary Materials for "A Minimal Chemo-mechanical Markov Model for Rotary Catalysis of F_1_-ATPase"

*For manuscript:*

### Supplementary Figures (referenced in the main article)

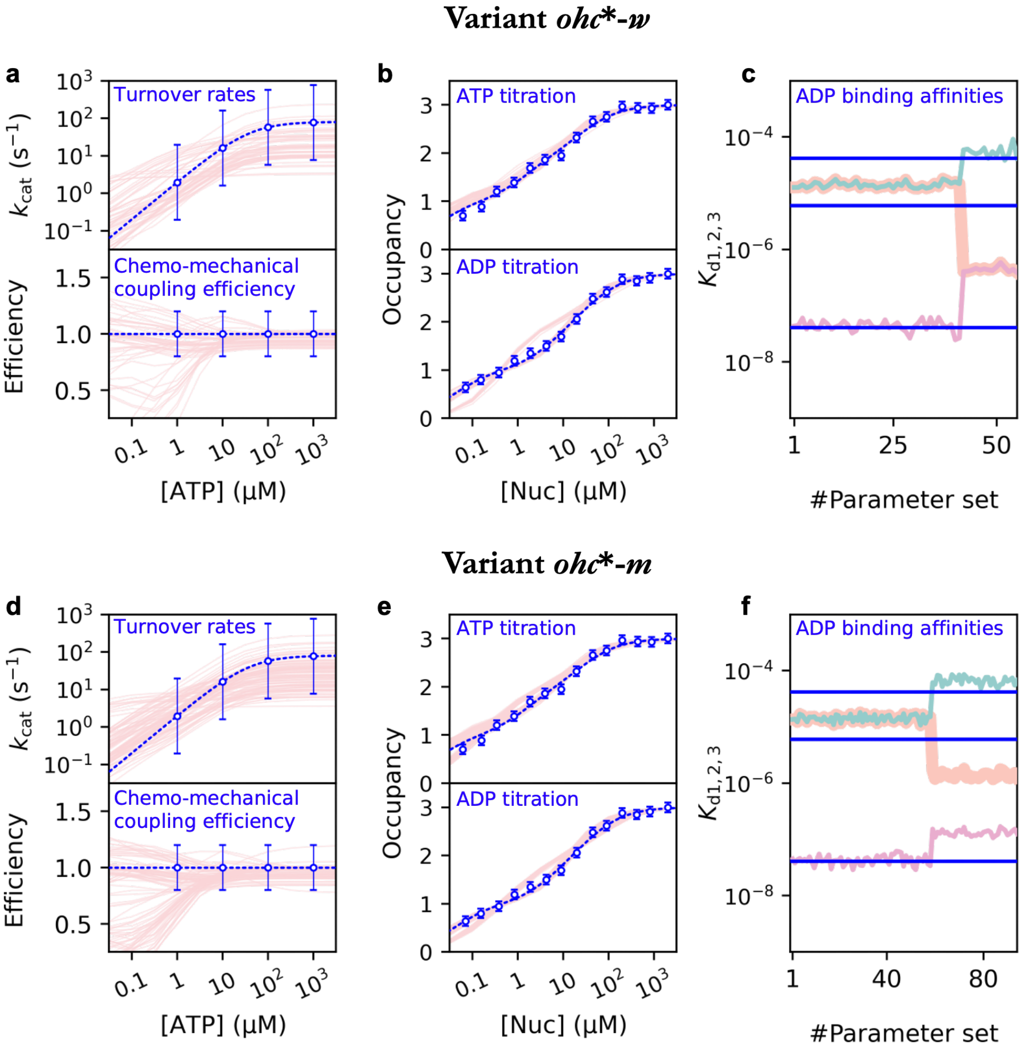


**Supplementary Fig. 1**: **Predicted properties of two *ohc*-variants (Table 2) against training data**

Predictions by the *ohc-w* (**a**–**c**) and *ohc-m* (**d**–**f**) variants, using the ensembles of parameter sets obtained from Bayesian training (Methods). The plots follow the same representation as in Fig. 2a–c of the main text.

**a**, **d** Turnover rates (upper panels) and chemo-mechanical coupling efficiencies (lower panels) as a function of [ATP].

**b**, **e** ATP (upper) and ADP (lower) titration curves showing the total nucleotide occupancy.

**c**, **f** Apparent ADP binding affinities ($K_{d1}, K_{d2}, K_{d3}$) derived from the ADP titration curves (Equation (17)). The blue horizontal lines indicate the experimental values [21].

Note that while both variants can reproduce the turnover rates and efficiency, they fail to reproduce the three distinct experimental binding affinities (which span orders of magnitude). Specifically, the *ohc-w* variant predicts two degenerate affinities (overlapping curves in **c**) due to the inherent symmetry of its γ-β restrictions, qualitatively contradicting the experimental data.

Source data are provided as a Source Data file.

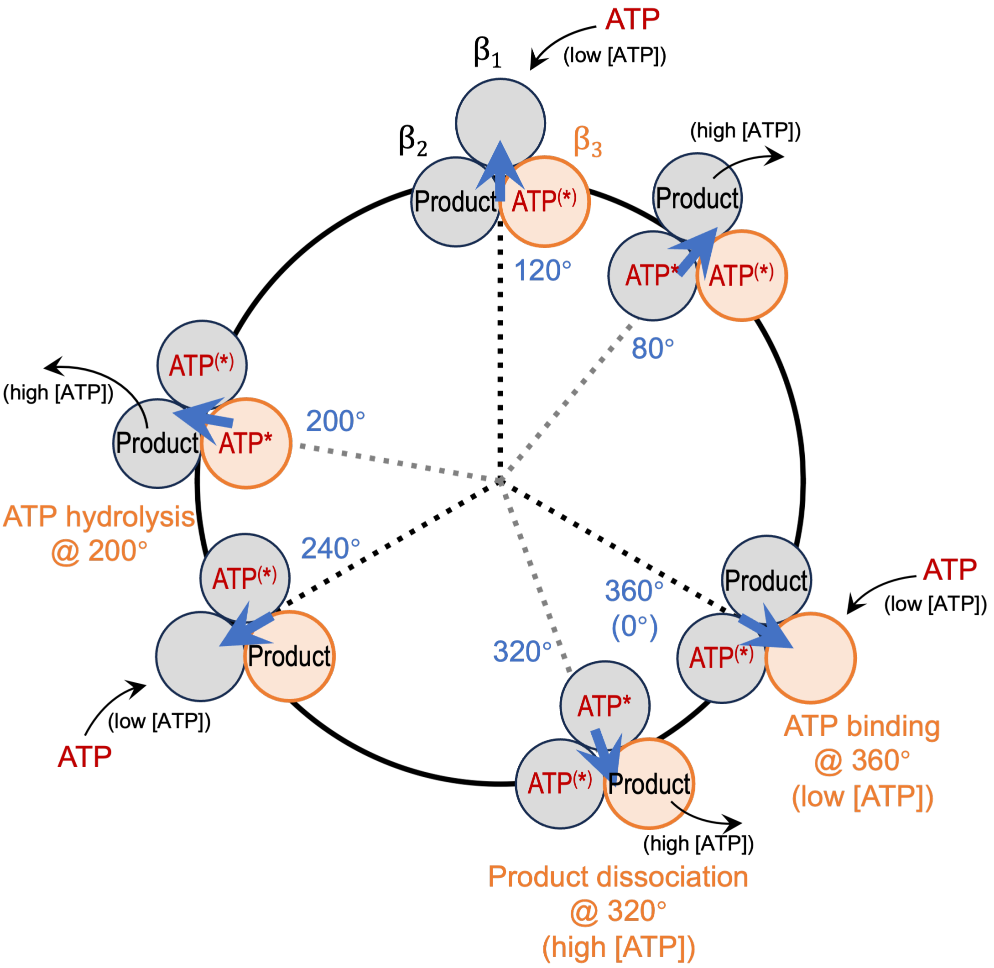


**Supplementary Fig. 2**: **Consensus chemo-mechanical coupling scheme**
This schematic summarizes the $\gamma$-orientations of the major catalytic steps for a single β-subunit (exemplified here by the $\beta_3$-subunit, highlighted in orange) relative to the rotation of the $\gamma$-subunit, as synthesized from a broad range of experimental studies [8, 15, 20–22, 32–34, 54–57, 65–67, 70, 75]. (i) At low ATP concentrations, ATP binding occurs at the ATP-waiting dwell (defined as 360°, or 0°). Relative to this binding angle, (ii) ATP hydrolysis occurs at +200°; (iii) At high ATP concentrations, product dissociation occurs at +320°. This step represents the effective rate-limiting dissociation of the two products (ADP and Pi).

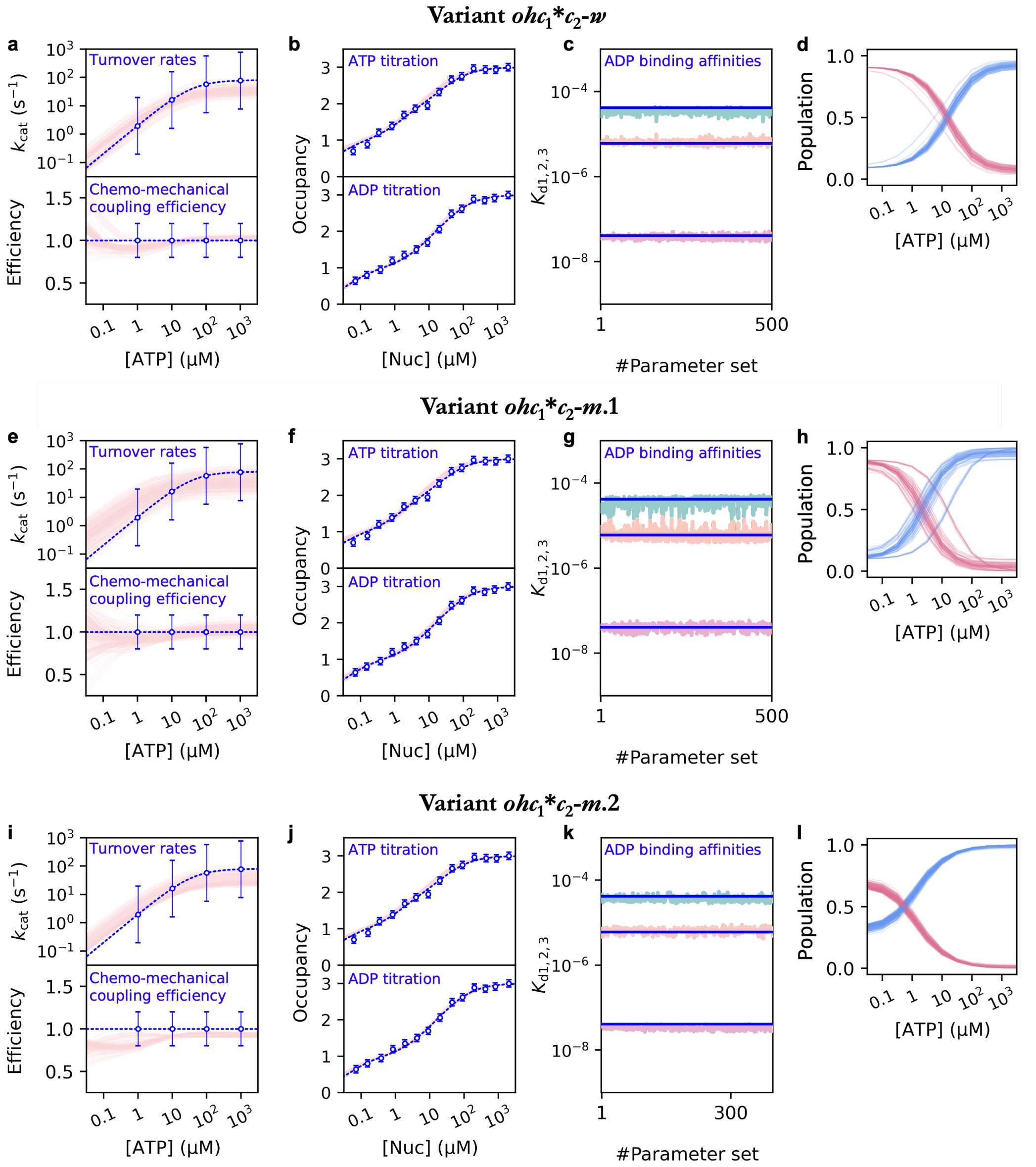


**Supplementary Fig. 3**: **Predicted properties of the three $ohc_1^*c_2$-variants (Table 3) against training data**

Predictions by the $ohc_1^*c_2$-$w$ (**a**–**d**), $ohc_1^*c_2$-$m.1$ (**e**–**h**), and $ohc_1^*c_2$-$m.2$ (**i**–**l**) variants, using the ensembles of parameter sets obtained from Bayesian training (Methods).

**a**–**c**, **e**–**g**, **i**–**k** Comparison against the training data (turnover rates, efficiency, titration curves, and apparent binding affinities), following the representation in Fig. 2a–c of the main text. Notably, unlike the $ohc$-variants (Supplementary Fig. 1), all three $ohc_1^*c_2$-variants successfully reproduce the three distinct experimental binding affinities (**c**, **g**, **k**).

**d**, **h**, **l** First cross-validation test: predicted steady-state populations of the 80°-dwells (blue curves) and 120°-dwells (pink curves) as a function of [ATP]. All three variants correctly predict the experimentally observed population shift from the ATP-waiting dwells (120°) at low [ATP] to the catalytic dwells (80°) as [ATP] increases from submicromolar to millimolar [8].

Source data are provided as a Source Data file.

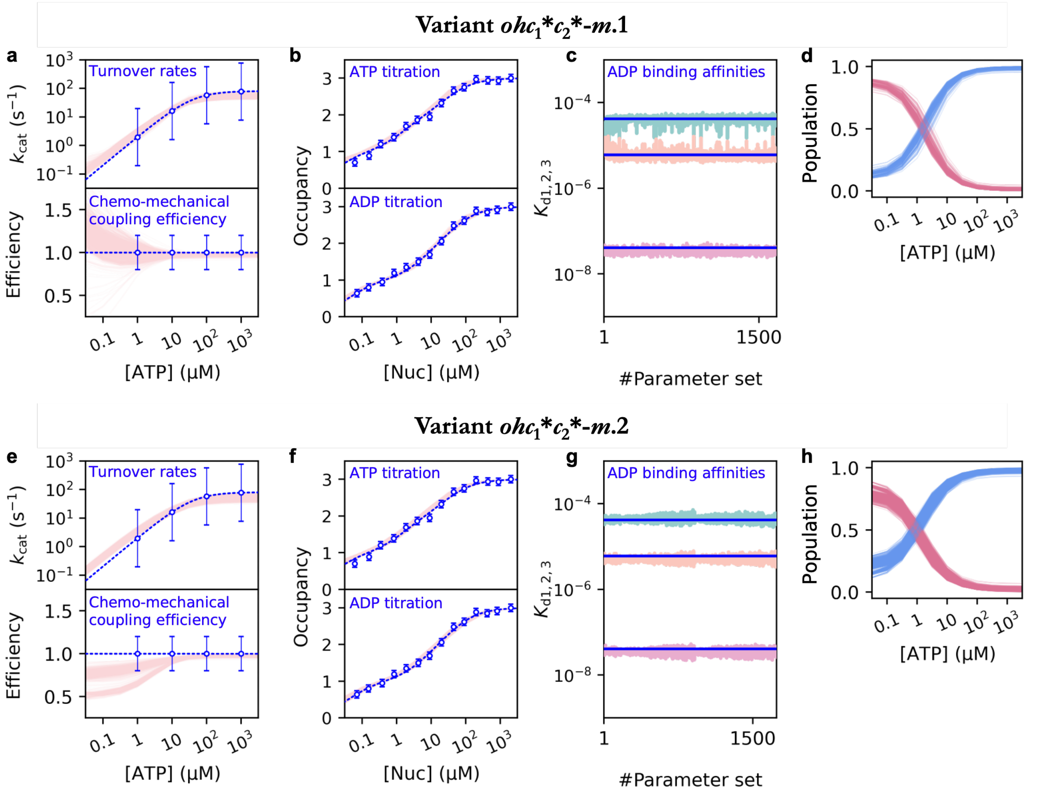


**Supplementary Fig. 4**: **Predicted properties of the two $ohc_1^*c_2^*$-variants (Table 3) against training data**

Predictions by the $ohc_1^*c_2^*$-$m$.1 (**a**–**d**) and $ohc_1^*c_2^*$-$m$.2 (**e**–**h**) variants, using the ensembles of parameter sets obtained from Bayesian training (Methods).

**a**–**c**, **e**–**g** Comparison against the training data (turnover rates, efficiency, titration curves, and apparent binding affinities), following the representation in Fig. 2a–c of the main text. Notably, unlike the *ohc*-variants (Supplementary Fig. 1), both $ohc_1^*c_2^*$-variants successfully reproduce the three distinct experimental binding affinities (**c**, **g**, **k**).

**d**, **h**, **l** First cross-validation test: predicted steady-state populations of the 80°-dwells (blue curves) and 120°-dwells (pink curves) as a function of [ATP]. Both variants correctly predict the experimentally observed population shift from the ATP-waiting dwells (120°) at low [ATP] to the catalytic dwells (80°) as [ATP] increases from submicromolar to millimolar [8].

Source data are provided as a Source Data file.

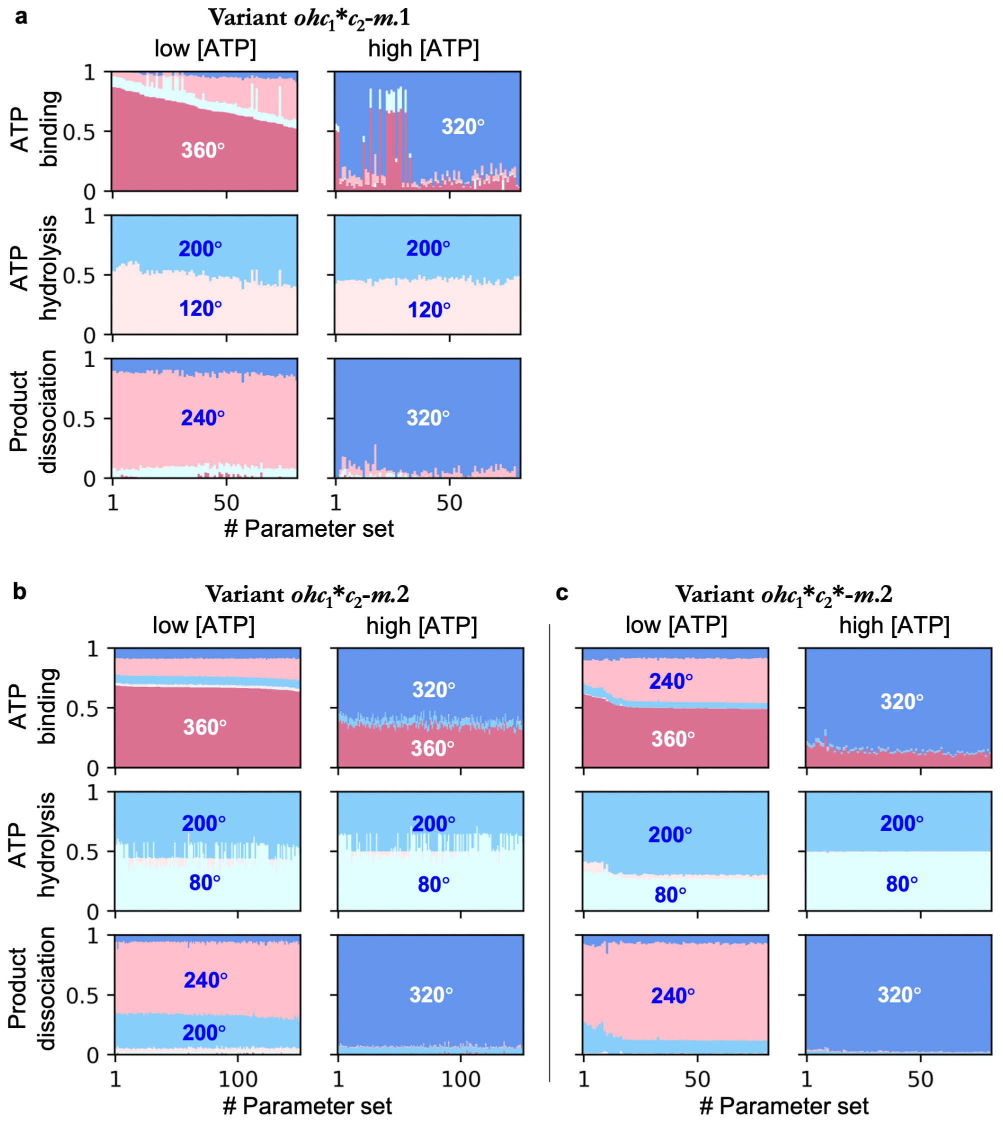


**Supplementary Fig. 5**: **Predicted chemo-mechanical coupling schemes of three** $ohc_1c_2$**-variants (Table 3)**

Second cross-validation test: predicted probability distributions of the major catalytic steps (ATP binding, ATP hydrolysis, and product dissociation) over different γ-subunit orientation for the $ohc_1^*c_2$-$m.1$ (**a**), $ohc_1^*c_2$-$m.2$ (**b**), and $ohc_1^*c_2^*$-$m.2$ (**c**) variants.

The plots follow the same representation as in Fig. 2e and Fig. 3a,b of the main text. For each variant, the probabilities are shown for low (0.1 μM, left columns) and high (1 mM, right columns) [ATP]. Note that all three variants predict an [ATP]-dependent shift in the γ-orientation of product dissociation (from ∼240° at low [ATP] to ∼320° at high [ATP]), consistent with "Scenario 2" discussed in the main text (Results). Crucially, all three variants correctly predict ATP hydrolysis at ∼200° and product dissociation under high [ATP] at ∼320°, thereby satisfying the criteria of the consensus chemo-mechanical coupling scheme (Table 6 and Supplementary Fig. 2).

Source data are provided as a Source Data file.

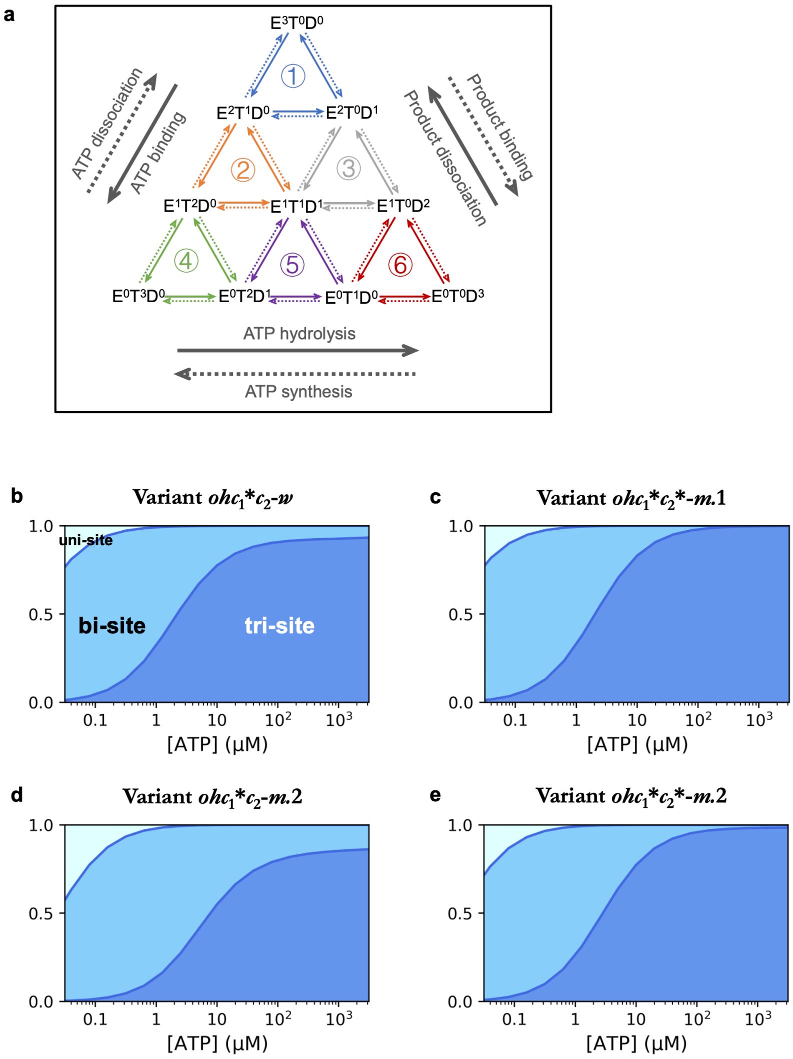


**Supplementary Fig. 6**: **Predicted contributions of uni-site, bi-site and tri-site pathways to the overall catalytic activity**

**a** Schematic of the catalytic pathways grouped by chemical composition. The Markov states are aggregated into 10 groups denoted as $\{E^xT^yD^z\}$, where $x$, $y$, $z$ represent the number of empty, ATP-bound, and product-bound catalytic sites, respectively ($x + y + z = 3$). Arrows indicate transitions between these groups: solid arrows for net ATP hydrolysis (counter-clockwise flow) and dotted arrows for net ATP synthesis (clockwise flow). Circled numbers ①–⑥ denote distinct catalytic cycles: Cycle ① represents the *uni-site* pathway (total nucleotide occupancy switches between 0 and 1); Cycles ② and ③ represent *bi-site* pathways (occupancy switches between 1 and 2); Cycles ④, ⑤, and ⑥ represent *tri-site* pathways (occupancy switches between 2 and 3).

**b**–**e** Stacked area plots showing the relative contributions of these pathways to the total turnover rate as a function of [ATP] for the four candidate variants: $ohc_1^*c_2$-$w$ (**b**), $ohc_1^*c_2^*$-$m.1$ (**c**), $ohc_1^*c_2$-$m.2$ (**d**), and $ohc_1^*c_2^*$-$m.2$ (**e**). Data are shown for a representative parameter set from the trained ensemble of each variant. Light blue: uni-site contribution; Medium blue: bi-site contribution; Dark blue: tri-site contribution.

Consistent across all variants, the model predicts a transition from bi-site-dominated catalysis (medium blue) at micromolar [ATP] to tri-site-dominated catalysis (dark blue) at physiological millimolar [ATP], reconciling the controversy regarding the stoichiometry of the catalytic cycle.

Source data are provided as a Source Data file.

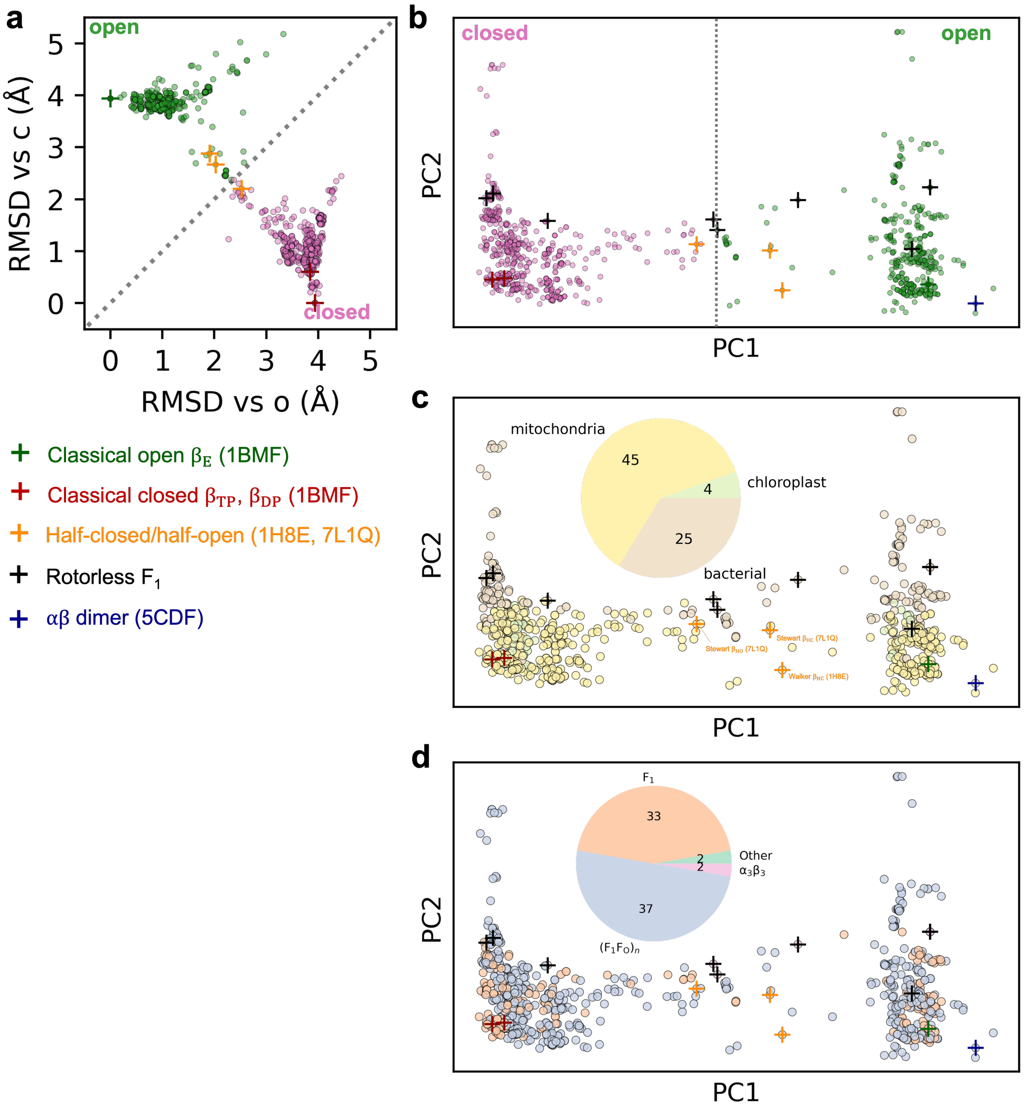


**Supplementary Fig. 7**: **Structural heterogeneity of β-subunits revealed by Principal Component Analysis (PCA) and RMSD clustering**

Analysis of 1077 β-subunit structures extracted from 331 PDB entries reported in 74 independent studies.

**a** RMSD scatter plot. Each dot represents a single β-subunit structure. The x-axis and y-axis show the $C_\alpha$-RMSD relative to the classical open ($\beta_E$, PDB: 1BMF) and closed ($\beta_{TP}$, PDB: 1BMF) reference structures, respectively. The structures fall into two primary clusters: "open-like" (green, $RMSD_o < RMSD_c$) and "closed-like" (pink, $RMSD_o > RMSD_c$).

**b–d** Projection of the 1077 structures onto the first two principal components (PC1 and PC2) obtained from a global PCA. Crosses (+) denote the positions of representative reference structures labeled in **a**.

**b** Structures colored by their RMSD classification (consistent with **a**). PC1 primarily captures the opening/closing motion, clearly separating open (right) and closed (left) states.

**c** Structures colored by source organism (mitochondria, chloroplast, bacteria).

**d** Structures colored by complex composition, including (1) an isolated β-subunit and a αβ dimer ("Other"), (2) several $\alpha_3\beta_3$ subcomplexes of $F_1$-ATPase ("$\alpha_3\beta_3$", also referred to as rotorless $F_1$), (3) several $\alpha_3\beta_3\gamma$ subcomplexes of $F_1$-ATPase and complete $F_1$-ATPase ("$F_1$"), as well as (4) monomers or multimers of $F_1F_O$-ATP synthase ("$(F_1F_O)_n$").

Source data are provided as a Source Data file.

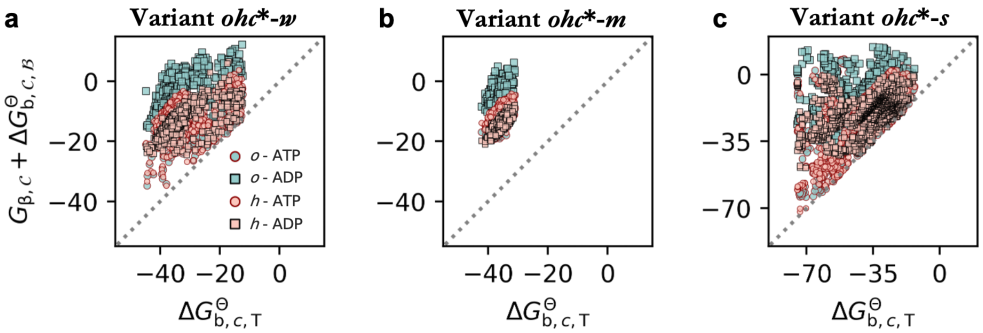


**Supplementary Fig. 8**: **Energetic preference for the closed conformation in nucleotide-bound β-subunits**
Scatter plots comparing the total free energies of a nucleotide-bound β-subunit in non-closed conformations $(o, h)$ against the closed conformation $(c)$.
**a**–**c** Results for the *ohc-w*, *ohc-m*, and *ohc-s* variants, respectively, derived from the ensembles of trained parameter sets.
The x-axis shows the total free energy of the ATP-bound closed state ($G_{\beta,c}+\Delta G^{\ominus}_{\mathrm{b},c,\mathrm{T}}$; the conformational free energy $G_{\beta,c}$ is set to 0, and is thus omitted from the label). The y-axis shows the total free energy ($G_{\beta,\mathcal{C}} + \Delta G^{\ominus}_{\mathrm{b},\mathcal{C},\mathcal{B}}$) for ATP-bound (circles) and ADP-bound (squares) subunits in the open ($o$, green) and half-closed ($h$, red) conformations. These free energies are in unit of $k_{\mathrm{B}}T$, assuming temperature $T = 298$ K. The dashed diagonal line indicates equi-energy ($y = x$).
The observation that virtually all data points lie above this diagonal demonstrates that, for a nucleotide-bound β-subunit, the catalytically active closed conformation $c$ is energetically more favorable than the open or half-closed conformations. As discussed in the main text (Discussion) and Supplementary Note 1, this energetic stability of the closed state is likely a necessary condition for the model to achieve high chemo-mechanical coupling efficiency.
Source data are provided as a Source Data file.

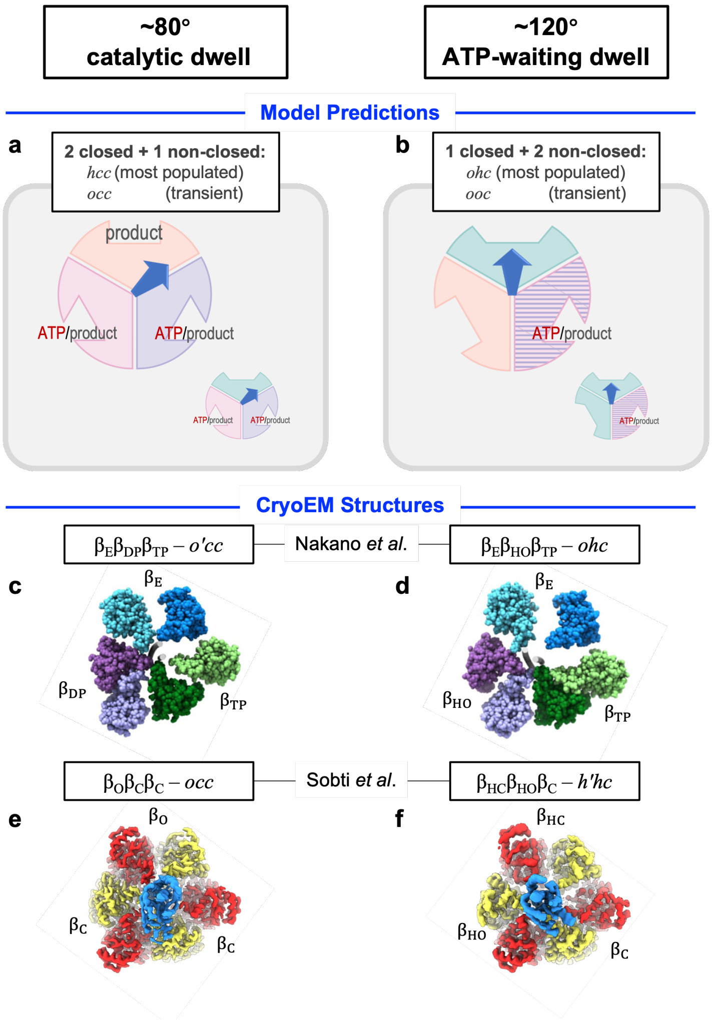


**Supplementary Fig. 9: Comparison between model-predicted functional stages and cryoEM structures**

**a**, **b** Major joint functional states of the three β-subunits (functional stages) within the 80°- and 120°-dwells predicted by the $ohc_1^* c_2^*$-$m.1$, $ohc_1^* c_2$-$m.2$, and $ohc_1^* c_2^*$-$m.2$ variants (Table 4). The schematics illustrate the most populated stages (large icons) and relevant transient or minor stages (small icons). **a** For the ∼80° catalytic dwell, the three variants predict an ensemble dominated by *hcc* and *occ* states ($c_1$, $c_2$ are collectively denoted as $c$), featuring a "2 closed + 1 non-closed" nomenclature compatible with single-molecule FRET data [70]. The *hcc* state is most populated, representing the stage where the $\beta_1$-subunit awaits product dissociation; the *occ* appears as a transient intermediate after product dissociation in the $\beta_1$-subunit but before ATP hydrolysis completes in the $\beta_2$-subunit. **b** For the ∼120° ATP-waiting dwell, the three variants predict an ensemble dominated by *ohc* and *ooc* states, featuring a "1 closed + 2 non-closed" nomenclature also compatible with single-molecule FRET data [70]. The *ohc* state is most populated, representing the stage where the empty $\beta_1$-subunit awaits substrate ATP binding. The *ooc* state appears when product dissociation in the $\beta_2$-subunit precedes ATP binding in the $\beta_1$-subunit, which is likely at low [ATP]. The [ATP]-dependent populations of these states are shown in Supplementary Figure 10.

**c**–**f** Re-interpretation of cryoEM structures based on model predictions. **c**, **d** Structures by Nakano *et al.* [65]: Although labeled "open" ($\beta_E$) in both dwells, the $\beta_1$-subunit (blue) appears structurally more compact at 80° than at 120°. This structural variance aligns with our model-predicted most populated states where the $\beta_1$-subunit transitions from the tighter *h* state to the looser *o* state upon the 80°→120° rotation. **e**, **f** Structures by Sobti *et al.* [32], where, in contrast, the $\beta_1$-subunit (upper) appears structurally less compact at 80° than at 120° (cf. $\beta_{HC}$ and $\beta_O$). Given additional experimental evidence [8, 15, 22, 64, 78, 79], we suggest the 80°-structure which resembles our model-predicted transient *occ* state might represent a post-product-dissociation stage; the hydrolysis-slowing mutation might have kinetically stabilized this otherwise transient intermediate. Similarly, we suggest the 120°-structure might have been captured shortly after ATP binding, as the $\beta_1$-subunit begins to close.

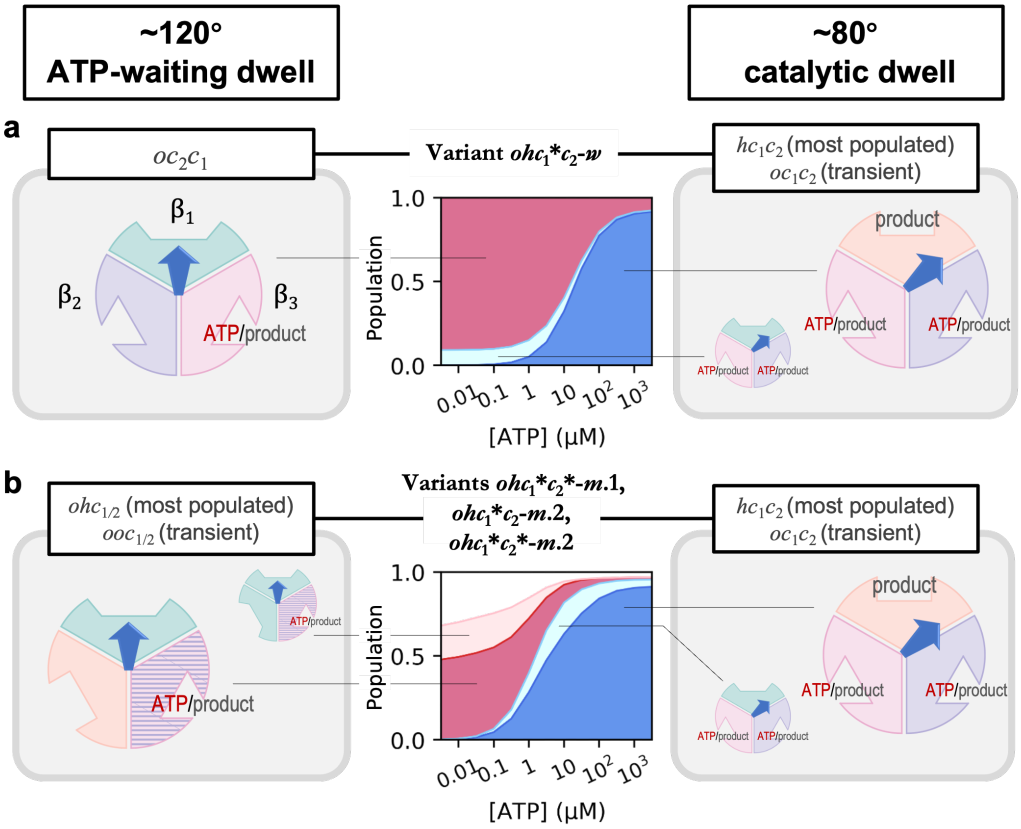


**Supplementary Fig. 10**: **[ATP]-dependent steady-state populations of the major functional stages**

Stacked area plots (middle panels) show the model-predicted probability distributions of the major functional stages within the 80°- (blue shadows) and 120°-dwells (pink shadows) as a function of [ATP]. These data supplement the model predictions presented in Supplementary Figure 10, as detailed in Supplementary Section Supplementary Note 7.

**a** Predictions by the $ohc_1^*c_2^*$-$w$ variant. The 120°-dwell is dominated by an $oc_2c_1$ state (dark pink). As discussed in Supplementary Section Supplementary Note 7, although this state involves a closed $c_2$ conformation of the $\beta_2$-subunit, it functions as an intermediate connecting the active $c_1$ and open $h$ conformations, effectively behaving as an *ohc*-like functional state.

**b** Predictions by the $ohc_1^*c_2^*$-$m.1$, $ohc_1^*c_2$-$m.2$, and $ohc_1^*c_2^*$-$m.2$ variants. The 120°-dwell is dominated by the *ohc* state (one closed, one half-closed, one open; dark pink), coexisting with a lower-populated *ooc* state (light pink). In both **a** and **b**, the 80°-dwell is consistently dominated by the *hcc* state (dark blue), representing the stable pre-product-dissociation stage, while the *occ* state (light blue) remains transient across all ATP concentrations.

Source data are provided as a Source Data file.

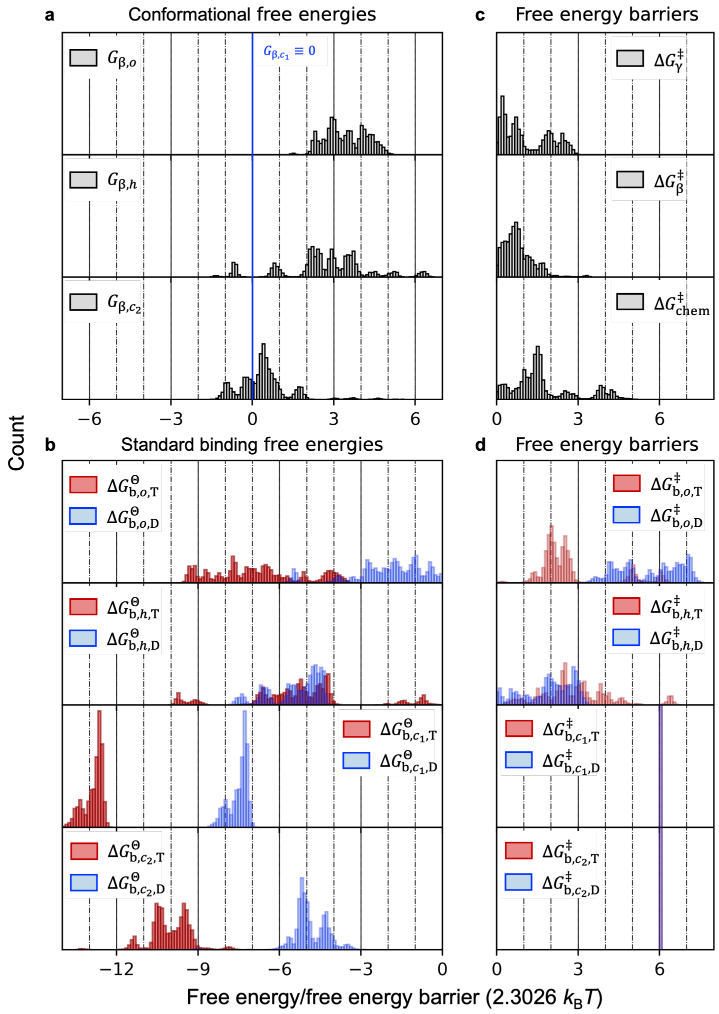


**Supplementary Fig. 11**: **Distributions of the trained parameters for the** $ohc_1^* c_2$-$w$ **variant** Histograms showing the frequencies of the trained parameters (free energies and free energy barriers) for the $ohc_1^* c_2^*$-$w$ variant. Note that these distributions do not strictly reflect the true marginalized posterior distributions. The x-axis is in units of $2.3026\, k_B T$ (equivalent to $\ln(10)\, k_B T$); thus, a difference of 1 unit corresponds to a tenfold change in the corresponding equilibrium constant or rate coefficient.

**a** Conformational free energies ($G_{\beta,\mathcal{C}}$) for the open ($o$), half-closed ($h$), and closed ($c_2$) conformations, relative to the reference active closed conformation ($G_{\beta,c_1} \equiv 0$, indicated by the vertical blue line).

**b** Standard binding free energies ($\Delta G^{\ominus}_{\mathrm{b},\mathcal{C},\mathcal{B}}$) for ATP (red) and ADP (blue) binding to each conformation. Note that $\Delta G^{\ominus}_{\mathrm{b},c_1,\mathrm{T}}$ and $\Delta G^{\ominus}_{\mathrm{b},c_1,\mathrm{D}}$ are bound by Equation 7.

**c** Global free energy barriers for γ-rotation ($\Delta G^{\ddagger}_{\gamma}$), β-conformational transition ($\Delta G^{\ddagger}_{\beta}$), and reversible ATP hydrolysis/synthesis ($\Delta G^{\ddagger}_{\mathrm{chem}}$).

**d** Free energy barriers for ATP (red) and product (blue) binding ($\Delta G^{\ddagger}_{b,\mathcal{C},\mathcal{B}}$). Note that the binding barriers for the $c_1$ and $c_2$ conformations are fixed in the prior (indicated by the purple line), assuming nucleotide exchange in these closed conformations is slow.

Source data are provided as a Source Data file.

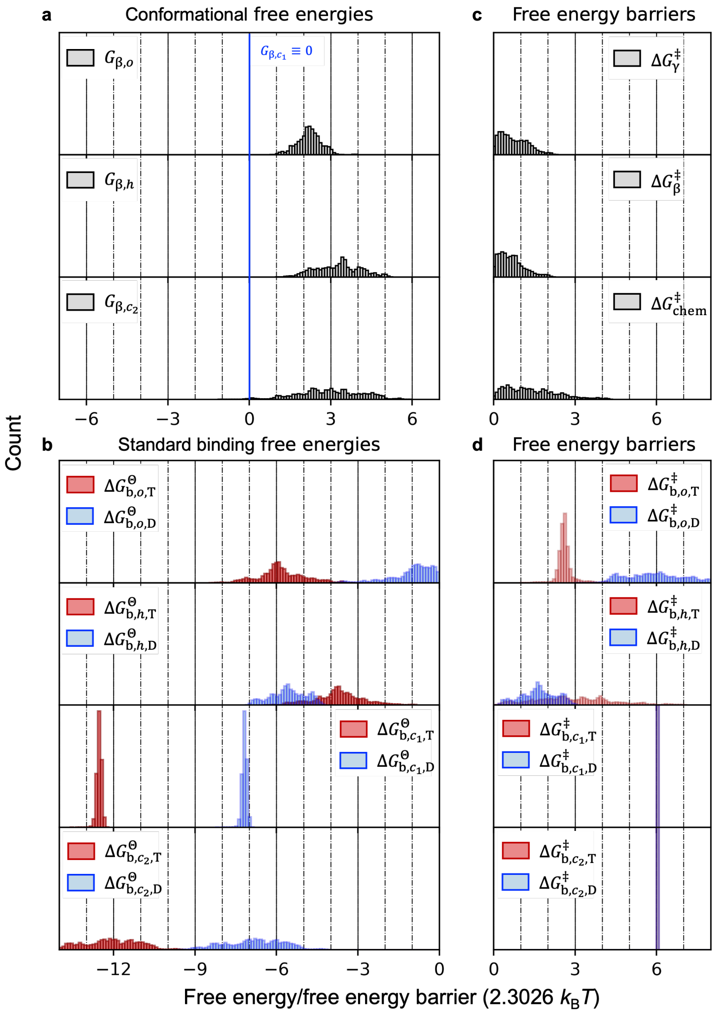


**Supplementary Fig. 12: Distributions of the trained parameters for the $ohc_1^*c_2^*$-$m$.1 variant**

Histograms showing the frequencies of the trained parameters (free energies and free energy barriers) for the $ohc_1^*c_2^*$-$w$ variant. Note that these distributions do not strictly reflect the true marginalized posterior distributions. The x-axis is in units of $2.3026\,k_\mathrm{B}T$ (equivalent to $\ln(10)\,k_\mathrm{B}T$); thus, a difference of 1 unit corresponds to a tenfold change in the corresponding equilibrium constant or rate coefficient.

**a** Conformational free energies ($G_{\beta,\mathcal{C}}$) for the open ($o$), half-closed ($h$), and closed ($c_2$) conformations, relative to the reference active closed conformation ($G_{\beta,c_1} \equiv 0$, indicated by the vertical blue line).

**b** Standard binding free energies ($\Delta G^{\ominus}_{\mathrm{b},\mathcal{C},\mathcal{B}}$) for ATP (red) and ADP (blue) binding to each conformation. Note that $\Delta G^{\ominus}_{\mathrm{b},\mathcal{C},\mathrm{T}}$ and $\Delta G^{\ominus}_{\mathrm{b},\mathcal{C},\mathrm{D}}$ are bound by Equation 7 for $\mathcal{C} = c_1$, $c_2$.

**c** Global free energy barriers for γ-rotation ($\Delta G^{\ddagger}_{\gamma}$), β-conformational transition ($\Delta G^{\ddagger}_{\beta}$), and reversible ATP hydrolysis/synthesis ($\Delta G^{\ddagger}_{\mathrm{chem}}$).

**d** Free energy barriers for ATP (red) and product (blue) binding ($\Delta G^{\ddagger}_{b,\mathcal{C},\mathcal{B}}$). Note that the binding barriers for the $c_1$ and $c_2$ conformations are fixed in the prior (indicated by the purple line), assuming nucleotide exchange in these closed conformations is slow.

Source data are provided as a Source Data file.

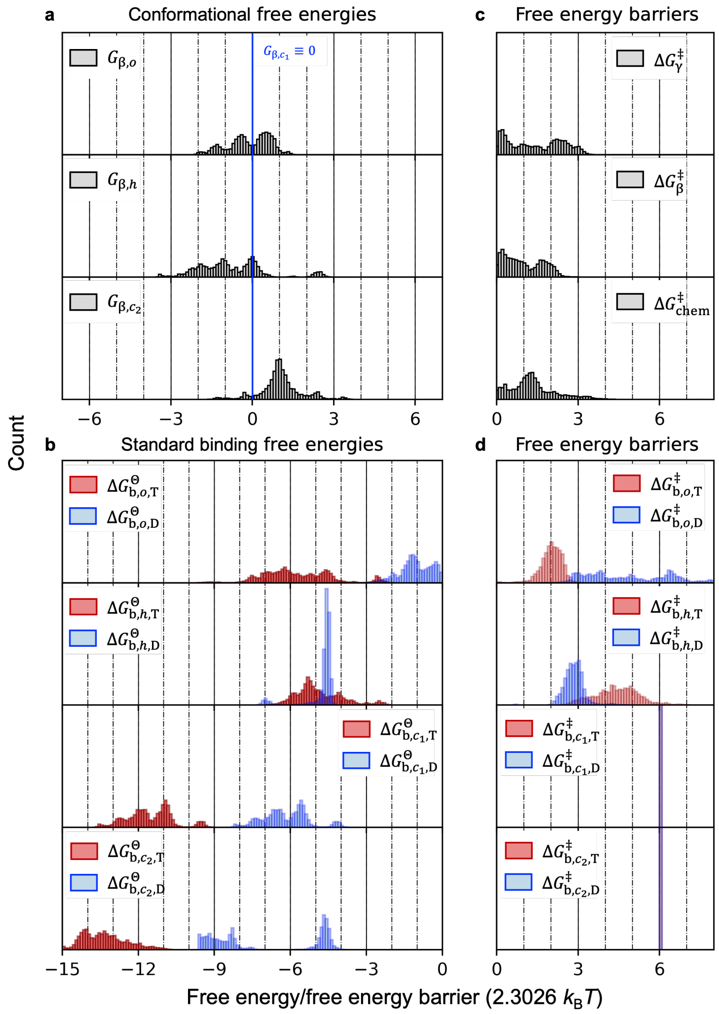


**Supplementary Fig. 13**: **Distributions of the trained parameters for the $ohc_1^*c_2$-$m$.2 variant**

Histograms showing the frequencies of the trained parameters (free energies and free energy barriers) for the $ohc_1^*c_2^*$-$w$ variant. Note that these distributions do not strictly reflect the true marginalized posterior distributions. The x-axis is in units of $2.3026\,k_\mathrm{B}T$ (equivalent to $\ln(10)\,k_\mathrm{B}T$); thus, a difference of 1 unit corresponds to a tenfold change in the corresponding equilibrium constant or rate coefficient.

**a** Conformational free energies ($G_{\beta,\mathcal{C}}$) for the open ($o$), half-closed ($h$), and closed ($c_2$) conformations, relative to the reference active closed conformation ($G_{\beta,c_1} \equiv 0$, indicated by the vertical blue line).

**b** Standard binding free energies ($\Delta G^\ominus_{\mathrm{b},\mathcal{C},\mathcal{B}}$) for ATP (red) and ADP (blue) binding to each conformation. Note that $\Delta G^\ominus_{\mathrm{b},c_1,\mathrm{T}}$ and $\Delta G^\ominus_{\mathrm{b},c_1,\mathrm{D}}$ are bound by Equation 7.

**c** Global free energy barriers for γ-rotation ($\Delta G^\ddagger_\gamma$), β-conformational transition ($\Delta G^\ddagger_\beta$), and reversible ATP hydrolysis/synthesis ($\Delta G^\ddagger_\mathrm{chem}$).

**d** Free energy barriers for ATP (red) and product (blue) binding ($\Delta G^\ddagger_{b,\mathcal{C},\mathcal{B}}$). Note that the binding barriers for the $c_1$ and $c_2$ conformations are fixed in the prior (indicated by the purple line), assuming nucleotide exchange in these closed conformations is slow.

Source data are provided as a Source Data file.

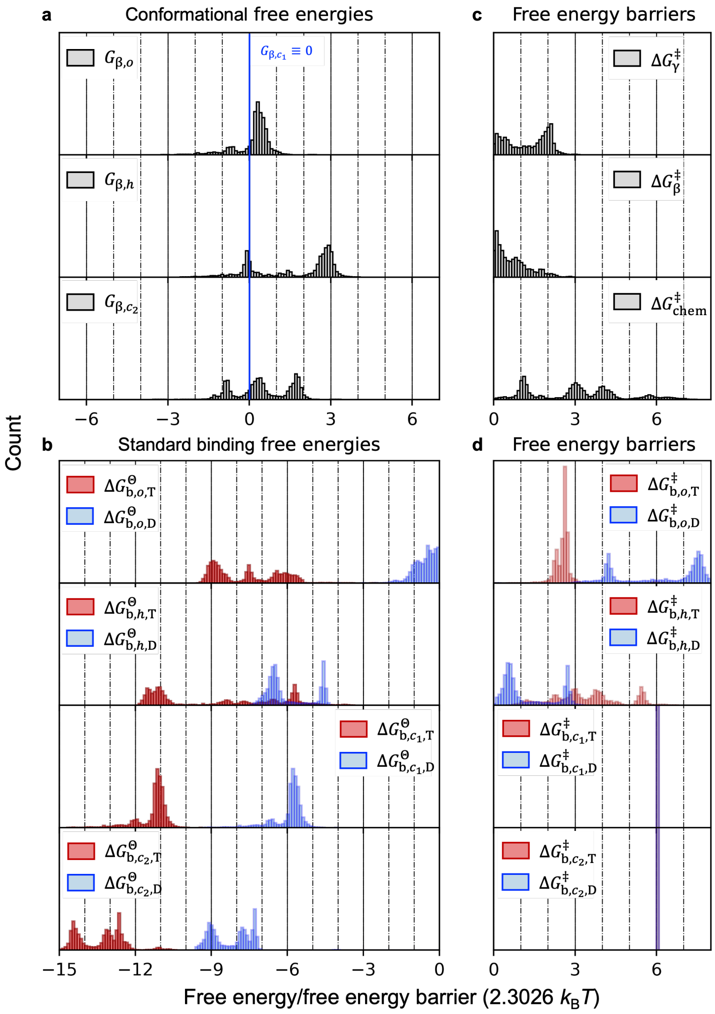


**Supplementary Fig. 14**: **Distributions of the trained parameters for the $ohc_1^*c_2^*$-$m$.2 variant**

Histograms showing the frequencies of the trained parameters (free energies and free energy barriers) for the $ohc_1^*c_2^*$-$w$ variant. Note that these distributions do not strictly reflect the true marginalized posterior distributions. The x-axis is in units of $2.3026\, k_B T$ (equivalent to $\ln(10)\, k_B T$); thus, a difference of 1 unit corresponds to a tenfold change in the corresponding equilibrium constant or rate coefficient.

**a** Conformational free energies ($G_{\beta,\mathcal{C}}$) for the open ($o$), half-closed ($h$), and closed ($c_2$) conformations, relative to the reference active closed conformation ($G_{\beta,c_1} \equiv 0$, indicated by the vertical blue line).

**b** Standard binding free energies ($\Delta G^{\ominus}_{\mathrm{b},\mathcal{C},\mathcal{B}}$) for ATP (red) and ADP (blue) binding to each conformation. Note that $\Delta G^{\ominus}_{\mathrm{b},\mathcal{C},\mathrm{T}}$ and $\Delta G^{\ominus}_{\mathrm{b},\mathcal{C},\mathrm{D}}$ are bound by Equation 7 for $\mathcal{C} = c_1, c_2$.

**c** Global free energy barriers for γ-rotation ($\Delta G^{\ddagger}_{\gamma}$), β-conformational transition ($\Delta G^{\ddagger}_{\beta}$), and reversible ATP hydrolysis/synthesis ($\Delta G^{\ddagger}_{\mathrm{chem}}$).

**d** Free energy barriers for ATP (red) and product (blue) binding ($\Delta G^{\ddagger}_{b,\mathcal{C},\mathcal{B}}$). Note that the binding barriers for the $c_1$ and $c_2$ conformations are fixed in the prior (indicated by the purple line), assuming nucleotide exchange in these closed conformations is slow.

Source data are provided as a Source Data file.

### Supplementary Tables (referenced in the main article)

**Supplementary Table 1: Prior distributions of the model parameters**
The ranges for these free energies and energy barriers are defined in units of 2.3026 $k_B T$, which is equivalent to log(10) $k_B T$. This choice of unit facilitates a direct comparison of the order of magnitude of the corresponding Boltzmann factors, as a difference of one unit in a free energy or free energy barrier corresponds to a tenfold change in the corresponding transition rate.

| **Parameter** | **Prior**<br>(Unit: 2.3026 $k_B T$) |
|---|---|
| $G_{\beta,\mathcal{C}}$ ($\mathcal{C} = o, h$ or $\mathcal{C} = o, h, c_2$) | Uniform(-10, 10) |
| $G_{\beta,c}$ or $G_{\beta,c_1}$ | Fixed at 0 |
| $\Delta G^{\ominus}_{\beta,\mathcal{C},\mathcal{B}}$ ($\mathcal{C} = o, h, c$ or $o, h, c_1, c_2$, $\mathcal{B} =$ T, D) | Uniform(-20, 0) |
| $\Delta G^{\ddagger}_{\beta,\mathcal{C},\mathcal{B}}$ ($\mathcal{C} = o, h$, $\mathcal{B} =$ T, D) | Uniform(0, 9) |
| $\Delta G^{\ddagger}_{\beta,\mathcal{C},\mathcal{B}}$ ($\mathcal{C} = c$ or $c_1, c_2$) | Fixed at 6 |
| $\Delta G^{\ddagger}_{\gamma}$, $\Delta G^{\ddagger}_{\beta}$, $\Delta G^{\ddagger}_{\text{chem}}$ | Uniform(0, 9) |

**Supplementary Table 2: Definition of transition rates for the elementary transitions**

A global pre-exponential factor $f_{\mathrm{att}} = 10^9\,\mathrm{s}^{-1}$ (attempt frequency) appears in all equations. In Equations (1), (2), $G_i \equiv G(\mathbf{s}_i, c_{\mathrm{T}}, c_{\mathrm{D}})$ and $G_j \equiv G(\mathbf{s}_j, c_{\mathrm{T}}, c_{\mathrm{D}})$ are the free energies of states $i$ and $j$, respectively (Equation (2)). These two definition limits the maximum rates of γ-subunit rotations and β-subunit conformational transitions to $r_{\gamma}^{\max}$ and $r_{\beta}^{\max}$, respectively.

| **Elementary transition (Molecularity)** | **Transition rate expression** ($r_{i\to j}$) | **Free energy barrier(s) introduced as model parameter(s)** |
|---|---|---|
| Conformational transitions of a β-subunit (Uni-molecular) | $$r_{i\to j} = \begin{cases} r_{\beta}^{\max}, & G_j - G_i \le 0; \\ r_{\beta}^{\max} \exp\left(-\frac{G_j - G_i}{k_{\mathrm{B}}T}\right), & G_j - G_i > 0; \end{cases} \quad (1)$$ with maximum rate $r_{\beta}^{\max} = f_{\mathrm{att}} \exp\left(-\frac{\Delta G_{\beta}^{\ddagger}}{k_{\mathrm{B}}T}\right)$ | A uniform free energy barrier $\Delta G_{\beta}^{\ddagger}$ for all β-subunit conformational transitions regardless of the initial and final conformations |
| Stepwise rotations of the γ-subunit (Uni-molecular) | $$r_{i\to j} = \begin{cases} r_{\gamma}^{\max}, & G_j - G_i \le 0; \\ r_{\gamma}^{\max} \exp\left(-\frac{G_j - G_i}{k_{\mathrm{B}}T}\right), & G_j - G_i > 0; \end{cases} \quad (2)$$ with $r_{\gamma}^{\max} = f_{\mathrm{att}} \exp\left(-\frac{\Delta G_{\gamma}^{\ddagger}}{k_{\mathrm{B}}T}\right)$. | A uniform free energy barrier $\Delta G_{\gamma}^{\ddagger}$ for all γ-subunit rotations, including the ±40° and ±80° substeps |
| Reversible ATP hydrolysis/synthesis (Uni-molecular) | $$r_{i\to j} = r_{j\to i} = f_{\mathrm{att}} \exp\left(-\frac{\Delta G_{\mathrm{chem}}^{\ddagger}}{k_{\mathrm{B}}T}\right) \quad (3)$$ | A uniform free energy barrier $\Delta G_{\mathrm{chem}}^{\ddagger}$ for ATP hydrolysis in any catalytically active β-subunit |
| ATP/product binding (Bi-molecular) | $$r_{i\to j} = f_{\mathrm{att}} \exp\left(-\frac{\Delta G_{b,\mathcal{C},\mathcal{B}}^{\ddagger}}{k_{\mathrm{B}}T}\right) c_{\mathcal{B}}, \quad (4)$$ with $\mathcal{B} = \mathrm{T}$ or D for ATP and ADP binding, respectively. | A distinct free energy barrier $\Delta G_{b,\mathcal{C},\mathcal{B}}^{\ddagger}$ is assigned to each β-subunit conformation $\mathcal{C}$ for each binding event |
| ATP/product dissociation (Uni-molecular) | $$r_{j\to i} = f_{\mathrm{att}} \exp\left(-\frac{\Delta G_{\mathrm{b},\mathcal{C},\mathcal{B}}^{\ddagger} - \Delta G_{\mathrm{b},\mathcal{C},\mathcal{B}}^{\ominus}}{k_{\mathrm{B}}T}\right) \quad (5)$$ | No additional parameters |

**Supplementary Table 3**: **Chemo-mechanical coupling scheme predicted by variant *ohc**-*s***

Summary of the most likely $\gamma$-subunit orientations at which the three major catalytic steps (ATP binding, ATP hydrolysis, and product dissociation) occur in the $\beta_3$-subunit, as predicted by the four distinct clusters of trained parameter sets identified for the *ohc*-s variant (see Figure fig:ohc-se). The angles denote the absolute orientation of the $\gamma$-subunit. Values in parentheses indicate the rotation angle relative to the ATP binding event at low [ATP]. Note that none of these four predicted schemes matches the consensus chemo-mechanical coupling scheme derived from experimental data (Table 6 and Supplementary Fig. 2), particularly regarding the relative timing of ATP hydrolysis (+200°) and product dissociation (+320°). These discrepancies serve as a key criterion for excluding the *ohc*-s variant.

| **Cluster** | **ATP binding** | | **ATP hydrolysis** | **Product dissociation** | |
|---|---|---|---|---|---|
| | Low [ATP] | High [ATP] | Low and high [ATP] | Low [ATP] | High [ATP] |
| 1 | 360° (0°) | 240°, 320° (-120°, -40°) | 120°(+120°) | 120°, 200° (+120°, +200°) | 200°, 320° (+200°, +320°) |
| 2 | 360° (0°) | 320° (-40°) | 80°, 120° (+80°, +120°) | 240°, 360° (+240°, +360°) | 320° (+320°) |
| 3 | 240°, 360° | 320° | 80°, 120° | 240°, 360° | 320° |
| 4 | 200° (0°) | 200° (0°) | 80°, 120° (+240°, +280°) | 240° (+400°) | 240° (+400°) |

**Supplementary Table 4**: **Physical constraints from the consensus chemo-mechanical coupling scheme**

Note that all Family B variants (Equation (7)) are excluded as reasoned in Results.

| **Condition** | **Physical/functional rationale and experimental support** |
|---|---|
| $\Gamma_3^{360^\circ} \neq \Gamma_3^{80^\circ}, \Gamma_3^{200^\circ} \neq \Gamma_3^{240^\circ}, \Gamma_3^{320^\circ} \neq \Gamma_3^{360^\circ}.$ (6) | According to the fundamental binding-change principle [6, 15–17, 35, 68, 73], occurrence of a catalytic step at $\phi$ requires the subsequent rotation to induce change of the accessible conformational ensemble, i.e., the $\gamma$-$\beta$ restrictions. |
| $(\Gamma_3^{320^\circ}, \Gamma_3^{360^\circ}) = \begin{cases} (\{o,h\},\{o\}), & \text{Family A variants;} \\ (\{o\},\{o,h\}), & \text{Family B variants (excluded).} \end{cases}$ (7) | Product dissociation at 320° and ATP binding at 360° require both $\Gamma_3^{320^\circ}$ and $\Gamma_3^{360^\circ}$ to exclude all closed conformation(s) ($c$ or $c_1/c_2$). |
| $\Gamma_3^{200^\circ} = \begin{cases} \{o,h,c\}, & \text{for } ohc\text{-variants;} \\ \{o,h,c_1\} \text{ or } \{o,h,c_1,c_2\}, & \text{for } ohc_1c_2\text{-variants.} \end{cases}$ (8) <br> $\Gamma_3^{240^\circ} = \begin{cases} \{o\} \text{ or } \{o,h\}, & \text{for } ohc\text{- and } ohc_1^*c_2^*\text{-variants;} \\ \{o\}, \{o,h\}, \text{ or } \{o,h,c_2\}, & \text{for } ohc_1^*c_2\text{-variants.} \end{cases}$ (9) | ATP hydrolysis at 200° requires $\Gamma_3^{200^\circ}$ to include at least one catalytically active conformation, while $\Gamma_3^{240^\circ}$ to exclude any catalytically active conformation. |
| For *ohc*-variants, <br> If $c \in \Gamma_3^{80^\circ}, \quad c \in \Gamma_3^{120^\circ}.$ (10) <br> For $ohc_1^*c_2$-variants, <br> If $c_1 \in \Gamma_3^{80^\circ}, \quad c_1 \in \Gamma_3^{120^\circ}.$ (11) <br> For $ohc_1^*c_2^*$-variants, <br> If $\Gamma_3^{80^\circ} = \{o,h,c_1\}, \quad c_1 \in \Gamma_3^{120^\circ};$ (12) <br> if $\Gamma_3^{80^\circ} = \{o,h,c_2\}, \quad c_2 \in \Gamma_3^{120^\circ};$ (13) <br> if $\Gamma_3^{80^\circ} = \{o,h,c_1,c_2\}, \quad c_1 \in \Gamma_3^{120^\circ}$ or $c_2 \in \Gamma_3^{120^\circ}.$ (14) | ATP hydrolysis at 200° requires combinations of $\Gamma_3^{80^\circ}$ and $\Gamma_3^{120^\circ}$ to allow the $\beta_3$-subunit, once converted to a catalytically active conformation at 80° or 120°, to stay in the catalytically active conformation(s) up to 200°. |

**Supplementary Table 5**: **Additional constraints for selecting plausible $ohc_1c_2$-variants**

| Condition | Physical/functional rationale and experimental support |
|---|---|
| Condition *structure*: $\Gamma_2^{80°}, \Gamma_3^{80°} = \{o,h,c_1\}, \{o,h,c_2\}, \text{or } \{o,h,c_1,c_2\}.$ (15) | Available structures of $F_1$-ATPase show two closed β-subunits in the 80°-dwell [32, 65]. Thus, both $\Gamma_2^{80°}$ and $\Gamma_3^{80°}$ should include at least one closed conformation. |
| Condition *monotonic*: $\Gamma_3^{80°} \cap \Gamma_3^{120°}, \Gamma_3^{120°} \cap \Gamma_3^{200°} = \{c_1\}, \{c_2\}, \text{ or } \{c_1, c_2\};$ (16) $\Gamma_3^{240°} \subseteq \Gamma_3^{200°}, \quad \Gamma_3^{320°} \subseteq \Gamma_3^{240°}.$ (17) | For physical parsimony, the closing (360° → 80° → 120° → 200°) and re-opening (200° → 240° → 360°) of the $β_3$-subunit in a catalytic cycle are assumed to progress monotonically without regression (see explanations in Supplementary Note 3 & Supplementary Note 4). |
| Condition *asymmetric*: $\Gamma_1^{\phi} \neq \Gamma_2^{\phi}, \Gamma_1^{\phi} \neq \Gamma_3^{\phi}, \Gamma_2^{\phi} \neq \Gamma_3^{\phi}, \text{ for } \phi = 80°, 120°.$ (18) | The intrinsically asymmetric γ-subunit is likely to impose different restrictions on the three β-subunits in both the 80°- and 120°-dwells [4, 6, 17, 19, 22, 32, 64, 65]. Thus, both $\Gamma_k^{80°}$ and $\Gamma_k^{120°}$ should be unique for $k = 1, 2, 3$. |

**Supplementary Table 6: Calculation of basic observables from the steady-state distribution**
$\rho_i^{\mathrm{st}}$ is the steady-state population of state $i$. $n_i^{\mathcal{B}}$ is the number of $\mathcal{B}$-bound sites in state $i$, $\mathcal{B}$=T, D for ATP and ADP, respectively. $f_{i\to j}$ is the net flux from state $i$ to $j$ (Equation (10)). Summation limits specify relevant states or transitions.

| Observable | Calculation Method | Description |
|---|---|---|
| ATP/ADP occupancy ($\nu_{\mathrm{T}}$, $\nu_{\mathrm{D}}$) | $$\nu_{\mathcal{B}} = \sum_i \rho_i^{\mathrm{st}} n_i^{\mathcal{B}}, \text{ for } \mathcal{B} = \mathrm{T}, \mathrm{D} \quad (19)$$ | Ensemble average of the number of occupied catalytic sites per $F_1$-ATPase molecule.<br>Note: all β-subunits of the binding state D are counted as ADP-bound. Whereas D merges the ADP-, Pi- and ADP+Pi-bound states, we assume that the Pi-bound state is of low population, as supported by Refs. [20, 22, 32]. |
| Populations of the 80°- and 120°-dwells ($\rho_{80^\circ}$, $\rho_{120^\circ}$) | $$\rho_\phi = \sum_{\{i\mid\phi_i=\phi\}} \rho_i^{\mathrm{st}}, \text{ for } \phi = 80^\circ, 120^\circ, \quad (20)$$ summation over all states $i$ whose γ-orientation $\phi_i = 80^\circ$ (80°-states) or 120° (120°-states). | Total populations of the 80°-states and 120°-states. |
| Turnover rate ($k_{\mathrm{cat}}$) | $$k_{\mathrm{cat}} = \sum_{\{(i,j)\in\mathrm{hydrolysis}\}} f_{i\to j}, \quad (21)$$ summation over all pairs of states $(i,j)$ for which the transition from $i \to j$ represents ATP hydrolysis. | Total net flux corresponding to ATP hydrolysis across the entire state network (equivalent to the net flux of ATP binding or product dissociation). |
| Rotation rate ($k_{\mathrm{rot}}$) | $$k_{\mathrm{rot}} = \sum_{\{(i,j)\in\mathrm{rotation}\}} f_{i\to j}, \quad (22)$$ summation over all pairs of states $(i,j)$ for which the transition from $i \to j$ represents γ-subunit rotation from 80° to 120°. | Total net flux from all 80°-states to their subsequent 120°-states (representing the +40° substep). |
| Orientation- and site-specific rate of a catalytic step ($R_\phi^{\Pi,k}$) | $$R_\phi^{\Pi,k} = \sum_{\{(i,j)\in(\Pi,\beta_k,\phi)\}} f_{i\to j}, \quad (23)$$ summation over all pairs of states $(i,j)$ that meet two conditions: both states have the γ-orientation $\phi$, and the transition $i \to j$ represents catalytic step $\Pi$ occurring in the $\beta_k$-subunit. | Total net flux corresponding to a catalytic step $\Pi$ occurring in the $\beta_k$-subunit when the γ-orientation is $\phi$. |

**Supplementary Table 7**: **Components of the likelihood function**

| Observable | Likelihood function | Rationale |
|---|---|---|
| Turnover rates | Log-normal distribution: $P(k_{\text{cat},l}^{\text{exp}}\|\mathbf{\Omega}) =$ $$\frac{1}{\sqrt{2\pi}\sigma_\kappa}\exp\left(-\frac{\left(\log_{10} k_{\text{cat},l}^{\text{exp}} - \log_{10} k_{\text{cat},l}^{\text{mod}}(\mathbf{\Omega})\right)^2}{2\sigma^2}\right), \quad (24)$$ with the standard deviation $\sigma = 1$ | Measured turnover rates at [ATP] from micromolar to millimolar span orders of magnitude. This choice of log-normal distribution ensures that our Bayesian approach only aims to reproduce the correct order of magnitude of the measured rates, rather than their exact numerical values. |
| Chemo-mechanical coupling efficiencies | Normal distribution: $P(\eta_l^{\text{exp}}\|\mathbf{\Omega}) =$ $$\frac{1}{\sqrt{2\pi}\sigma_\eta}\exp\left(-\frac{\left(\eta_l^{\text{exp}} - \eta_l^{\text{mod}}(\mathbf{\Omega})\right)^2}{2\sigma_\eta^2}\right), \quad (25)$$ with $\sigma_\eta = 0.2$ | We assumed normally distributed errors around the expected value ($\eta_l^{\text{exp}} = 100\%$) [8]. |
| Nucleotide occupancies | Normal distribution: $P(\nu_l^{\text{exp}}\|\mathbf{\Omega}) =$ $$\frac{1}{\sqrt{2\pi}\sigma_\nu}\exp\left(-\frac{\left(-\nu_l^{\text{exp}} - \nu_l^{\text{mod}}(\mathbf{\Omega})\right)^2}{2\sigma_\nu^2}\right), \quad (26)$$ with $\sigma_\nu = 0.08$ | The residuals of the measured ATP/ADP titration curves [21] compared to the equilibrium binding model fits (Equation (17)) are approximately normally distributed. $\sigma_\nu$ is estimated from these residuals. |

### Supplementary Note 1 100% efficiency requires $c$ to be the most stable conformation for a nucleotide-bound β-subunit

In the main article, we have stated the conclusion that the $c$ (closed and catalytically active) conformation being the most stable conformation for a nucleotide-bound β-subunit is a necessary condition for our model to reproduce the near 100% efficiency. Here, we present the evidence that leads us to this conclusion.

In Results, we have shown for variant *ohc-s* that all the trained parameter sets share the property that nucleotide-bound β-subunit has the lowest free energy when adopting the catalytically active, closed conformation $c$ (Figure 2f). However, because the training data included in addition to the 100% efficiency also the measured ATP/ADP titration curves, it is not conclusive from Figure 2f alone whether this common property of the trained parameter sets is a requirement for our model to reproduce the 100% efficiency or the titration curves.

To address this question, we carried out a another round of training of variant *ohc-s* using our Bayesian approach (Methods), where only the experimental turnover rates and 100% efficiency were included as training data. Notably, the experimental turnover rates were still included in the training here to find meaningful parameter sets that predict effective rotary catalysis; otherwise, the training may be stuck in a region of the parameter space where hardly any catalysis or rotation is predicted, although the efficiency as a ratio of turnover and rotational rates could still be near 100%.

From the parameter sets found in this training we chose the sets that reproduce the experimental turnover rates and 100% efficiency. For these parameters, Supplementary Fig. 8c shows the free energies of an ATP- or ADP-bound β-subunit adopting the $o$ or $h$ conformation ($G_{\beta,\mathcal{C}} + \Delta G^{\ominus}_{\mathrm{b},\mathcal{C},\mathcal{B}}$ for $\mathcal{C} = o, h$ and $\mathcal{B} = \mathrm{T}, \mathrm{D}$) against the free energy of an ATP-bound β-subunit adopting the $c$ conformation ($\Delta G^{\ominus}_{\mathrm{b},c,\mathrm{T}}$). Similar to the previous observation (Figure 2f), here, all the parameter sets share the property that a nucleotide-bound β-subunit has the lowest free energy when adopting the catalytically active, closed conformation $c$.

Additionally, we also trained the other two investigated *ohc*-variants (variants *ohc-w* and *ohc-m*) against only the experimental turnover rates and 100% efficiency. Similar with variant *ohc-s*, parameter sets were found for both variants to reproduce these training data; all these parameter sets share the property that a nucleotide-bound β-subunit has the lowest free energy when adopting the catalytically active, closed conformation $c$ (Supplementary Fig. 8a, b).

These results provide strong evidence for our conclusion that the $c$ conformation being the most stable conformation for a nucleotide-bound β-subunit is a necessary condition for our model to reproduce the near 100% efficiency. This condition is mathematically expressed by

$$G_{\beta,\mathcal{C}} + \Delta G^{\Theta}_{\mathrm{b},\mathcal{C},\mathcal{B}} > \Delta G^{\Theta}_{\mathrm{b},c,\mathrm{T}} \tag{27}$$

for $\mathcal{C} = o, h$ and $\mathcal{B} = \mathrm{T}, \mathrm{D}$. In the main article (Results), we have also provided a plausible explanation for this condition.

Most importantly, this condition is not restricted to a specific variant of our model; given the plausible explanation, we suggest that it is highly likely a general conclusion depending only on the very basic assumptions of our model (Results) which are actually common notions about $F_1$-ATPase. Therefore, we suggest that $c$ being the most stable conformation is not only the mechanism by which our model reproduces the near 100 %efficiency, but also the way in which $F_1$-ATPases have evolved to maintain the tight chemo-mechanical coupling between ATP hydrolysis within the β-subunits and the rotation of the γ-subunit under a wide range of ATP concentrations.

### Supplementary Note 2 Consensus chemo-mechanical coupling scheme

Here, we summarize a series of single-molecule experiments that are intended to determine the $\gamma$-subunit orientations at which the three major catalytic steps in a $\beta$-subunit occur, namely, ATP binding, ATP hydrolysis and the rate-limiting product dissociation. As explained at the end of this section, these summarized observations, in retrospect, lead to a consensus chemo-mechanical coupling scheme (Table 6 and Supplementary Fig. 2) that assigns each catalytic step to a particular $\gamma$-orientation; we used this consensus chemo-mechanical coupling scheme as validation data to evaluate our Markov model.

#### A brief review of the single-molecule observations

The major technique employed in the single-molecule experiments on $F_1$-ATPase is fluorescence microscopy imaging, which allows direct visualization of the rotation of the $\gamma$-subunit. An $\alpha_3\beta_3\gamma$ subcomplex of $F_1$-ATPase derived from the thermophilic *Bacillus PS3* with several mutations was used. These mutations include $\alpha$-C193S and an extra his-tag (consisted of ten histidines) at the amino terminus of each $\beta$-subunit to immobilize the stator $\alpha_3\beta_3$ onto a surface, as well as $\gamma$-S107C and $\gamma$-I210C to attach a colloidal gold or magnetic bead to the $\gamma$-subunit via a biotin-streptavidin-BSA linker [8, 13, 14]. The bead was imaged under fluorescence microscopy to record a time trace of its orientation; the orientation of the bead is supposed to closely follow the orientation of the $\gamma$-subunit [8, 15, 33, 54, 57].

R. Yasuda *et al.* first resolved the 90°- and 30°-substeps of $\gamma$-subunit rotation from such time traces of bead orientations [8]. These traces show two dwells before and after the 30°-substeps, which are termed the 90°- and 120°-dwell respectively. At low ATP concentrations ([ATP] $\sim$ 0.2 μM and below), the 120°-dwell is dominant while the 90°-dwell is hardly seen. The 120°-dwell becomes longer with decreasing [ATP]; the distribution of the duration is fitted by a single exponential function $f(t) \propto \exp(-k_1 t)$, where the rate $k_1$ is dependent on [ATP], $k_1 \approx (3 \times 10^7\,\mathrm{s}^{-1} \cdot \mathrm{M}^{-1}) \times [\mathrm{ATP}]$. It was therefore hypothesized that the 120°-dwell is limited by ATP binding to an *apo* $\beta$-subunit (ATP-waiting dwell), and the ATP binding rate coefficient is estimated to be $3 \times 10^7\,\mathrm{s}^{-1} \cdot \mathrm{M}^{-1}$. At high [ATP] (60 μM and above), the 90°-dwell becomes dominant while the 120°-dwell is hardly seen. The distribution of the duration of the 90°-dwell is fitted by a double exponential function $f(t) \propto \exp(-k_2 t) - \exp(-k_3 t)$, where the two rates $k_2 = (0.71 \pm 0.02)\,\mathrm{ms}^{-1}$ and $k_3 = (1.64 \pm 0.06)\,\mathrm{ms}^{-1}$ do not depend on [ATP]. It was therefore hypothesized that the 90°-dwell is limited by two $\sim$1 ms reactions, which may be, *e.g.*, ATP hydrolysis and product dissociation. The 90°-dwell is thus termed the catalytic dwell. At intermediate [ATP] (2 μM – 20 μM), both dwells are seen, and the distributions of the dwell times are compatible with the three rate limiting steps of rates $k_1$ ([ATP]-dependent), $k_2$ and $k_3$ ([ATP]-independent).

The two ∼1 ms reactions limiting the catalytic dwell are suggested to be ATP hydrolysis and phosphate dissociation, respectively. First, K. Shimabukuro *et al.* [54] introduced an extra mutation β-E190D into each of the three β-subunits; this mutation had been shown to remarkably slow down ATP hydrolysis. Additionally, they used a slowly hydrolysable ATP analog ATPγS instead of ATP as substrate. As a result, they observed significant elongation of the catalytic dwell, which suggests that ATP hydrolysis is a rate limiting step of the catalytic dwell, and may correspond to one of the two ∼1 ms reactions observed in [8]. Additionally, this study refined the previously reported 90°- and 30°-substeps [8] to be 80° and 40° instead, still within the previously estimated errorbar [8], giving rise to the notions of catalytic dwell around 80° and ATP-waiting dwell around 120° [15, 34, 57].

Further, to determine the exact γ-orientations of ATP hydrolysis in a β-subunit, T. Ariga *et al.* [34] visualized the rotation of hybrid $\alpha_3\beta_3\gamma$ constructs containing one or two β-subunits with mutation β-E190D. The observed time traces of the bead orientations show that in a mutant β-subunit, the catalytic dwell at +200° relative to ATP binding is remarkably elongated, suggesting that ATP hydrolysis in this β-subunit limits the rotation from +200° to +240°. Additionally, the catalytic dwell at +320° is also elongated, suggesting a third catalytic event (∼20 ms) in this β-subunit limiting the rotation from +320° to +360°. However, this study did not determine what this third catalytic event represents.

Second, K. Adachi *et al.* [15] observed that the catalytic dwell is also remarkably elongated when an excessive amount of phosphate is added into the buffer. This observation suggests that the other ∼1 ms reaction within the catalytic dwell is likely phosphate dissociation. Naïvely, phosphate concentration in the buffer does not directly affect the rate of phosphate dissociation; however, as suggested by the authors, high phosphate concentration results in frequent phosphate rebinding, which in turn results in reversal of the +40°-substep and thus elongation of the catalytic dwell.

Combining the conclusions of Refs. [34] and [15], R. Watanabe *et al.* [57] suggested that the third catalytic event observed for the hybrid $F_1$-ATPase constructs at +320° [34] actually represents phosphate dissociation. Using high concentration (1 mM) of ATPγS as substrate, they arrested the γ-subunit within a catalytic dwell for a while using magnetic tweezers; then, they released the γ-subunit and observed whether it proceeded forward to the next catalytic angle or stayed at the stalled angle. They observed these two behaviors of similar frequencies, which suggested that ATP hydrolysis remains reversible at +200°. Because the reversibility of ATP hydrolysis requires the presence of phosphate still in the catalytic site, this study provides evidence for phosphate dissociation later than +200° at high [ATP]. Considering the conclusion from Ref. [15] that phosphate dissociation occurs within a catalytic dwell, the authors concluded that phosphate dissociation occurs at +320°.

Besides ATP binding, ATP hydrolysis, and phosphate dissociation, one would expect the dissociation of the other product, *i.e.*, ADP, to also be a major catalytic step. Unexpectedly, evidences have been reported showing that ADP dissociation is a very fast reaction within the ATP-waiting dwell under room temperature, and occurs between +240° and +320°, earlier than phosphate dissociation [15, 33, 55, 56].

**Discussion about the single-molecule data and synthesis of the consensus chemo-mechanical coupling scheme**

In summary of the above observations, Watanabe *et al.* proposed a deterministic model for the $\gamma$-orientations of the major catalytic steps, to which we refer as the Watanabe model (see particularly Figure 6 in Ref. [57]). At low [ATP], ATP binding in a $\beta$-subunit occurs in a 120°-dwell, and this $\gamma$-orientation is taken as 0°. In this very $\beta$-subunit, the bound ATP is hydrolyzed at +200°; the product ADP is dissociated at +240°, but this reaction is too fast to be observed under room temperature; finally, the other product, phosphate, is dissociated at +320°. At high [ATP], ATP binding may occur immediately after phosphate dissociation at +320°.

We note that several statements in the Watanabe model remain controversial. Below, we attempt to propose a more general and inclusive model that reconciles these controversies and aligns with other experimental data as well as theoretical insights.

First, the $\gamma$-orientations of ADP and phosphate dissociation have been debated in the publications. The opposite opinion argues that phosphate dissociation is fast and precedes ADP dissociation. The crystal and cryoEM structures of $F_1$-ATPase often capture states containing ADP- or ADP+Pi-bound $\beta$-subunits [4, 5, 32], and only in rare cases are $\beta$-subunits bound with only phosphate or phosphate analogs observed [32, 91]. The higher chance to observe ADP- and ADP+Pi-bound states than to observe Pi-bound states indicates that ADP is likely to linger in the catalytic site longer than phosphate. Additionally, it has been reported that ADP has a much higher binding affinity with the $\beta$-subunits than phosphate [92], which renders it physically implausible that phosphate dissociation is a rate-limiting step whereas ADP dissociation is not.

Another evidence against late and slow phosphate dissociation is that under saturating [ATP], the total nucleotide occupancy approaches three [20–22]. If ADP were dissociated as early as +240° whereas phosphate remained in the catalytic site until +320°, and if phosphate dissociation were also a rate-limiting step of the catalytic dwell taking about 1 ms [8], the total nucleotide occupancy would be substantially below three, contradicting the measured occupancy. A possible fix is that phosphate dissociation is actually a fast reaction and occurs at the beginning of the +320° [57].

In all, it is unlikely that phosphate dissociation is both slower and later than ADP dissociation. The remaining possibilities are: (1) ADP dissociation is a rate-limiting step of the +320°-dwell, while phosphate dissociation is a fast reaction; (2) phosphate dissociation is a rate-limiting step of the +320°-dwell, while ADP dissociation is a fast reaction, but ADP lingers in the catalytic site longer. Considering both possibilities, we adopt the view in our consensus chemo-mechanical coupling scheme (Table 6 and Supplementary Fig. 2) that there is only one product dissociation step that is rate-limiting, either ADP dissociation or phosphate dissociation, and this rate-limiting product dissociation step occurs at at +320°.

Second, we note that, according to the above experimental data alone, the possibility cannot be ruled out that under low [ATP], the rate-limiting product dissociation step occurs at $\gamma$-orientations other than +320°. In fact, under low [ATP], the 80°-dwells are much shorter than the 120°-dwells and therefore hardly recognized in the recorded trajectories. Therefore, this consensus chemo-mechanical coupling scheme (Table 6 and Supplementary Fig. 2) only assumes that the rate-limiting product dissociation step occurs at +320° under high [ATP], but allows for variations under low [ATP].

Third, the hypothesis in Watanabe model that ATP hydrolysis occurs at +200° also seems inconclusive. As already explained, this hypothesis is based on the observation of a mutant $F_1$-ATPase hydrolyzing an ATP analog where hydrolysis is drastically slowed down [34]. For native $F_1$-ATPase and substrate, ATP hydrolysis might occur at $\gamma$-orientations earlier than +200°, but not later. Accordingly, we assume in the consensus chemo-mechanical coupling scheme (Table 6 and Supplementary Fig. 2) that ATP hydrolysis occurs no later than +200°.

In summary, the consensus chemo-mechanical coupling scheme (Supplementary Fig. 2) is formulated as following: ATP binding in a $\beta$-subunit under low [ATP] occurs within a 120°-dwell; denoting the $\gamma$-orientation at which ATP binding in $\beta_3$ occurs as 360° (0°), the bound ATP is hydrolyzed no later than +200°, and the rate-limiting product dissociation occurs under high [ATP] at +320°.

### Supplementary Note 3 Reducing the choices of $\gamma$-$\beta$ restrictions via physical constraints

Here, we detail our derivation of physical constraints on the $\gamma$-$\beta$ restrictions from available experimental data of *E. coli* and *Bacillus PS3* $F_1$-ATPase, including the consensus chemo-mechanical coupling scheme (see Table 6 and Supplementary Note 2). This derivation allows us to systematically reduce the huge hyperparameter space of $\gamma$-$\beta$ restrictions for both the *ohc*-variants and $ohc_1c_2$-variants.

For clarity, we will use formal notations for the $\gamma$-$\beta$ restrictions below. Due to the three-fold rotational pseudo-symmetry of the $F_1$-ATPase structure, specifying six $\Gamma_k^\phi$ sets, *e.g.*, $\{\Gamma_k^\phi | k = 1, 2, 3; \phi = 80°, 120°\}$ (denoted as $\{\Gamma_{1,2,3}^{80°,120°}\}$) or $\{\Gamma_3^\phi | \phi = 360°, 80°, 120°, 200°, 240°, 320°\}$ (denoted as $\{\Gamma_3^\phi\}$), is sufficient to uniquely determine a specific variant. These two sets of six $\Gamma_k^\phi$ are equivalent via

$$\Gamma_1^{80°} = \Gamma_3^{320°}, \Gamma_1^{120°} = \Gamma_3^{360°} (\equiv \Gamma_3^{0°}), \Gamma_2^{80°} = \Gamma_3^{200°}, \Gamma_2^{120°} = \Gamma_3^{240°}. \tag{28}$$

The other $\Gamma_k^\phi$, knowing $\{\Gamma_3^\phi\}$, are determined by

$$\Gamma_1^\phi = \Gamma_3^{\mathrm{mod}(\phi+240°,360°)}, \Gamma_2^\phi = \Gamma_3^{\mathrm{mod}(\phi+120°,360°)}, \tag{29}$$

for $\phi = 360°, 80°, 120°, 200°, 240°, 320°$, where $\mathrm{mod}(x, y)$ represents the modulus of $x$ divided by $y$.

The consensus model applies to any of the three $\beta$-subunits. For clarity, we will focus on the $\beta_3$-subunit below and, without loss of generality, define that ATP binding occurs in the $\beta_3$-subunit at the $\gamma$-orientation of 360° (Fig. 1c), so that ATP hydrolysis and product dissociation occur at the $\gamma$-orientations of 200° and 320°, respectively. Accordingly, we will first focus on deriving the conditions of $\{\Gamma_3^\phi\}$, and then convert them to $\{\Gamma_{1,2,3}^{80°,120°}\}$ to provide a compact denotation in tabular form (see Supplementary Table 11 for example).

As stated in Methods, for *ohc*-variants, each $\Gamma_3^\phi$ may be chosen from $\{o\}$, $\{o, h\}$, and $\{o, h, c\}$, with $c$ being the only catalytically active conformation. For $ohc_1c_2$-variants, each $\Gamma_3^\phi$ may be chosen from $\{o\}$, $\{o, h\}$, $\{o, h, c_1\}$, $\{o, h, c_2\}$, and $\{o, h, c_1, c_2\}$, with $c_1$ being catalytically active ($ohc_1^*c_2$-variants), or both $c_1$ and $c_2$ being catalytically active ($ohc_1^*c_2^*$-variants).

#### Conditions from the consensus chemo-mechanical coupling scheme

The most obvious condition is that for ATP hydrolysis to occur at 200°, $\Gamma_3^{200°}$ must include at least one catalytically active conformation (Condition *timing*-I):

$$\Gamma_3^{200°} = \begin{cases} \{o, h, c\}, & \text{for } ohc\text{-variants;} \\ \{o, h, c_1\} \text{ or } \{o, h, c_1, c_2\}, & \text{for } ohc_1c_2\text{-variants.} \end{cases} \quad (30)$$

Here, for $ohc_1^*c_2^*$-variants, because $c_1$ and $c_2$ are functionally equivalent, we require $\Gamma_3^{200°}$ to include $c_1$ (without losing generality), thus excluding the option $\Gamma_3^{200°} = \{o, h, c_2\}$.

ßAdditionally, we note the key finding that for our model to achieve near 100% chemo-mechanical coupling efficiency, the closed, catalytically active conformation must be the energetically most favorable conformation for a nucleotide-bound β-subunit (see Results, Discussion, and Supplementary Note 1). In other words, in our model, a nucleotide-bound β-subunit remains closed unless restricted to the $o$ and $h$ conformations by the γ-subunit. Therefore, for ATP binding and product dissociation to occur at 360° and 320°, respectively, $\Gamma_3^{360°}$ and $\Gamma_3^{320°}$ must exclude all closed conformation(s), thereby forcing the $\beta_3$-subunit to adopt the $o$ or $h$ conformation, which facilitates nucleotide exchange:

$$\Gamma_3^{360°},\ \Gamma_3^{320°} = \{o\} \text{ or } \{o, h\}. \quad (31)$$

Our analysis is extended by considering the fundamental "binding change" idea [6, 16, 67]: rotation of the γ-subunit drives each β-subunit through an ordered, cyclic progression of conformational changes ("open → closed → open" within a full revolution); as the conformation of a β-subunit changes, its nucleotide binding affinity changes too, inducing the major catalytic steps. Combined with our key finding and Supplementary Equations 30, 31, this cyclic progression of the $\beta_3$-subunit must proceed in two phases:

(1) Spontaneous closing from 360° (ATP binding) to 200°(ATP hydrolysis). After ATP binding at 360°, as the γ-subunit rotates further, the steric push must be released, so that the $\beta_3$-subunit (now bound to ATP) can spontaneously relax into the closed, catalytically active conformation by 200°.

(2) Forced re-opening from 200° (ATP hydrolysis) to 320° (product dissociation). After ATP hydrolysis at 200°, the γ-subunit rotation must force the $\beta_3$-subunit out of the catalytically active closed conformation and into an $o$ or $h$ conformation to facilitate product dissociation at 320°.

A general condition of γ-β restrictions implied from the "binding-change idea" is that $\Gamma_3^{\phi}$ must change between the known γ-orientation of each catalytic step and the subsequent orientation (Condition *timing*-II):

$$\Gamma_3^{360^\circ} \neq \Gamma_3^{80^\circ}, \Gamma_3^{200^\circ} \neq \Gamma_3^{240^\circ}, \Gamma_3^{320^\circ} \neq \Gamma_3^{360^\circ}. \tag{32}$$

Moreover, the change of $\Gamma_3^{\phi}$ must both "allow" and "enforce" proper conformational transition of the $\beta_3$-subunit at these orientations that lead to the corresponding catalytic steps.

Particularly, for ATP binding to occur at 360°, $\Gamma_3^{80^\circ}$ must be different from $\Gamma_3^{360^\circ}$ (Supplementary Equation (32)). Further, the $\beta_3$-subunit should be allowed to convert to a catalytically active conformation no later than 200°. Notably, once this conversion occurs, the $\beta_3$-subunit must be allowed to stay in the catalytically active conformation(s) up to 200° (Condition *timing*-III). Otherwise, ATP hydrolysis would occur earlier than 240°. Formally, this condition requires:

For *ohc*-variants,

$$\text{If } c \in \Gamma_3^{80^\circ}, \quad c \in \Gamma_3^{120^\circ}. \tag{33}$$

For $ohc_1^*c_2$-variants,

$$\text{If } c_1 \in \Gamma_3^{80^\circ}, \quad c_1 \in \Gamma_3^{120^\circ}. \tag{34}$$

For $ohc_1^*c_2^*$-variants,

$$\text{If } \Gamma_3^{80^\circ} = \{o, h, c_1\}, \quad c_1 \in \Gamma_3^{120^\circ}; \tag{35}$$

$$\text{if } \Gamma_3^{80^\circ} = \{o, h, c_2\}, \quad c_2 \in \Gamma_3^{120^\circ}; \tag{36}$$

$$\text{if } \Gamma_3^{80^\circ} = \{o, h, c_1, c_2\}, \quad c_1 \in \Gamma_3^{120^\circ} \text{ or } c_2 \in \Gamma_3^{120^\circ}. \tag{37}$$

Next, to enforce the irreversible conversion of the bound ATP to product upon rotation from 200° to 240°, the $\beta_3$-subunit must be forced to convert to an inactive conformation. Thus, $\Gamma_3^{240^\circ}$ must exclude any catalytically active conformation (Condition *timing*-IV):

$$\Gamma_3^{240^\circ} = \begin{cases} \{o\} \text{ or } \{o, h\}, & \text{for } ohc\text{- and } ohc_1^*c_2^*\text{-variants;} \\ \{o\}, \{o, h\}, \text{ or } \{o, h, c_2\}, & \text{for } ohc_1^*c_2\text{-variants.} \end{cases} \tag{38}$$

Finally, for product dissociation to occur at 320°, $\Gamma_3^{320^\circ}$ must be different from $\Gamma_3^{360^\circ}$. Combined with Supplementary Equation (32), there are two possible combinations of $\Gamma_3^{320^\circ}$ and $\Gamma_3^{360^\circ}$ (Condition *timing*-V), defining "Family A" and "Family B" variants, respectively:

$$(\Gamma_3^{320^\circ}, \Gamma_3^{360^\circ}) = \begin{cases} (\{o, h\}, \{o\}), & \text{"Family A" variants;} \\ (\{o\}, \{o, h\}), & \text{"Family B" variants.} \end{cases} \tag{39}$$

#### Condition *monotonic*

Further, we restrict the choices of $\Gamma_3^{80°}$, $\Gamma_3^{120°}$, and $\Gamma_3^{240°}$ with an additional requirement, which we term Condition *monotonic*. This condition posits that the $\beta_3$-subunit must be allowed to close "monotonically" during the closing phase; combinations of $\Gamma_3^{360°}$, $\Gamma_3^{80°}$, and $\Gamma_3^{120°}$ that force the $\beta_3$-subunit to "regress" into a less-closed conformation once it achieves a more-closed conformation are excluded, *e.g.*, the combination $\Gamma_3^{360°} = \{o\}$, $\Gamma_3^{80°} = \{o, h\}$, and $\Gamma_3^{120°} = \{o\}$ that would force the $\beta_3$-subunit to go through conformational changes $o \to h \to o \to c$ from 360° to 200°. While direct experimental evidence for the exact paths is limited, we employ this condition as a minimal model assumption based on physical parsimony and efficiency. A path like $o \to h \to o \to c$ from 360° to 200° is both unnecessarily complex and physically inefficient.

Similarly, this condition also posits that the $\beta_3$-subunit must be enforced to open "monotonically" during the re-opening phase combinations of $\Gamma_3^{240°}$, $\Gamma_3^{320°}$, and $\Gamma_3^{360°}$ that allow the $\beta_3$-subunit to "regress" into a more-closed conformation once it achieves a less-closed conformation are excluded, *e.g.*, the combination $\Gamma_3^{240°} = \{o\}$, $\Gamma_3^{320°} = \{o, h\}$, and $\Gamma_3^{360°} = \{o\}$ that would allow the $\beta_3$-subunit to go through conformational changes $c \to o \to h \to o$ from 200° to 360°.

For *ohc*-variants, Condition *monotonic* is formalized by the expression:

$$\Gamma_3^{360°} \subseteq \Gamma_3^{80°}, \quad \Gamma_3^{80°} \subseteq \Gamma_3^{120°}, \quad \Gamma_3^{120°} \subseteq \Gamma_3^{200°}, \quad \Gamma_3^{320°} \subseteq \Gamma_3^{240°}. \tag{40}$$

For $ohc_1c_2$-variants including two closed conformations $c_1$ and $c_2$, Condition *monotonic* becomes more complex (further detailed in Supplementary Note 4).

#### Family B $ohc_1c_2$-variants are excluded based on product binding affinities

Two families of model variants have been defined above by the choices of $\Gamma_3^{320°}$ and $\Gamma_3^{360°}$ (Supplementary Equation (39)), or equivalently, $\Gamma_1^{80°}$ and $\Gamma_1^{120°}$, with Family A assuming $\Gamma_1^{80°} = \{o, h\}$, $\Gamma_1^{120°} = \{o\}$, and Family B assuming $\Gamma_1^{80°} = \{o\}$, $\Gamma_1^{120°} = \{o, h\}$. These $\gamma$-$\beta$ restrictions dictate that the ADP binding affinity of the low-affinity site ($\beta_1$) decreases for Family A variants upon the 80°→120° rotation, but increases for Family B variants. Below, we explain the reason based on the analytical expression of the binding affinity derived in Supplementary Note 5 (Supplementary Equation (68)).

For Family A variants, the ADP binding affinities at 80° and 120° are:

$$(K_1^{80°})^{-1} = w_o(K_{\mathrm{d},o})^{-1} + w_h(K_{\mathrm{d},h})^{-1},$$
$$(K_1^{120°})^{-1} = (K_{\mathrm{d},o})^{-1}, \tag{41}$$

where the weights $w_o + w_h = 1$ ($w_o > 0, w_h > 0$), $K_{\mathrm{d},o}$ and $K_{\mathrm{d},h}$ are microscopic ADP binding affinities of the $o$ and $h$ conformations ($K_{\mathrm{d},o}, K_{\mathrm{d},h} > 0$). Given our model definition that the $o$ conformation is the low-affinity state and the $h$ conformation is an intermediate state, the plausible condition is $K_{\mathrm{d},o} > K_{\mathrm{d},h}$, thus $K_1^{80^\circ} < K_1^{120^\circ}$ (affinity decreases from 80° to 120°). For Family B variants, the ADP binding affinities at 80° and 120° are reversed:

$$\begin{aligned}(K_1^{80^\circ})^{-1} &= (K_{\mathrm{d},o})^{-1},\\ (K_1^{120^\circ})^{-1} &= w_o(K_{\mathrm{d},o})^{-1} + w_h(K_{\mathrm{d},h})^{-1},\end{aligned} \tag{42}$$

thus $K_1^{80^\circ} > K_1^{120^\circ}$ (affinity increases from 80° to 120°). As reasoned in Results, an increased affinity of the $\beta_1$-subunit from 80° to 120° contradicts with both physical intuition and recent nucleotide titration experiments. Therefore, we exclude all Family B variants and proceed with only Family A variants.

#### Candidate *ohc*-variants

Combining the above five conditions, we deduce the possible choices of $\{\Gamma_3^\phi\}$ for Family A *ohc*-variants:

$$\begin{aligned}&\Gamma_3^{320^\circ} = \{o, h\}, \quad \Gamma_3^{360^\circ} = \{o\},\\ &\Gamma_3^{80^\circ}, \Gamma_3^{120^\circ} = \{o, h\} \text{ or } \{o, h, c\},\\ &\Gamma_3^{200^\circ} = \{o, h, c\}, \quad \Gamma_3^{240^\circ} = \{o, h\}.\end{aligned} \tag{43}$$

Notably, some combinations must be excluded according to Condition *monotonic*. Accordingly, three candidate *ohc*-variants that fully meet Condition *timing* I-IV (thus would be potentially compatible with the consensus chemo-mechanical coupling scheme) are derived, as listed in Supplementary Table 8. However, all of them are excluded because of incompatibility with other experimental data (detailed in the fourth column).

Particularly, we exclude two *ohc*-variants (A2 and A3), because they allow at most one β-subunit to adopt the closed conformation at any γ-orientation, contradicting the experimental observations that suggest two β-subunits of *Bacillus* PS3 $F_1$-ATPase are closed in the catalytic dwell ∼80° [32, 65]. This comparison against observed structures leads to an additional condition of γ-β restrictions (Condition *structure*): both $\Gamma_3^{80^\circ}$ and $\Gamma_3^{200^\circ}$ must include at least one closed conformation.

This analysis corroborates our conclusion in Results that no *ohc*-variant can explain all available experimental data.

**Supplementary Table 8: The five candidate *ohc*-variants**

<table>
<tr><th>Variant</th><th>$\gamma$-$\beta$ restrictions $\{\Gamma_k^\phi\}$</th><th>Number of states</th><th>Reason for exclusion</th></tr>
<tr><td>A1</td><td><table><tr><td>$\phi$</td><td>$\beta_1$</td><td>$\beta_2$</td><td>$\beta_3$</td></tr><tr><td>80°</td><td>$o, h$</td><td>$o, h, c$</td><td>$o, h, c$</td></tr><tr><td>120°</td><td>$o$</td><td>$o, h$</td><td>$o, h, c$</td></tr></table></td><td>648</td><td>Tested in Results (variant *ohc-m*); training failed</td></tr>
<tr><td>A2</td><td><table><tr><td>$\phi$</td><td>$\beta_1$</td><td>$\beta_2$</td><td>$\beta_3$</td></tr><tr><td>80°</td><td>$o, h$</td><td>$o, h, c$</td><td>$o, h$</td></tr><tr><td>120°</td><td>$o$</td><td>$o, h$</td><td>$o, h, c$</td></tr></table></td><td>486</td><td>Would predict at most one closed $\beta$-subunit at any $\gamma$-orientation, disagreeing with cryoEM structures of bacterial $F_1$-ATPase where two $\beta$-subunits adopt similar closed conformations at 80° and/or 120° [32].</td></tr>
<tr><td>A3</td><td><table><tr><td>$\phi$</td><td>$\beta_1$</td><td>$\beta_2$</td><td>$\beta_3$</td></tr><tr><td>80°</td><td>$o, h$</td><td>$o, h, c$</td><td>$o, h$</td></tr><tr><td>120°</td><td>$o$</td><td>$o, h$</td><td>$o, h$</td></tr></table></td><td>432</td><td>Same with A2.</td></tr>
</table>

### Supplementary Note 4 Selection of candidate $ohc_1c_2$-variants

Here we explain our selection of the five most plausible candidate $ohc_1c_2$-variants for rigorous tests via our "training + cross-validation" procedures (Table 3). In Supplementary Note 3, we have derived several constraints from the consensus chemo-mechanical coupling scheme (Conditions *timing* I-V), physical parsimony (Condition *monotonic*), and observed $F_1$-ATPase structures (Condition *structure*). For $ohc_1c_2$-variants, Condition *structure* restricts $\Gamma_3^{80°}$ to $\{o,h,c_1\}$, $\{o,h,c_2\}$, $\{o,h,c_1,c_2\}$. Condition *monotonic* should allow each β-subunit to convert directly between the two closed conformations $c_1$ and $c_2$, without opening (partially) first and closing again ($c_1/c_2 \to h \to c_1/c_2$). Particularly, this condition implies: (1) In the closing phase, $\Gamma_3^{\phi}$ at every two adjacent orientations of 80°, 120°, and 200° should include at least one closed conformation in common:

$$\begin{aligned} \Gamma_3^{80°} \cap \Gamma_3^{120°} &= \{c_1\}, \{c_2\}, \text{ or } \{c_1, c_2\}; \\ \Gamma_3^{120°} \cap \Gamma_3^{200°} &= \{c_1\}, \{c_2\}, \text{ or } \{c_1, c_2\}. \end{aligned} \tag{44}$$

(2) In the opening phase,

$$\Gamma_3^{240°} \subseteq \Gamma_3^{200°}, \quad \Gamma_3^{320°} \subseteq \Gamma_3^{240°}. \tag{45}$$

Combining these conditions still leaves 30 $ohc_1^*c_2^*$-variants and 45 $ohc_1^*c_2$-variants (summarized in Supplementary Table 9 and Supplementary Fig. 15).

**Supplementary Table 9: Possible choices of $\Gamma_3^{\phi}$ for $ohc_1c_2$-variants**
For Family A of $ohc_1c_2$-variants, $\Gamma_3^{320°} = \{o,h\}$, $\Gamma_3^{360°} = \{o\}$; for Family B, $\Gamma_3^{320°} = \{o\}$, $\Gamma_3^{360°} = \{o,h\}$. Those combinations of ($\Gamma_3^{80°}$, $\Gamma_3^{120°}$, $\Gamma_3^{200°}$) contradicting Condition *monotonic* must be excluded.

| Choice of ($\Gamma_3^{320°}, \Gamma_3^{360°}$) | Family A | Family B |
|---|---|---|
| $\Gamma_3^{80°}$ | $\{o,h,c_1\}$, $\{o,h,c_2\}$, $\{o,h,c_1,c_2\}$; | $\{o,h,c_1\}$, $\{o,h,c_2\}$, $\{o,h,c_1,c_2\}$ |
| $\Gamma_3^{120°}$ | $\{o,h,c_1\}$, $\{o,h,c_2\}$, $\{o,h,c_1,c_2\}$; | $\{o,h,c_1\}$, $\{o,h,c_2\}$, $\{o,h,c_1,c_2\}$ |
| $\Gamma_3^{200°}$ | $\{o,h,c_1\}$, $\{o,h,c_1,c_2\}$ | $\{o,h,c_1\}$, $\{o,h,c_1,c_2\}$ |
| $\Gamma_3^{240°}$ | For $ohc_1^*c_2^*$-variants: $\{o,h\}$;<br>For $ohc_1^*c_2$-variants: $\{o,h\}$, $\{o,h,c_2\}$ | For $ohc_1^*c_2^*$-variants: $\{o\}$, $\{o,h\}$;<br>For $ohc_1^*c_2$-variants: $\{o\}$, $\{o,h\}$, $\{o,h,c_2\}$ |

To further reduce this number, we introduce an additional constraint, which we term Condition *asymmetry*, based on the asymmetry of $F_1$-ATPase structure. Considering the γ-subunit is intrinsically asymmetric in its structure, as it rotates within the stator ring, it should impose different steric restrictions on the three β-subunits at any orientation. Indeed, in virtually all available structures of $F_1$-ATPase (with intact γ-subunit), the three β-subunits are differentiated in conformational states, corroborating this asymmetry [32, 64, 65]. More quantitatively, tryptophan fluorescence

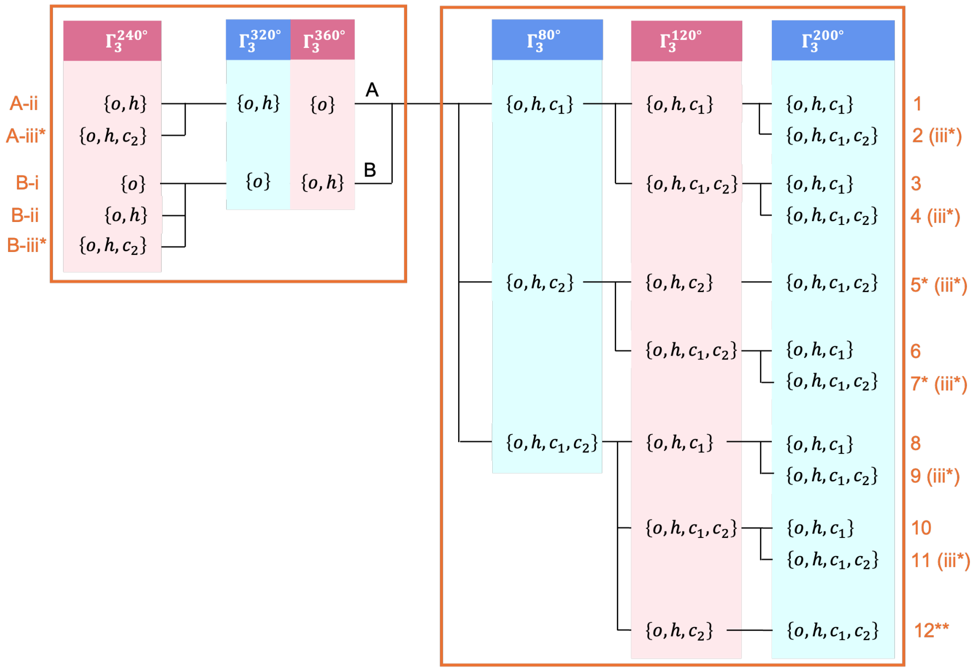


**Supplementary Fig. 15: Candidate $ohc_1c_2$-variants selected by Conditions *timing* I – IV, *structure*, and *monotonic***
Two $ohc_1^*c_2^*$-variants are omitted (5 and 7), because they are equivalent to the other two $ohc_1^*c_2^*$-variants (2 and 4) considering the equivalence of the $c_1$ and $c_2$ conformations.

experiments measuring nucleotide titration curves of *E coli.* $F_1$-ATPase suggest that the three catalytic sites are differentiated in nucleotide binding affinities [19, 21, 66, 67]; recent such experiments even demonstrate that the nucleotide binding affinities of the three catalytic sites are distinct when the $\gamma$-subunit is stalled at 80° or 120° [22]. Thus, we introduce Condition *asymmetry* that requires $\Gamma_k^\phi$ to be different for all the three $\beta$-subunits, which is formally equivalent to

$$\Gamma_3^{80^\circ} \neq \Gamma_3^{200^\circ}, \Gamma_3^{80^\circ} \neq \Gamma_3^{320^\circ}, \Gamma_3^{200^\circ} \neq \Gamma_3^{320^\circ},$$
$$\Gamma_3^{120^\circ} \neq \Gamma_3^{240^\circ}, \Gamma_3^{120^\circ} \neq \Gamma_3^{360^\circ}, \Gamma_3^{240^\circ} \neq \Gamma_3^{360^\circ}. \tag{46}$$

Applying this condition reduces the number of candidate $ohc_1c_2$-variants to 10 $ohc_1^*c_2^*$-variants + 20 $ohc_1^*c_2$-variants (Supplementary Figure 16).

#### Search for the minimal and most plausible $ohc_1c_2$-variants

These remaining 30 $ohc_1c_2$-variants are distributed equally between Family A and Family B (Supplementary Table 9). As reasoned in Results, we excluded all Family B variants based on thermodynamic constraints of product dissociation, and considered only the fifteen $ohc_1c_2$-variants in Family A.

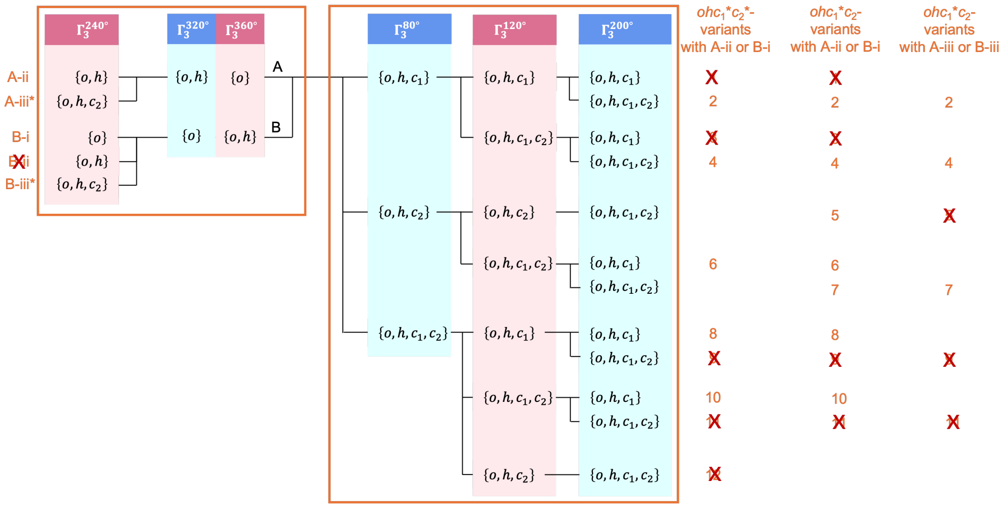


**Supplementary Fig. 16**: **Candidate** $ohc_1c_2$**-variants selected by Conditions** ***timing*** **I – IV,** ***structure*****,** ***monotonic*****, and** ***titration***

We classified these fifteen $ohc_1c_2$-variants into five groups (Supplementary Table 10) according to the number of Markov states included in each variant. From group 1 to 5, the number of included Markov states decreases from 972 to 702. We consider the variants including less Markov states more complex than those including less states, which may seem counterintuitive. However, this is because the variants including less Markov states assume stronger and/or more delicate interactions between the $\gamma$- and $\beta$-subunits (in the form of $\gamma$-$\beta$ restrictions), which could result in bias and over-selection of the states.

**Supplementary Table 10**: The finally remaining 5 $ohc_1^*c_2^*$-variants and 10 $ohc_1^*c_2$-variants, classified into five groups

| Group | Number of Markov states | Number of $ohc_1^*c_2^*$-variants | Number of $ohc_1^*c_2$-variants |
|---|---|---|---|
| 1 | 972 | 0 | 2 |
| 2 | 891 | 0 | 1 |
| 3 | 864 | 2 | 3 |
| 4 | 810 | 2 | 3 |
| 5 | 702 | 1 | 1 |

To find the minimal $ohc_1c_2$-variant that explains all available experimental data, we naturally started searching from the simplest variants. To decide whether the second closed conformation $c_2$ is also catalytically active, the selection covered both $ohc_1^*c_2^*$- and $ohc_1^*c_2$-variants ($\gamma$-$\beta$ restrictions shown in Table 3, second column). We first selected the two simplest $ohc_1^*c_2^*$-variants (from group 3, termed $ohc_1^*c_2^*$-$m.1$ and $ohc_1^*c_2^*$-$m.2$) for detailed investigation following the previously introduced training-validation procedure (Results). For direct comparison, we also investigated the two

$ohc_1^*c_2$-variants from group 3 with identical $\gamma$-$\beta$ restrictions. Additionally, we investigated one of the two simplest $ohc_1^*c_2$-variants (from group 1, termed $ohc_1^*c_2$-$w$). The performance of these two $ohc_1^*c_2^*$- and three $ohc_1^*c_2$-variants are reported in the main article (Results).

Notably, we consider these five selected $ohc_1c_2$-variants the most plausible ones. We did not extensively investigate the other ten $ohc_1c_2$-variants, but a brief analysis shows that they are either subsets of the selected $ohc_1c_2$-variants and therefore do not require extra testing, or less plausible and likely to disagree with some experimental data:

(1) The two $ohc_1c_2$-variants in group 5 (including one $ohc_1^*c_2^*$-variant and one $ohc_1^*c_2$-variant) with identical $\gamma$-$\beta$ restrictions (Supplementary Tables 11) are actually subsets of variants $ohc_1c_2$-$m.1$ or variants $ohc_1c_2$-$m.2$.

(2) The two $ohc_1^*c_2^*$-variants in group 4 and those two $ohc_1^*c_2$-variants with identical $\{\Gamma_{1,2,3}^{80°,120°}\}$ (Supplementary Table 12) are unlikely to reproduce the observed population shift from the 120°-states to the 80°-states with increasing [ATP]. With both $\{\Gamma_{1,2,3}^{80°,120°}\}$ shown in Supplementary Table 12, at 120°, all the three $\beta$-subunits are restricted from the most stable conformation ($c_2$ for the first set, $c_1$ for the second set). Therefore, it is likely that the 120°-states are of higher free energies than the 80°-states even at substantially low [ATP].

(3) The other four $ohc_1^*c_2$-variants from groups 1–4 respectively (Supplementary Table 13) are unlikely to reproduce the measured binding affinities of $F_1$-ATPase locked within the 80°-dwell [22]. In these four $ohc_1^*c_2$-variants, because only $c_1$ is catalytically active, the microscopic ATP binding affinity of $c_1$ should be higher than that of $c_2$. According to the analysis in Supplementary Section Supplementary Note 5, the apparent binding affinities of $\beta_2$ and $\beta_3$ in $F_1$-ATPase locked within the 80°-dwell are limited by the microscopic binding affinity of $c_1$, and therefore of similar magnitudes, contradicting the measurements showing three distinct binding affinities [22].

**Supplementary Table 11**: γ-β restrictions of the two $ohc_1c_2$-variants in group 5

| $\phi$ | β$_1$ | β$_2$ | β$_3$ |
|---|---|---|---|
| 80° | $o$, $h$ | $o$, $h$, $c_1$ | $o$, $h$, $c_2$ |
| 120° | $o$ | $o$, $h$ | $o$, $h$, $c_1$, $c_2$ |

**Supplementary Table 12**: γ-β restrictions of the four $ohc_1c_2$-variants in group 4

| $\phi$ | β$_1$ | β$_2$ | β$_3$ |
|---|---|---|---|
| 80° | $o$, $h$ | $o$, $h$, $c_1$ | $o$, $h$, $c_1$, $c_2$ |
| 120° | $o$ | $o$, $h$ | $o$, $h$, $c_1$ |

| $\phi$ | β$_1$ | β$_2$ | β$_3$ |
|---|---|---|---|
| 80° | $o$, $h$ | $o$, $h$, $c_1$, $c_2$ | $o$, $h$, $c_2$ |
| 120° | $o$ | $o$, $h$ | $o$, $h$, $c_2$ |

**Supplementary Table 13**: γ-β restrictions of the other four $ohc_1^*c_2$-variants from group 1–4 respectively

| $\phi$ | β$_1$ | β$_2$ | β$_3$ |
|---|---|---|---|
| 80° | $o$, $h$ | $o$, $h$, $c_1$, $c_2$ | $o$, $h$, $c_1$ |
| 120° | $o$ | $o$, $h$, $c_2$ | $o$, $h$, $c_1$, $c_2$ |

| $\phi$ | β$_1$ | β$_2$ | β$_3$ |
|---|---|---|---|
| 80° | $o$, $h$ | $o$, $h$, $c_1$, $c_2$ | $o$, $h$, $c_1$ |
| 120° | $o$ | $o$, $h$, $c_2$ | $o$, $h$, $c_1$ |

| $\phi$ | β$_1$ | β$_2$ | β$_3$ |
|---|---|---|---|
| 80° | $o$, $h$ | $o$, $h$, $c_1$, $c_2$ | $o$, $h$, $c_1$ |
| 120° | $o$ | $o$, $h$ | $o$, $h$, $c_1$, $c_2$ |

| $\phi$ | β$_1$ | β$_2$ | β$_3$ |
|---|---|---|---|
| 80° | $o$, $h$ | $o$, $h$, $c_1$, $c_2$ | $o$, $h$, $c_1$ |
| 120° | $o$ | $o$, $h$ | $o$, $h$, $c_1$ |

### Supplementary Note 5 Analytical expressions of the apparent ADP binding affinities in terms of the model parameters

Here, we derive analytical expressions for the apparent ADP binding affinities (Methods) in terms of the parameters of our Markov model, particularly the free energies. The apparent binding affinities $\{K_k\}$ are obtained by fitting an ADP titration curve to the equilibrium binding model [20–22]:

$$\nu(c_\mathrm{D}) = \sum_{k=1}^{3} \frac{c_\mathrm{D}}{c_\mathrm{D} + K_k}, \tag{47}$$

where $\nu$ is the total occupancy of the binding sites; $c_\mathrm{D}$ is the concentration of ADP; the summation is over all the three binding sites within $F_1$-ATPase. Our aim is to derive an expression for $\nu$ in terms of the free energies as well as ADP concentration $c_\mathrm{D}$ which will then be compared to Supplementary Equation (47) to obtain the expressions of $\{K_k\}$.

Within our Markov model, the total occupancy $\nu$ is evaluated by the ensemble average of the numbers of occupied catalytic sites $\{n_i\}$ of the Markov states $\{i\}$ in the steady-state of the system where the populations of the states are $\{\rho_i^\mathrm{st}\}$ (Equation (11) and Supplementary Equation (19)). To obtain the steady-state populations analytically, we use the Boltzmann distribution approximation. For ADP titration, ADP concentration is varied in a large range, whereas ATP concentration is typically low. Under this condition, the net rate of ATP hydrolysis/synthesis is typically small, and the populations of the Markov states with ATP-bound β-subunits (which result from reversible ATP synthesis in the catalytically active sites) are also low and can be ignored. Therefore, the steady-state distribution is expected to approach the Boltzmann distribution of those Markov states including only empty and ADP-bound β-subunits, where ADP binding and dissociation within each β-subunit is in equilibrium.

In the Boltzmann distribution, the population of a state $i$ including only empty and/or ADP-bound β-subunits depends on its free energy $G_i$:

$$\rho_i = \frac{1}{Z} \exp(-G_i/k_\mathrm{B}T), \tag{48}$$

where $T$ is the temperature and $Z$ is the partition function for normalizing the distribution,

$$Z = \sum_i \exp(-G_i/k_\mathrm{B}T), \tag{49}$$

where the summation is over all these states. Due to the three-fold pseudo-symmetry of $F_1$-ATPase, we only need to consider those states where the $\gamma$-subunit is at 80° or 120°.

For these states, the free energies are (adapted from Equations (2)-(3))

$$G(\mathbf{s}, c_\mathrm{D}) = \sum_{k=1,2,3} G_{\beta,\mathcal{C}^{(k)}} + \sum_{\{k|\mathcal{B}^{(k)}=\mathrm{D}\}} \left(\Delta G^{\ominus}_{\mathrm{b},\mathcal{C}^{(k)},\mathrm{D}} - k_\mathrm{B}T \ln c_\mathrm{D}\right) + G_{\gamma\beta}(\phi), \tag{50}$$

where $G_{\gamma\beta} = -2.3\, k_\mathrm{B}T$ if and only if the $\gamma$-subunit is at 120° ($\phi = 120°$; at this orientation, $\beta_1$ can only adopt the *o* conformation); otherwise, $G_{\gamma\beta} = 0$. The second term in Supplementary Equation (50) includes summation only over the ADP-bound $\beta$-subunit(s). The contribution to the partition function by this state is

$$\begin{aligned} Z(\mathbf{s}, c_\mathrm{D}) &= \exp\left(-\frac{G(\mathbf{s}, c_\mathrm{D})}{k_\mathrm{B}T}\right) \\ &= \exp\left(-\frac{G^{\ominus}(\mathbf{s})}{k_\mathrm{B}T}\right) c_\mathrm{D}^{b_1+b_2+b_3} \\ &\equiv Z^{\ominus}(\mathbf{s}) c_\mathrm{D}^{b_1+b_2+b_3}. \end{aligned} \tag{51}$$

Here, $b_k = 0$ and 1 ($k = 1, 2, 3$) for empty and ADP-bound $\beta_k$ respectively; $G^{\ominus}(\mathbf{s})$ is the [ADP]-independent part $G^{\ominus}(\mathbf{s})$ of the free energy of the state $\mathbf{s}$:

$$G^{\ominus}(\mathbf{s}) = \sum_{k=1,2,3} G_{\beta,\mathcal{C}^{(k)}} + \sum_{\{k|\mathcal{B}^{(k)}=\mathrm{D}\}} \Delta G^{\ominus}_{\mathrm{b},\mathcal{C}^{(k)},\mathrm{D}} + G_{\gamma\beta}(\phi), \tag{52}$$

and $Z^{\ominus}(\mathbf{s})$ is defined

$$Z^{\ominus}(\mathbf{s}) = \exp(-G^{\ominus}(\mathbf{s})/k_\mathrm{B}T). \tag{53}$$

The total occupancy $\nu$ is (adapted from Supplementary Equation (19)):

$$\nu(c_\mathrm{D}) = \sum_{\mathbf{s}} b(\mathbf{s})\rho(\mathbf{s}), \tag{54}$$

where $b(\mathbf{s}) = (b_1 + b_2 + b_3)$ is the number of occupied binding sites in state $\mathbf{s}$. Accordingly, $\nu$ is expressed in terms of $Z^{\ominus}(\mathbf{s})$ by:

$$\begin{aligned} \nu(c_\mathrm{D}) &= \frac{\sum_{\mathbf{s}} b(\mathbf{s}) Z^{\ominus}(\mathbf{s}) c_\mathrm{D}^{b(\mathbf{s})}}{\sum_{\mathbf{s}} Z^{\ominus}(\mathbf{s}) c_\mathrm{D}^{b(\mathbf{s})}} \\ &= \frac{\sum\limits_{b(\mathbf{s})=1} Z^{\ominus}(\mathbf{s}) c_\mathrm{D} + 2 \sum\limits_{b(\mathbf{s})=2} Z^{\ominus}(\mathbf{s}) c_\mathrm{D}^2 + 3 \sum\limits_{b(\mathbf{s})=3} Z^{\ominus}(\mathbf{s}) c_\mathrm{D}^3}{\sum\limits_{b(\mathbf{s})=0} Z^{\ominus}(\mathbf{s}) + \sum\limits_{b(\mathbf{s})=1} Z^{\ominus}(\mathbf{s}) c_\mathrm{D} + \sum\limits_{b(\mathbf{s})=2} Z^{\ominus}(\mathbf{s}) c_\mathrm{D}^2 + \sum\limits_{b(\mathbf{s})=3} Z^{\ominus}(\mathbf{s}) c_\mathrm{D}^3} \\ &\equiv \frac{W_1 c_\mathrm{D} + 2W_2 c_\mathrm{D}^2 + 3W_3 c_\mathrm{D}^3}{W_0 + W_1 c_\mathrm{D} + W_2 c_\mathrm{D}^2 + W_3 c_\mathrm{D}^3}, \end{aligned} \tag{55}$$

with

$$\begin{aligned} W_0 &= \sum_{b(\mathbf{s})=0} Z^{\Theta}(\mathbf{s}) = \sum_{b_{1,2,3}=0} Z^{\Theta}(\mathbf{s}), \\ W_1 &= \sum_{b(\mathbf{s})=1} Z^{\Theta}(\mathbf{s}), \\ W_2 &= \sum_{b(\mathbf{s})=2} Z^{\Theta}(\mathbf{s}), \\ W_3 &= \sum_{b(\mathbf{s})=3} Z^{\Theta}(\mathbf{s}) = \sum_{b_{1,2,3}=1} Z^{\Theta}(\mathbf{s}). \end{aligned} \tag{56}$$

Here, " $\sum_{b(\mathbf{s})=m}$ " $(m = 0, 1, 2, 3)$ represents the summation over all the states $\mathbf{s}$ where $b(\mathbf{s}) = m$.

Supplementary Equation (47) for $N = 3$ can be written as

$$\nu(c_{\rm D}) = \frac{\left(\sum_{j<k} K_j K_k\right) c_{\rm D} + 2\left(\sum_k K_k\right) c_{\rm D}^2 + 3c_{\rm D}^3}{K_1K_2K_3 + \left(\sum_{j<k} K_j K_k\right) c_{\rm D} + \left(\sum_k K_k\right) c_{\rm D}^2 + c_{\rm D}^3}, \tag{57}$$

with $j, k = 1, 2, 3$. Comparing Supplementary Equation (55) to Supplementary Equation (57), it is found that

$$\begin{aligned} K_1K_2K_3 &= W_0/W_3, \\ K_1K_2 + K_1K_3 + K_2K_3 &= W_1/W_3, \\ K_1 + K_2 + K_3 &= W_2/W_3, \end{aligned} \tag{58}$$

which implies that the apparent binding affinities $K_1$, $K_2$ and $K_3$ are the three roots of the cubic equation

$$W_3x^3 - W_2x^2 + W_1x - W_0 = 0. \tag{59}$$

Notably, the physical meaning of the binding affinities require $K_1$, $K_2$ and $K_3$ to be positive real numbers, whereas there is no guarantee (at least not clear yet) that Supplementary Equation (59) has three real roots. If Supplementary Equation (59) does have three real roots, these three roots must be positive considering Supplementary Equation (56) which shows $W_{0,1,2,3} > 0$ and Supplementary Equation (58). In this case, Supplementary Equation (58) provides a general analytical solution for the apparent binding affinities obtained by fitting the ADP titration curve predicted by our model to Supplementary Equation (47).

However, if Supplementary Equation (59) does not have three real roots, it must have one real positive root and two conjugate complex roots. In this case, the apparent binding affinities obtained by fitting the predicted ADP titration curve to Supplementary Equation (47) are likely different from those given by Supplementary Equation (58).

While Supplementary Equation (58) is general, it is rather complicated and unstraightforward. Below, we attempt to derive a more specific but simpler and easier-to-interpret expression for the apparent binding affinities.

Plug Supplementary Equations (53) and (52) into Supplementary Equation (56), we obtain

$$\begin{aligned}
W_0 &= \sum_{(\phi,\mathcal{C}^{(1)},\mathcal{C}^{(2)},\mathcal{C}^{(3)})} Z^{\mathrm{conf}}, \\
W_1 &= \sum_{(\phi,\mathcal{C}^{(1)},\mathcal{C}^{(2)},\mathcal{C}^{(3)})} Z^{\mathrm{conf}} \sum_{k=1,2,3} \exp\left(-\frac{\Delta G^{\ominus}_{\mathrm{b},\mathcal{C}^{(k)},\mathrm{D}}}{k_{\mathrm{B}}T}\right), \\
W_2 &= \sum_{(\phi,\mathcal{C}^{(1)},\mathcal{C}^{(2)},\mathcal{C}^{(3)})} Z^{\mathrm{conf}} \sum_{j<k} \exp\left(-\frac{\Delta G^{\ominus}_{\mathrm{b},\mathcal{C}^{(j)},\mathrm{D}}}{k_{\mathrm{B}}T}\right) \exp\left(-\frac{\Delta G^{\ominus}_{\mathrm{b},\mathcal{C}^{(k)},\mathrm{D}}}{k_{\mathrm{B}}T}\right), \\
W_3 &= \sum_{(\phi,\mathcal{C}^{(1)},\mathcal{C}^{(2)},\mathcal{C}^{(3)})} Z^{\mathrm{conf}} \prod_{k=1,2,3} \exp\left(-\frac{\Delta G^{\ominus}_{\mathrm{b},\mathcal{C}^{(k)},\mathrm{D}}}{k_{\mathrm{B}}T}\right),
\end{aligned} \tag{60}$$

where the summation "$\sum_{(\phi,\mathcal{C}^{(1)},\mathcal{C}^{(2)},\mathcal{C}^{(3)})}$" is over all the possible combinations of γ-orientations ($\phi = 80°, 120°$) and β-subunit conformations ($\mathcal{C}^{(k)} \in \Gamma^{\phi}_{k}$, $k = 1, 2, 3$); $Z^{\mathrm{conf}}$ is defined

$$Z^{\mathrm{conf}}(\phi, \mathcal{C}^{(1)}, \mathcal{C}^{(2)}, \mathcal{C}^{(3)}) = \exp\left(-\frac{G_{\gamma\beta}(\phi)}{k_{\mathrm{B}}T}\right) \prod_{k=1,2,3} \exp\left(-\frac{G_{\beta,\mathcal{C}^{(k)}}}{k_{\mathrm{B}}T}\right). \tag{61}$$

For simplicity of the equations to be presented further, we define the following equilibrium constants

$$\begin{aligned}
K_{\mathrm{I},\phi} &= \exp\left(-\frac{G_{\gamma\beta}(\phi)}{k_{\mathrm{B}}T}\right), \\
K_{\beta,\mathcal{C}} &= \exp\left(-\frac{G_{\beta,\mathcal{C}}}{k_{\mathrm{B}}T}\right), \\
K_{\mathrm{b},\mathcal{C}} &= \exp\left(-\frac{\Delta G^{\ominus}_{\mathrm{b},\mathcal{C}}}{k_{\mathrm{B}}T}\right),
\end{aligned} \tag{62}$$

for $\mathcal{C}$ being all the β-subunit conformations included in the variant, *i.e.*, $\mathcal{C} = o, h, c$ for *ohc*-variants, and $\mathcal{C} = o, h, c_1, c_2$ for *ohc*$_1$*c*$_2$-variants.

**$F_1$-ATPase locked in 80°- or 120°-dwell**

We first consider a simpler case where $F_1$-ATPase is locked in the catalytic dwell at 80°, *e.g.*, by disulfide crosslinking [22]. In this case, $\phi$ can only be 80°, while each of the three β-subunits could freely and independently choose any conformation $\mathcal{C}^{(k)} \in \Gamma_k^{80°}$, and the summation $\sum\limits_{(\phi,\mathcal{C}^{(1)},\mathcal{C}^{(2)},\mathcal{C}^{(3)})}$ can be dissected into the triple sum $\sum\limits_{\mathcal{C}^{(1)}}\sum\limits_{\mathcal{C}^{(2)}}\sum\limits_{\mathcal{C}^{(3)}}$ omitting $\phi$. Additionally, $K_{\mathrm{I},\phi} = 1$. Accordingly, and using the definitions in Supplementary Equation (62), $W_0$ and $W_1$ in Supplementary Equation (60) are converted to

$$W_0 = \prod_{k=1,2,3}\left(\sum_{\mathcal{C}^{(k)}} K_{\beta,\mathcal{C}^{(k)}}\right),$$
$$W_1 = \sum_{(i,j,k)}\left[\left(\sum_{\mathcal{C}^{(i)}} K_{\beta,\mathcal{C}^{(i)}} K_{\mathrm{b},\mathcal{C}^{(i)}}\right)\left(\sum_{\mathcal{C}^{(j)}} K_{\beta,\mathcal{C}^{(j)}}\right)\left(\sum_{\mathcal{C}^{(k)}} K_{\beta,\mathcal{C}^{(k)}}\right)\right], \tag{63}$$

where the summation $\sum\limits_{(i,j,k)}$ is over $(i,j,k) \in \{(1,2,3),(2,1,3),(3,1,2)\}$. Accordingly,

$$\frac{W_1}{W_0} = \frac{\sum\limits_{\mathcal{C}^{(1)}} K_{\beta,\mathcal{C}^{(1)}} K_{\mathrm{b},\mathcal{C}^{(1)}}}{\sum\limits_{\mathcal{C}^{(1)}} K_{\beta,\mathcal{C}^{(1)}}} + \frac{\sum\limits_{\mathcal{C}^{(2)}} K_{\beta,\mathcal{C}^{(2)}} K_{\mathrm{b},\mathcal{C}^{(2)}}}{\sum\limits_{\mathcal{C}^{(2)}} K_{\beta,\mathcal{C}^{(2)}}} + \frac{\sum\limits_{\mathcal{C}^{(3)}} K_{\beta,\mathcal{C}^{(3)}} K_{\mathrm{b},\mathcal{C}^{(3)}}}{\sum\limits_{\mathcal{C}^{(3)}} K_{\beta,\mathcal{C}^{(3)}}}. \tag{64}$$

Comparing this Supplementary Equation (64) to $W_1/W_0 = 1/K_1 + 1/K_2 + 1/K_3$ (given by Supplementary Equation (58)), each term in Supplementary Equation (64) is attributed to the apparent binding affinity of one β-subunit,

$$\frac{1}{K_k} = \frac{\sum\limits_{\mathcal{C}\in\Gamma_k^{80°}} K_{\beta,\mathcal{C}} K_{\mathrm{b},\mathcal{C}}}{\sum\limits_{\mathcal{C}\in\Gamma_k^{80°}} K_{\beta,\mathcal{C}}}. \tag{65}$$

It is easy to verify that $K_k$ given in Supplementary Equation (65) satisfy Supplementary Equation (58).

Supplementary Equation (65) can be written in a more symmetric form

$$(K_k^{80°})^{-1} = \frac{1}{\sum\limits_{\mathcal{C}\in\Gamma_k^{80°}} K_{\beta,\mathcal{C}}} \sum_{\mathcal{C}\in\Gamma_k^{80°}} K_{\beta,\mathcal{C}} (K_{\mathrm{d},\mathcal{C}})^{-1}, \tag{66}$$

where $K_{\mathrm{d},\mathcal{C}} = \exp(G_{\mathrm{b},\mathcal{C}}/k_{\mathrm{B}}T)$. Here, we use the superscript 80° in the apparent binding affinity $K_k^{80°}$ to emphasize that this equation is derived for $F_1$-ATPase stalled at 80°.

In contrast to $K_{1,2,3}$ which are interpreted as the *apparent* binding affinities (dissociation constants) of the β-subunits, $K_{\mathrm{d},\mathcal{C}}$ including $K_{\mathrm{d},o}$, $K_{\mathrm{d},h}$ and $K_{\mathrm{d},c}$ (in *ohc*-variants) or $K_{\mathrm{d},o}$, $K_{\mathrm{d},h}$, $K_{\mathrm{d},c_1}$ and $K_{\mathrm{d},c_2}$ (in $ohc_1c_2$-variants) can be interpreted as the *microscopic* binding affinities (dissociation constants) of the β-subunit conformations included in our Markov model. According to this Supplementary Equation (66), the *apparent* binding affinity of a β-subunit is a superposition (weighted average) of the *microscopic* binding affinities of the several conformations this β-subunit can adopt, with weights $K_{\beta,\mathcal{C}}$ as the Boltzmann factors of the conformational energies $G_{\beta,\mathcal{C}}$ (Supplementary Equation (62)).

For $F_1$-ATPase locked in the ATP-waiting dwell ($\phi = 120°$ exclusively), similar derivation can be done, which leads to

$$(K_k^{120°})^{-1} = \frac{1}{\sum\limits_{\mathcal{C}\in\Gamma_k^{120°}} K_{\beta,\mathcal{C}}} \sum_{\mathcal{C}\in\Gamma_k^{120°}} K_{\beta,\mathcal{C}}(K_{\mathrm{d},\mathcal{C}})^{-1}, \tag{67}$$

of exactly the same form with Supplementary Equation (66) derived for $F_1$-ATPase stalled at 80°. The additional favorable interaction energy between the γ- and $\beta_1$-subunit is cancelled-out and therefore does not appear in Supplementary Equation (67). Notably, because we assumed $\Gamma_1^{120°} = \{o\}$, the apparent binding affinity of $\beta_1$ in $F_1$-ATPase stalled at 120° is simply the microscopic binding affinity of the $o$ conformation.

Supplementary Equations (66) and (67) can be summarized into one equation:

$$(K_k^{\phi})^{-1} = \frac{1}{\Omega_k} \sum_{\mathcal{C}\in\Gamma_k^{\phi}} K_{\beta,\mathcal{C}}(K_{\mathrm{d},\mathcal{C}})^{-1}, \text{with}\, \Omega_k = \sum_{\mathcal{C}\in\Gamma_k^{\phi}} K_{\beta,\mathcal{C}}. \tag{68}$$

This equation can be generalized to freely-rotating $F_1$-ATPase by defining the concept of apparent binding affinity $K_k^{\phi}$ of the $\beta_k$-subunit at a particular γ-orientation $\phi$, which provides a simplified representation of the set of accessible conformations of the $\beta_k$-subunit at $\phi$ (*i.e.*, $\Gamma_k^{\phi}$), and, can be considered as a measure for the "open-ness" of $\beta_k$.

An important conclusion from Supplementary Equation (68) is that if two β-subunits are allowed to adopt the same set of conformations at a γ-orientation $\phi$ (*i.e.*, $\Gamma_k^{\phi} = \Gamma_j^{\phi}$), their apparent binding affinities at this γ-orientation are likely very similar. The ADP titration curves measured on $F_1$-ATPase locked into the catalytic and ATP-waiting dwells suggest three distinct binding affinities within both dwells [22]. Therefore, we excluded those $ohc_1c_2$-variants assuming identical γ-β restrictions $\Gamma_k^{\phi}$ for two β-subunits at the same γ-orientation (Supplementary Note 3).

#### Freely-rotating $F_1$-ATPase

For freely-rotating $F_1$-ATPase, the summations over $\phi$ in $W_0$ and $W_1$ (Supplementary Equation (60)) must be preserved, due to which the factorization of the three β-subunits like in Supplementary Equation (63) is not always possible. However, if the γ-β restrictions for a β-subunit are identical at 80° and 120° (*e.g.*, for $\beta_k$, $\Gamma_k^{80°} = \Gamma_k^{120°}$), the summations over the conformations of this β-subunit in $W_0$ and $W_1$ can be factorized out:

$$W_0 = \left(\sum_{\mathcal{C}^{(k)}} K_{\beta,\mathcal{C}^{(k)}}\right)\left(\sum_{(\phi,\mathcal{C}^{(i)},\mathcal{C}^{(j)})} K_{\mathrm{I},\phi} K_{\beta,\mathcal{C}^{(i)}} K_{\beta,\mathcal{C}^{(j)}}\right), \tag{69}$$

$$\begin{aligned} W_1 = &\left(\sum_{\mathcal{C}^{(k)}} K_{\beta,\mathcal{C}^{(k)}} K_{\mathrm{b},\mathcal{C}^{(k)}}\right)\left(\sum_{(\phi,\mathcal{C}^{(i)},\mathcal{C}^{(j)})} K_{\mathrm{I},\phi} K_{\beta,\mathcal{C}^{(i)}} K_{\beta,\mathcal{C}^{(j)}}\right) \\ &+ \left(\sum_{\mathcal{C}^{(k)}} K_{\beta,\mathcal{C}^{(k)}}\right)\left(\sum_{(\phi,\mathcal{C}^{(i)},\mathcal{C}^{(j)})} K_{\mathrm{I},\phi} K_{\beta,\mathcal{C}^{(i)}} K_{\beta,\mathcal{C}^{(j)}} (K_{\mathrm{b},\mathcal{C}^{(i)}} + K_{\mathrm{b},\mathcal{C}^{(j)}})\right), \end{aligned} \tag{70}$$

thus the ratio is

$$\frac{W_1}{W_0} = \frac{\sum_{\mathcal{C}^{(k)}} K_{\beta,\mathcal{C}^{(k)}} K_{\mathrm{b},\mathcal{C}^{(k)}}}{\sum_{\mathcal{C}^{(k)}} K_{\beta,\mathcal{C}^{(k)}}} + \frac{\sum_{(\phi,\mathcal{C}^{(i)},\mathcal{C}^{(j)})} K_{\mathrm{I},\phi} K_{\beta,\mathcal{C}^{(i)}} K_{\beta,\mathcal{C}^{(j)}} (K_{\mathrm{b},\mathcal{C}^{(i)}} + K_{\mathrm{b},\mathcal{C}^{(j)}})}{\sum_{(\phi,\mathcal{C}^{(i)},\mathcal{C}^{(j)})} K_{\mathrm{I},\phi} K_{\beta,\mathcal{C}^{(i)}} K_{\beta,\mathcal{C}^{(j)}}}, \tag{71}$$

and we attribute the first term in Supplementary Equation (71) to $K_k$:

$$(K_k)^{-1} = \frac{1}{\Omega_k} \sum_{\mathcal{C}\in\Gamma_k^{\phi}} K_{\beta,\mathcal{C}} (K_{\mathrm{d},\mathcal{C}})^{-1}, \text{with}\, \Omega_k = \sum_{\mathcal{C}\in\Gamma_k^{\phi}} K_{\beta,\mathcal{C}}, \tag{72}$$

where $\Gamma_k^{\phi} = \Gamma_k^{80°} = \Gamma_k^{120°}$. We say that this $\beta_k$-subunit is "decoupled" whereas the other two β-subunit are "coupled" due to the indirect interaction mediated via the γ-subunit,

$$\begin{aligned} (K_i)^{-1} + (K_j)^{-1} &= \frac{1}{\Omega_{(i,j)}} \sum_{(\phi,\mathcal{C}^{(i)},\mathcal{C}^{(j)})} \omega_{\phi,\mathcal{C}^{(i)},\mathcal{C}^{(j)}} \left((K_{\mathrm{d},\mathcal{C}^{(i)}})^{-1} + (K_{\mathrm{d},\mathcal{C}^{(j)}})^{-1}\right), \\ (K_i)^{-1}(K_j)^{-1} &= \frac{1}{\Omega_{(i,j)}} \sum_{(\phi,\mathcal{C}^{(i)},\mathcal{C}^{(j)})} \omega_{\phi,\mathcal{C}^{(i)},\mathcal{C}^{(j)}} (K_{\mathrm{d},\mathcal{C}^{(i)}})^{-1} (K_{\mathrm{d},\mathcal{C}^{(j)}})^{-1}, \\ \text{with} \quad \omega_{\phi,\mathcal{C}^{(i)},\mathcal{C}^{(j)}} &= K_{\mathrm{I},\phi} K_{\beta,\mathcal{C}^{(i)}} K_{\beta,\mathcal{C}^{(j)}}, \quad \Omega_{(i,j)} = \sum_{(\phi,\mathcal{C}^{(i)},\mathcal{C}^{(j)})} \omega_{\phi,\mathcal{C}^{(i)},\mathcal{C}^{(j)}}. \end{aligned} \tag{73}$$

If the γ-β restrictions for another β-subunit are also identical at 80° and 120° (*e.g.*, for $\beta_{\mathrm{j}}$, $\Gamma_j^{80^\circ} = \Gamma_j^{120^\circ}$), the summations over the conformations of this β-subunit in $W_0$ and $W_1$ can be further factorized out, *i.e.*, Equations (69) and (70) can be further converted to

$$W_0 = \left(\sum_{\mathcal{C}^{(k)}} K_{\beta,\mathcal{C}^{(k)}}\right)\left(\sum_{\mathcal{C}^{(j)}} K_{\beta,\mathcal{C}^{(j)}}\right)\left(\sum_{(\phi,\mathcal{C}^{(i)})} K_{\mathrm{I},\phi} K_{\beta,\mathcal{C}^{(i)}}\right), \tag{74}$$

$$\begin{aligned} W_1 = &\left(\sum_{\mathcal{C}^{(k)}} K_{\beta,\mathcal{C}^{(k)}} K_{\mathrm{b},\mathcal{C}^{(k)}}\right)\left(\sum_{\mathcal{C}^{(j)}} K_{\beta,\mathcal{C}^{(j)}}\right)\left(\sum_{(\phi,\mathcal{C}^{(i)})} K_{\mathrm{I},\phi} K_{\beta,\mathcal{C}^{(i)}}\right) \\ &+ \left(\sum_{\mathcal{C}^{(k)}} K_{\beta,\mathcal{C}^{(k)}}\right)\left(\sum_{\mathcal{C}^{(j)}} K_{\beta,\mathcal{C}^{(j)}} K_{\mathrm{b},\mathcal{C}^{(j)}}\right)\left(\sum_{(\phi,\mathcal{C}^{(i)})} K_{\mathrm{I},\phi} K_{\beta,\mathcal{C}^{(i)}}\right) \\ &+ \left(\sum_{\mathcal{C}^{(k)}} K_{\beta,\mathcal{C}^{(k)}}\right)\left(\sum_{\mathcal{C}^{(j)}} K_{\beta,\mathcal{C}^{(j)}}\right)\left(\sum_{(\phi,\mathcal{C}^{(i)})} K_{\mathrm{I},\phi} K_{\beta,\mathcal{C}^{(i)}} K_{\mathrm{b},\mathcal{C}^{(i)}}\right), \end{aligned} \tag{75}$$

thus the ratio is

$$\frac{W_1}{W_0} = \frac{\sum_{\mathcal{C}^{(k)}} K_{\beta,\mathcal{C}^{(k)}} K_{\mathrm{b},\mathcal{C}^{(k)}}}{\sum_{\mathcal{C}^{(k)}} K_{\beta,\mathcal{C}^{(k)}}} + \frac{\sum_{\mathcal{C}^{(j)}} K_{\beta,\mathcal{C}^{(j)}} K_{\mathrm{b},\mathcal{C}^{(j)}}}{\sum_{\mathcal{C}^{(j)}} K_{\beta,\mathcal{C}^{(j)}}} + \frac{\sum_{(\phi,\mathcal{C}^{(i)})} K_{\mathrm{I},\phi} K_{\beta,\mathcal{C}^{(i)}} K_{\mathrm{b},\mathcal{C}^{(i)}}}{\sum_{(\phi,\mathcal{C}^{(i)})} K_{\mathrm{I},\phi} K_{\beta,\mathcal{C}^{(i)}}}, \tag{76}$$

and we attribute the three terms in Supplementary Equation (76) to $K_k$, $K_j$ and $K_i$ respectively:

$$(K_i)^{-1} = \frac{1}{\Omega_i} \sum_{(\phi,\mathcal{C}^{(i)})} K_{\mathrm{I},\phi} K_{\beta,\mathcal{C}^{(i)}} (K_{\mathrm{d},\mathcal{C}^{(i)}})^{-1}, \text{with } \Omega_i = \sum_{(\phi,\mathcal{C}^{(i)})} K_{\mathrm{I},\phi} K_{\beta,\mathcal{C}^{(i)}}, \tag{77}$$

$$\begin{aligned} (K_j)^{-1} &= \frac{1}{\Omega_j} \sum_{\mathcal{C}\in\Gamma_j^{\phi}} K_{\beta,\mathcal{C}} (K_{\mathrm{d},\mathcal{C}})^{-1}, \text{with } \Omega_j = \sum_{\mathcal{C}\in\Gamma_j^{\phi}} K_{\beta,\mathcal{C}}, \\ (K_k)^{-1} &= \frac{1}{\Omega_k} \sum_{\mathcal{C}\in\Gamma_k^{\phi}} K_{\beta,\mathcal{C}} (K_{\mathrm{d},\mathcal{C}})^{-1}, \text{with } \Omega_k = \sum_{\mathcal{C}\in\Gamma_k^{\phi}} K_{\beta,\mathcal{C}}. \end{aligned} \tag{78}$$

where $\Gamma_j^{\phi} = \Gamma_j^{80^\circ} = \Gamma_j^{120^\circ}$ and $\Gamma_k^{\phi} = \Gamma_k^{80^\circ} = \Gamma_k^{120^\circ}$. Here, all the three β-subunits are "decoupled".

More explicitly, Supplementary Equation (77) is

$$(K_i)^{-1} = \frac{1}{\Omega_i} \left( \sum_{\mathcal{C}\in\Gamma_i^{80^\circ}} K_{\beta,\mathcal{C}} (K_{\mathrm{d},\mathcal{C}})^{-1} + K_{\mathrm{I}} \sum_{\mathcal{C}\in\Gamma_i^{120^\circ}} K_{\beta,\mathcal{C}} (K_{\mathrm{d},\mathcal{C}})^{-1} \right), \tag{79}$$

where $K_{\mathrm{I}} \equiv K_{\mathrm{I},120^\circ} = 10$, and

$$\Omega_i = \sum_{\mathcal{C}\in\Gamma_i^{80^\circ}} K_{\beta,\mathcal{C}} + K_{\mathrm{I}} \sum_{\mathcal{C}\in\Gamma_i^{120^\circ}} K_{\beta,\mathcal{C}}. \tag{80}$$

Notably, the additional favorable interaction energy between the $\gamma$- and $\beta_1$-subunits takes effect in the apparent binding affinity of $\beta_i$, the only $\beta$-subunit whose $\Gamma_i^{80^\circ} \neq \Gamma_i^{120^\circ}$, as an extra weight $K_{\mathrm{I}}$ on the 120°-conformations, even though $\beta_i$ may or may not be $\beta_1$.

#### Application to particular variants

These analytical expressions of the apparent ADP binding affinities in terms of the model parameters explain why variant *ohc-s* reproduces the measured ADP binding affinities [21] whereas variant *ohc-w* does not (Results).

For both variants *ohc-s* and *ohc-w*, the three $\beta$-subunits are *decoupled*. For variant *ohc-w*, applying Equations (78) and (79),

$$\begin{aligned}
(K_1)^{-1} =& \frac{1}{\Omega_1}\left((1+K_{\mathrm{I}})K_{\beta,o}(K_{\mathrm{d},o})^{-1} + K_{\beta,h}(K_{\mathrm{d},h})^{-1}\right), \\
&\text{with} \quad \Omega_1 = (1+K_{\mathrm{I}})K_{\beta,o} + K_{\beta,h}; \qquad (81)\\
(K_2)^{-1} = (K_3)^{-1} =& \frac{1}{\Omega_{2,3}}\left(K_{\beta,o}(K_{\mathrm{d},o})^{-1} + K_{\beta,h}(K_{\mathrm{d},h})^{-1} + K_{\beta,c}(K_{\mathrm{d},c})^{-1}\right), \\
&\text{with} \quad \Omega_{2,3} = K_{\beta,o} + K_{\beta,h} + K_{\beta,c}. \qquad (82)
\end{aligned}$$

Supplementary Equation (82) suggests that the apparent binding affinities of $\beta_2$ and $\beta_3$ are likely very close, which has been corroborated by the numerical results (Supplementary Fig. 1).

In contrast, for variant *ohc-s*,

$$\begin{aligned}
(K_1)^{-1} =& (K_{\mathrm{d},o})^{-1}; \\
(K_2)^{-1} =& \frac{1}{\Omega_2}\left((1+K_{\mathrm{I}})K_{\beta,o}(K_{\mathrm{d},o})^{-1} + K_{\beta,h}(K_{\mathrm{d},h})^{-1}\right), \\
&\text{with} \quad \Omega_2 = (1+K_{\mathrm{I}})K_{\beta,o} + K_{\beta,h}; \qquad (83)\\
(K_3)^{-1} =& \frac{1}{\Omega_3}\left(K_{\beta,o}(K_{\mathrm{d},o})^{-1} + K_{\beta,h}(K_{\mathrm{d},h})^{-1} + K_{\beta,c}(K_{\mathrm{d},c})^{-1}\right), \\
&\text{with} \quad \Omega_3 = K_{\beta,o} + K_{\beta,h} + K_{\beta,c}. \qquad (84)
\end{aligned}$$

It is therefore possible to find parameter sets for variant *ohc-s* to reproduce three distinct apparent binding affinities in agreement with the measured ones [21] (Figure 2c).

However, for variant *ohc-m*, only $\beta_3$ is decoupled ($\Gamma_3^{80^\circ} = \Gamma_3^{120^\circ} = \{o, h, c\}$),

$$(K_3)^{-1} = \frac{1}{\Omega_3}\left(K_{\beta,o}(K_{\mathrm{d},o})^{-1} + K_{\beta,h}(K_{\mathrm{d},h})^{-1} + K_{\beta,c}(K_{\mathrm{d},c})^{-1}\right),$$
$$\text{with} \quad \Omega_3 = K_{\beta,o} + K_{\beta,h} + K_{\beta,c}; \tag{85}$$

while $\beta_1$ and $\beta_2$. Nevertheless, the exact equations for $K_1$ and $K_2$ as adapted from Supplementary Equation (73) are too complicated to be solved analytically. We therefore attempt to obtain approximate solutions for $K_1$ and $K_2$.

To this aim, we notice that variant *ohc-m* includes only one less possible combinations of $(\phi, \mathcal{C}^{(1)}, \mathcal{C}^{(2)})$ compared to variant *ohc-w*, *i.e.*, $(120^\circ, o, c)$,

$$(K_1)^{-1} + (K_2)^{-1} = (K_1^w)^{-1} + (K_2^w)^{-1} + \frac{\omega}{\Omega_{(1,2)}}\left((K_1^w)^{-1} + (K_2^w)^{-1} - (K_{\mathrm{d},o})^{-1} - (K_{\mathrm{d},c})^{-1}\right),$$
$$(K_1)^{-1}(K_2)^{-1} = (K_1^w)^{-1}(K_2^w)^{-1} + \frac{\omega}{\Omega_{(1,2)}}\left((K_1^w)^{-1}(K_2^w)^{-1} - (K_{\mathrm{d},o})^{-1}(K_{\mathrm{d},c})^{-1}\right),$$
$$\text{with} \quad \omega = K_{\mathrm{I}} K_{\beta,o} K_{\beta,c}, \tag{86}$$

where $K_{1,2}^w$ are the apparent binding affinities of $\beta_{1,2}$ in variant *ohc-w*, which are given in Equations (81) and (82); $\Omega_{(1,2)}$ is the normalization factor given in Supplementary Equation (73) for variant *ohc-m*. Note that Supplementary Equation (86) means that $(K_1)^{-1}$ and $(K_2)^{-1}$ are the two roots of the quadratic equation $x^2 - Bx + C = 0$, where $B = (K_1)^{-1} + (K_2)^{-1}$ and $C = (K_1)^{-1}(K_2)^{-1}$, *i.e.*,

$$(K_{1,2})^{-1} = \frac{1}{2}\left(B \pm \sqrt{\Delta}\right), \tag{87}$$

where $\Delta = B^2 - 4C$.

Further, we restrict our discussion to the case where $(K_{\mathrm{d},c})^{-1} \gg (K_{\mathrm{d},o})^{-1}, (K_{\mathrm{d},h})^{-1}$ and $(K_2^w)^{-1} \gg (K_1^w)^{-1}$. With these conditions, and because $B^2$ contain quadratic terms of $(K_2^w)^{-1}$ and $(K_{\mathrm{d},c})^{-1}$ as well as their cross-terms, it is implied that $B^2 \gg C$. Therefore,

$$\sqrt{\Delta} = \sqrt{B^2 - 4C} = B\sqrt{1 - \frac{4C}{B^2}} \approx B(1 - \frac{1}{2}\frac{4C}{B^2}) = B - \frac{2C}{B}. \tag{88}$$

Assuming that $K_1^{-1} < K_2^{-1}$, we have

$$K_2^{-1} = \frac{1}{2}\left(B + \sqrt{\Delta}\right) \approx B - \frac{C}{B} \approx (K_2^w)^{-1} - \frac{\omega}{\Omega_{(1,2)}}\left((K_{\mathrm{d},c})^{-1} - (K_2^w)^{-1}\right), \tag{89}$$

where $(K_{\mathrm{d},c})^{-1} - (K_2^w)^{-1} > 0$.

$K_2$ is expected to be close to $K_2^w$, thus $K_2$ is likely close to $K_3$. This explains why this variant *ohc-m* also failed to reproduce the measured ADP binding affinities [21] (Results).

### Supplementary Note 6 Hidden Markov analysis of the simulated single-molecule trajectories

As outlined in Methods, we employed hidden Markov analysis (HMA) to extract the dwells positions and kinetics (Fig. 4e-g) from the simulated kinetic Monte-Carlo trajectories (Results). This section provides a detailed explanation for the HMA algorithm.

#### Theoretical Description

The simulated trajectory $\boldsymbol{\phi}_{\mathrm{p}}$ is a vector of $L$ components $\phi_{\mathrm{p},l} = \phi_{\mathrm{p}}(t_l)$, $l = 1, 2, \ldots, L$, where $\{t_l\}$ are the discrete time points at which the angular position of the probe $\phi_{\mathrm{p}}(t)$ is recorded (Methods). Considering the rotational symmetry, the trajectory is projected onto $[0°, 360°)$ for further analysis.

Within HMA, we assume that the trajectory $\boldsymbol{\phi}_{\mathrm{p}}$ depends on the latent ("hidden") discrete time Markov process, where the system transitions among several dwells $\{\psi_k\}$ $(0 \leq \psi_k < 120°)$. Following the single-molecule studies [8, 15], we assume either one $(0 \leq \psi_1 < 120°)$ or two asymmetric dwells $(0 \leq \psi_1 < \psi_2 < 120°)$. According to the three-fold pseudo-symmetry of $F_1$-ATPase, we assume that each dwell contributes to three symmetric dwells $\varphi_{n,k} = \psi_k + 120° \times n$, $n = 1, 2, 3$. These dwells correspond to the "hidden Markov states" in general HMA.

The transitions between the dwells are quantified by the transition probability matrix $\mathbf{Q}$. The non-diagonal component $Q_{ji}\,(i \neq j)$ represents the transition probability of rotation from the $i^{\mathrm{th}}$ dwell to the $j^{\mathrm{th}}$ dwell, and the diagonal component $Q_{ii} = 1 - \sum_{j \neq i} Q_{ji}$ represents the probability to stay at the $i^{\mathrm{th}}$ dwell. We further assume that the system can only rotate between neighboring dwells. Accordingly, we define the transition probability matrix $\mathbf{Q}$ as below. If there is only one asymmetric dwell $\psi_1$ resulting in three symmetric dwells $(\psi_1, \psi_1 + 120°, \psi_1 + 240°)$, $\mathbf{Q}$ has the form:

$$\mathbf{Q} = \begin{bmatrix} 1 - P_+ - P_- & P_- & P_+ \\ P_+ & 1 - P_+ - P_- & P_- \\ P_- & P_+ & 1 - P_+ - P_- \end{bmatrix}, \tag{90}$$

where $P_+$ and $P_+$ represent the transition probabilities of counterclockwise and clockwise 120°-rotations, respectively. If there are two asymmetric dwells resulting in six symmetric dwells $(\psi_1, \psi_2, \psi_1 + 120°, \psi_2 + 120°, \psi_1 + 240°, \psi_2 + 240°)$, $\mathbf{Q}$ has the form:

$$\mathbf{Q} = \begin{bmatrix} 1-P_1 & P_{-\alpha} & 0 & 0 & 0 & P_{+\beta} \\ P_{+\alpha} & 1-P_2 & P_{-\beta} & 0 & 0 & 0 \\ 0 & P_{+\beta} & 1-P_1 & P_{-\alpha} & 0 & 0 \\ 0 & 0 & P_{+\alpha} & 1-P_2 & P_{-\beta} & 0 \\ 0 & 0 & 0 & P_{+\beta} & 1-P_1 & P_{-\alpha} \\ P_{-\beta} & 0 & 0 & 0 & P_{+\alpha} & 1-P_2 \end{bmatrix}, \tag{91}$$

with $P_1 = P_{+\alpha} + P_{-\beta}$, and $P_2 = P_{-\alpha} + P_{+\beta}$. Here, $\alpha$ and $\beta$ represent the two substeps within every 120°-step, $\alpha = \psi_2 - \psi_1$, $\beta = 120° - \alpha$; $P_{\pm\alpha}$ and $P_{\pm\beta}$ represent the transition probabilities of counterclockwise and clockwise $\alpha$-rotations and $\beta$-rotations, respectively.

Given that the system is in the $i^{\text{th}}$ dwell of angle $\varphi_{n,k}$, there is a certain probability $f_{\mathrm{E},i}(\phi_{\mathrm{p}})$ to observe the probe at angular position $\phi_{\mathrm{p}}$, known as the emission probability. Given the rotational symmetry, $f_{\mathrm{E},i}(\phi_{\mathrm{p}})$ follows the von Mises distribution,

$$f_{\mathrm{E},i}(\phi_{\mathrm{p}}) = \frac{\exp\left(\kappa_k \cos(\phi_{\mathrm{p}} - \varphi_{n,k})\right)}{2\pi I_0(\kappa_k)}, \tag{92}$$

where $I_0(\kappa_k)$ represents the modified Bessel function of the first kind of zero order; $\kappa_k$ measures the concentration of the distribution (angular variances).

While the von Mises distribution rigorously describes the circular statistics in our theoretical derivation (Supplementary Equation (92)), exact evaluation of the modified Bessel functions during the iterative parameter search can be prone to numerical instability when estimating highly concentrated distributions. Given that the angular variances within the dwells of $F_1$-ATPase are typically small (<20°), the resulting probability densities are highly concentrated around the true dwell angle. In such cases, the von Mises distribution is mathematically equivalent to a wrapped normal distribution. Therefore, to ensure algorithmic robustness and prevent severe arithmetic overflow, the wrapped normal distribution was employed as a reliable approximation in our practical code implementation:

$$f_{\mathrm{E},i}(\phi_{\mathrm{p}}) = \sum_{h=-\infty}^{+\infty} \frac{1}{\sqrt{2\pi}\sigma_k} \exp\left(-\frac{(\phi_{\mathrm{p}} - \varphi_{n,k} + 360° \times h)^2}{2\sigma_k^2}\right). \tag{93}$$

In summary, the parameters in our HMA include $\{\psi_k\}$, $\{\kappa_k\}$ (or its Gaussian equivalent standard deviation $\{\sigma_k\}$), and the transition probabilities ($P_+$ and $P_-$ if assuming only one asymmetric dwell; $P_{+\alpha}$, $P_{-\alpha}$, $P_{+\beta}$ and $P_{-\beta}$ if assuming two asymmetric dwells).

**The Baum-Welch Algorithm**

To obtain the maximum likelihood estimation of these parameters given the trajectory $\boldsymbol{\phi}_\mathrm{p}$, we adapted the Baum-Welch (BW) algorithm [93] to our system considering the three-fold rotational symmetry.

The BW algorithm requires the evaluation of the forward and backward probabilities. The forward probability $A_{il}$ quantifies the conditional probability of the system to produce the given observations until the $l^{\mathrm{th}}$ time point $t_l$, given that the system is in the $i^{\mathrm{th}}$ dwell at this time point $t_l$. Initialized by $A_{i1} = f_{\mathrm{E},i}(\phi_{\mathrm{p},1})$, $A_{il}$ is evaluated recursively for the following time points,

$$A_{il} = f_{\mathrm{E},i}(\phi_{\mathrm{p},l}) \sum_j Q_{ij} A_{j,l-1}, \quad l = 2, 3, \ldots, L. \tag{94}$$

Similarly, the backward probability $B_{il}$ quantifies the conditional probability of the system to continue with the given observations after a time point $l$, given that the system is in dwell $i$ at this time point $l$. Initialized by $B_{iL} = 1$, $B_{il}$ is evaluated recursively for time points in the reverse sequence,

$$B_{il} = \sum_j Q_{ji} f_{\mathrm{E},j}(\phi_{\mathrm{p},l+1}) B_{j,l+1}, \quad l = L-1, L-2, \ldots, 1. \tag{95}$$

Starting from an initial guess of the parameters, each iteration of our adapted BW algorithm includes an **estimation** step (E-step), followed by a **maximization** step (M-step).

In the E-step, the forward and backward probabilities are evaluated according to Supplementary Equations (94),(95). To ensure numerical stability when evaluating these probabilities for long trajectories, an underflow-prevention mechanism via recursive dynamic-scaling (utilizing renormalization counters) is implemented. This mechanism addresses the inherent numerical challenge where the sequential multiplication of probabilities systematically drives the forward and backward values toward numerical zero. To prevent this, our algorithm monitors these values at each time step: whenever the summed probabilities fall below a predetermined threshold, all state probabilities are divided by a small scaling factor, and a corresponding renormalization counter is incremented. These log-scaled corrections are later systematically accumulated when computing the full trajectory likelihood, thereby ensuring the mathematical integrity and numerical stability of the complete iterative optimization.

In the consecutive M-step, the parameters are reestimated using the forward and backward probabilities. To reestimate the transition probabilities, first, the transition probability matrix is updated using the original BW algorithm:

$$\hat{Q}_{ji} = \frac{\sum_{l=1}^{L-1} A_{il} f_{\mathrm{E},j}(\phi_{\mathrm{p},l+1}) B_{j,l+1}}{\sum_{l=1}^{L-1} A_{il} B_{il}} Q_{ji}. \tag{96}$$

Next, the transition probabilities are reestimated by averaging the respective components of the updated transition probability matrix $\hat{\mathbf{Q}}$:

$$\hat{P}_{+} = \frac{1}{3}(\hat{Q}_{13} + \hat{Q}_{21} + \hat{Q}_{32}), \quad \hat{P}_{-} = \frac{1}{2}(\hat{Q}_{31} + \hat{Q}_{12} + \hat{Q}_{23}), \tag{97}$$

or

$$\begin{aligned} \hat{P}_{+\alpha} &= \frac{1}{3}(\hat{Q}_{21} + \hat{Q}_{43} + \hat{Q}_{65}), \quad \hat{P}_{-\alpha} = \frac{1}{3}(\hat{Q}_{12} + \hat{Q}_{34} + \hat{Q}_{56}), \\ \hat{P}_{+\beta} &= \frac{1}{3}(\hat{Q}_{16} + \hat{Q}_{32} + \hat{Q}_{54}), \quad \hat{P}_{-\beta} = \frac{1}{3}(\hat{Q}_{61} + \hat{Q}_{23} + \hat{Q}_{45}). \end{aligned} \tag{98}$$

Finally, to maintain the symmetry, $\hat{\mathbf{Q}}$ is reconstructed according to Supplementary Equation (96) or (98) using the reestimated transition probabilities.

To reestimate $\{\psi_k\}$ and $\{\kappa_k\}$ (or $\{\sigma_k\}$), first, the emission probabilities are reestimated in a non-parameterized and discretized form $\mathbf{E}$ as in the original BW algorithm. The $[0°, 360°)$ interval is partitioned into 180 intervals of 2°, the $m^{\text{th}}$ interval being $\mathcal{I}_m = \left[2° \times m, 2° \times (m+1)\right)$ $(m = 0, 1, \ldots, 179)$, whose midpoint is $\omega_m = 2° \times m + 1°$. The probability for the $i^{\text{th}}$ dwell to produce an observed angular position within the interval $\mathcal{I}_m$ is updated by

$$\hat{E}_{im} = \frac{\sum_{\{l \mid \phi_{\mathrm{p},l} \in \mathcal{I}_m\}} A_{il} B_{il}}{\sum_{l=1}^{L} A_{il} B_{il}}. \tag{99}$$

Then, $\{\psi_k\}$ and $\{\kappa_k\}$ (or $\{\sigma_k\}$) are updated by the maximum likelihood estimates of the distribution parameters. $\hat{\psi}_k$ is determined by

$$\hat{\psi}_k = \arccos\left(\frac{\overline{x}_k}{\overline{R}_k}\right). \tag{100}$$

Here, $\overline{x}_k$ and $\overline{R}_k$ are determined by

$$\overline{x}_k = \frac{\sum_i \sum_m \hat{E}_{im} \cos(\omega_m)}{\sum_i \sum_m \hat{E}_{im}}, \quad \overline{y}_k = \frac{\sum_i \sum_m \hat{E}_{im} \sin(\omega_m)}{\sum_i \sum_m \hat{E}_{im}}, \quad \overline{R}_k = \sqrt{\overline{x}_k^2 + \overline{y}_k^2}, \tag{101}$$

where the summation of $i$ is over all the three symmetric dwells associated with the asymmetric dwell $\psi_k$; the summation of $m$ goes from 0 to 179. $\{\hat{\psi}_k\}$ are then transformed to the range $[0°, 120°)$ according to rotational symmetry.

For a rigorous implementation of the von Mises distribution (Supplementary Equation (92)), $\hat{\kappa}_k$ can be determined exactly by the equation $I_1(\hat{\kappa}_k)/I_0(\hat{\kappa}_k) = \overline{R}_k$. An approximate solution of this equation is available [94]:

$$\hat{\kappa}_k = \frac{\overline{R}_k(2 - \overline{R}_k^2)}{1 - \overline{R}_k^2}. \tag{102}$$

However, as stated above, we exclusively employed the wrapped normal distribution (Supplementary Equation (93)) in our actual code implementation. Accordingly, the standard deviation $\sigma_k$ is reestimated by directly computing the angular variance via weighted angular differences.

#### The Viterbi Algorithm

After obtaining the maximum likelihood parameters, we used the Viterbi algorithm [90, 95] to infer the sequence of dwells that is most probable to produce the trajectory $\boldsymbol{\phi}_\mathrm{p}$. First, the maximal probability $V_{il}$ of the system to produce the given observations until the $l^\mathrm{th}$ time point, given that the system is in dwell $i$ at this time point, is calculated. Initialized by $V_{i1} = A_{i1} f_{\mathrm{E},i}(\phi_{\mathrm{p},1})$, $V_{il}$ is evaluated recursively for the following time points $l = 2, 3, \ldots, L$,

$$V_{il} = f_{\mathrm{E},i}(\phi_{\mathrm{p},l}) \max_j \{V_{j,l-1} Q_{ij}\}. \tag{103}$$

Then, the most probable sequence of dwells $(s_1, s_2, \ldots, s_L)$ is inferred by backtracking from the last time point. Initialized by $s_L = \arg\max_j \{V_{jL}\}$, $s_l$ is determined recursively for the time points in the reverse sequence $(l = L-1, L-2, \ldots, 1)$,

$$s_l = \arg\max_j \{V_{jl} Q_{s_{l+1},j}\}. \tag{104}$$

With the inferred sequence of dwells, the trajectory $\boldsymbol{\phi}_\mathrm{p}$ was partitioned into parts where the system remained in one dwell, and the durations of these parts were counted to obtain the distributions of the dwell lifetime as shown in Fig. 4e–g.

### Supplementary Note 7 Detailed explanation about the functional states predicted by candidate model variants

In Discussion, we used our model predicted ensembles of joint functional states of the three β-subunits (functional stages) within the 80°- and 120°-dwells (Table 4, Supplementary Fig. 9) to interpret the functional roles of recent cryoEM structures. Here, we provide a more detailed explanation of these model predictions, as visualized in Supplementary Fig. 10 for the four candidate model variants (panel a for the $ohc_1^*c_2$-$w$ variant; panel b for the $ohc_1^*c_2^*$-$m.1$, $ohc_1^*c_2$-$m.2$, and $ohc_1^*c_2^*$-$m.2$ variants). The middle panels of Supplementary Fig. 10a,b present the predicted populations of the major functional stages depending on [ATP] by the heights of the differently colored areas.

**Predicted 80°-dwell ensemble** For all the four variants, the 80°-dwell is most populated by an $hc_1c_2$ state (dark blue), coexisting with a lower-populated $oc_1c_2$ state (light blue). As summarized in Supplementary Fig. 9a, these predictions match the "2 closed + 1 non-closed" nomenclature observed in single-molecule FRET measurements [70] and the recent cryoEM structures [32, 65] (Supplementary Fig. 9c,e).

**Predicted 120°-dwell ensemble** For the 120°-dwell, the predictions vary between the $ohc_1^*c_2$-$w$ variant and the other three variants.

For the other three variants (Supplementary Fig. 10b), the 120°-dwell is most populated by an $ohc_1/ohc_2$ state (dark pink), coexisting with a lower-populated $ooc_1/ooc_2$ state (light pink), and several other states of even lower populations (individual population $< 0.1$; the total population of these states is represented by the white area). As summarized in Fig. 9b, these predictions match the observed "1 closed + 2 non-closed" nomenclature [32, 65, 70] (Supplementary Fig. 9d,f). Specifically, the single closed β-subunit ($\beta_3$) is catalytically active and of high affinity.

For the $ohc_1^*c_2$-$w$ variant (Supplementary Fig. 10a), the 120°-dwell is almost exclusively dominated by an $oc_2c_1$ state (dark pink). At first glance, this state appears to possess two "closed" β-subunits, potentially conflicting with the "one closed" consensus. Particularly, the $\beta_2$-subunit adopts a closed conformation $c_2$, rather than a half-closed conformation $h$ that would directly align with the cryoEM structures [32, 65] ($\beta_{HO}$, Supplementary Fig. 9d,f). However, a closer inspection of the functional role of the $c_2$ conformation within this specific variant resolves this apparent discrepancy. In the $ohc_1^*c_2$-$w$ variant, the $c_2$ conformation represents a catalytically inactive state of intermediate affinity, in contrast to the catalytically active, high-affinity, and energetically most stable $c_1$ conformation. Considering the conformational progression of the $\beta_2$-subunit within a catalytic cycle, the $\beta_2$-subunit completes ATP hydrolysis at

80°, converts to this inactive $c_2$ conformation at 120°, and further opens to dissociate product at 200°. Thus, functionally, this $c_2$ conformation of the $\beta_2$-subunit at 120° serves as an intermediate state that connects the high-affinity, catalytically active $c_1$ conformation and the partially open $h$ conformation that facilitates product dissociation. Consequently, this $c_2$ conformation can be seen as an alternative intermediate $h$-like state, and the predicted $oc_2c_1$ state becomes effectively an $ohc$-like state. This interpretation aligns the predictions of the $ohc_1^*c_2$-$w$ variant with the consensus "1 closed + 2 non-closed" nomenclature discussed in Discussion (Supplementary Fig. 9b), demonstrating that all the four candidate variants are fundamentally consistent with available experimental evidence despite their detailed mechanistic differences.
